# Benchmarking machine learning models for the analysis of genetic data using FRESA.CAD Binary Classification Benchmarking

**DOI:** 10.1101/733675

**Authors:** Javier de Velasco Oriol, Antonio Martinez-Torteya, Victor Trevino, Israel Alanis, Edgar E. Vallejo, Jose Gerardo Tamez-Pena

**Affiliations:** Escuela Nacional de Medicina y, Ciencias de la Salud, Tecnologico, de Monterrey, Ave. Morones, Prieto 3000, 64710 Monterrey, Nuevo Leon, Mexico; Universidad de Monterrey, Morones Prieto 4500 Pte, 66238 San Pedro Garza Garcia, s Nuevo Leon, Mexico

**Author notes:** Equal contributor (Javier de Velasco Oriol, Jose Gerardo Tamez-Pena). Deceased June 11,2019 (Edgar E. Vallejo).

**Keywords:** Machine learning, model selection, R, FRESA.CAD

## Abstract

**Background:** Machine learning models have proven to be useful tools for the analysis of genetic data. However, with the availability of a wide variety of such methods, model selection has become increasingly difficult, both from the human and computational perspective.

**Results:** We present the R package FRESA.CAD Binary Classification Benchmarking that performs systematic comparisons between a collection of representative machine learning methods for solving binary classification problems on genetic datasets.

**Conclusions:** FRESA.CAD Binary Benchmarking demonstrates to be a useful tool over a variety of binary classification problems comprising the analysis of genetic data showing both quantitative and qualitative advantages over similar packages.

## Background

There is an increasing interest in using Machine Learning (ML) models for addressing questions on the analysis of genetic data [1, 2]. This is evident from the growing collection of papers related to these studies available on publication repositories. In effect, the number of papers in the PubMed repository with the keywords “machine learning” has increased almost tenfold during the last ten years (see Figure 1).

**Figure 1.**
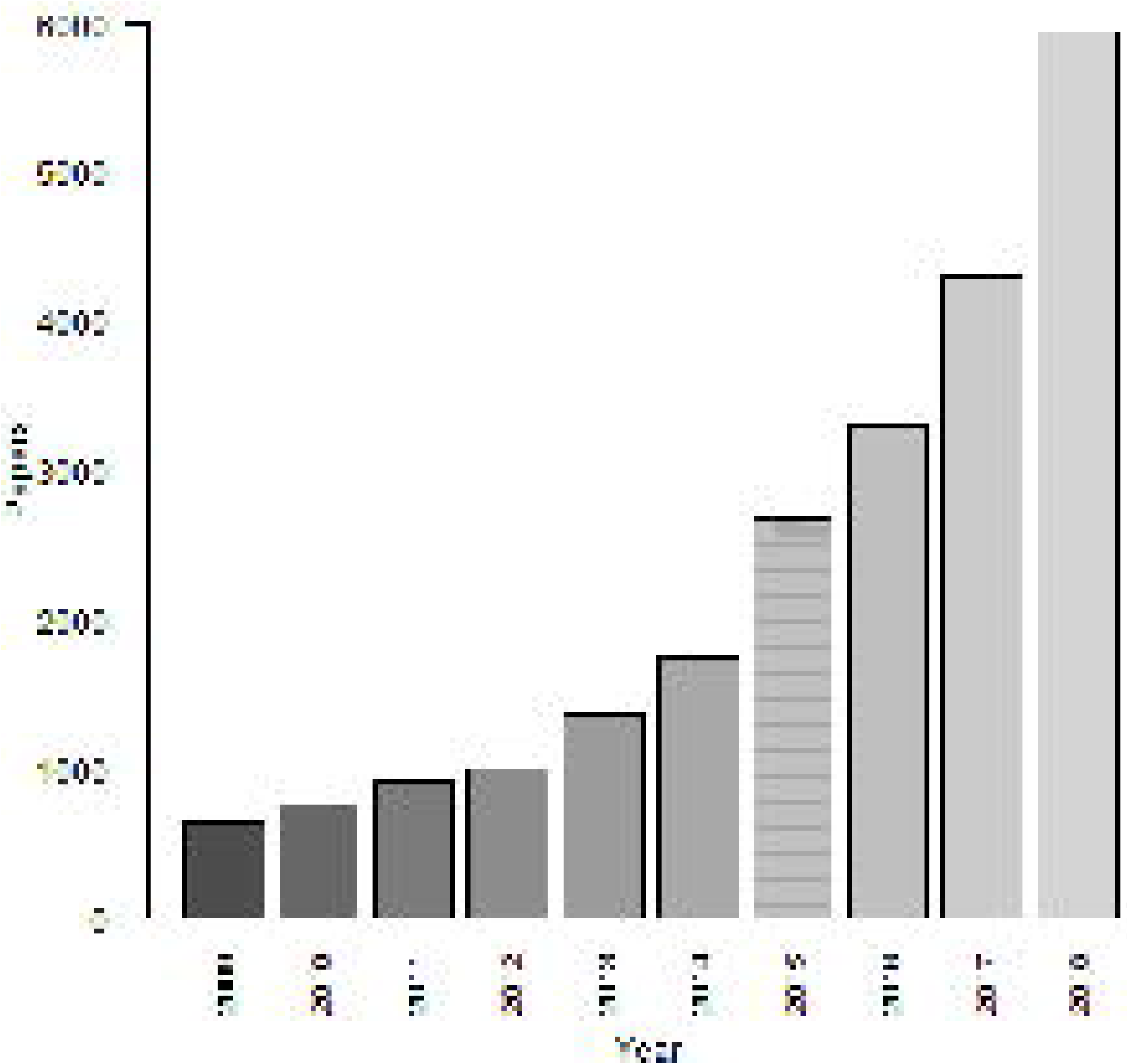
Collection of papers on the Pubmed repository. The paper counts were obtained using a query with the keywords “Machine learning”.

Similarly, an increasing collection of ML and statistical models are becoming available to address questions on the analysis of genetic data [3, 4]. Unfortunately, none of these ML models has shown to be superior at solving a variety of prediction problems involving genetic data. This limitation of ML models has been previously formulated as the “no free lunch theorem” [5]. The no free lunch theorem for classification models states that, averaged over all possible problems, every classification model has the same error rate when classifying previously unseen data; therefore, no ML model is always better than any other [6]. In the practice, this means that the goal of ML is not to seek the best learning model for all problems, but to find a model that performs best for a particular problem.

With the increasing availability of different ML models, systematic comparisons on the performance of a variety of these models is rapidly coming to be vital as part of model selection in the construction of predictive models applied to a variety of problems using genetic data. In effect, most studies reported in the literature [7, 8, 9] include validations of proposed methods consisting of comparisons of a variety of ML models. In addition, the majority of genetic variation and gene expression datasets consists of a large number of predictors and a small number of samples. The performance evaluation of classification models requires the splitting of data into training and validation sets. The holdout approach requires the random selection of a fraction of the data for training and the remaining fraction is used to evaluate the test performance of the method. Unfortunately, the validation set approach is typically not appropriate when using genetic data due to the high variance of the model. In effect, a single validation set would not be sufficient to assess the performance of the model accurately. Therefore, re-sampling techniques such as *k*-fold cross-validation and bootstrapping are often required for these problems [10].

Similarly, clinical diagnosis studies involving binary classification typically suffer from class imbalance: more control observations than cases, or vice-versa [11, 12]. In these problems, most binary classifiers often show biased classifications []. As a result, the performance of the classifier is not balanced with respect to sensitivity and specificity metrics. Moreover, traditional classification metrics such as accuracy can be misleading [13] as ML methods applied to a highly unbalanced dataset (i.e. 90% cases) might classify all the observations as cases, achieving a 90% accuracy but failing completely at the task of discriminating correctly between classes.

In most clinical diagnosis problems [14], the focus is on the construction of predictive models that show excellent performance on classification tasks. However, identifying the most informative predictors is often required in order to identify potential determinants of the outcome and for appropriate model selection. This is particularly evident for genetic diagnosis applications in which the principal aim of the experiments is often to identify associations of genetic loci with phenotypes, typically genome association studies.

In this article, we present FRESA.CAD Binary Classification Benchmarking extending the functionality of FRESA.CAD for conducting comparisons of a variety of representative ML models for rapid experimentation on the performance of prediction problems from genetic data. We demonstrate the functionality of the proposed package using a working problem consisting of the prediction of Type 2 diabetes using genetic variation data borrowed from OpenSNP [15]. We also provide summary test results on a set of molecular datasets and their vignettes are available at RPubs. We compared the performance and usability of FRESA.CAD Binary Classification Benchmarking using both quantitative and qualitative criteria with respect to those provided by WEKA and Caret, two popular packages that provide binary classification benchmarking capabilities [16] [17]. Our experiments using a variety of problems comprising the analysis of genetic data demonstrate that FRESA.CAD is a useful tool for conducting rapid experimentation addressing binary classification problems from genetic data, regarding performance, model selection, feature selection, presentation of results and usability.

## Implementation

Binary Classification Benchmarking was developed in R and it is a key component of the FRESA.CAD R package. FRESA.CAD evaluates the repeated holdout cross-validation (RHCV) [18] of binary classification algorithms or feature selection (FS) algorithms and returns a set of all the test results. Figure 2 shows the RHCV implemented in FRESA.CAD. The repeated test results are then combined into a single prediction per sample. The classifier performance can be visualized by ROC plots, described by confusion matrices and summarized by statistical performance metrics. The summary performance statistics are computed with the epiR package, which returns the 95% confidence intervals for all performance metrics [19]. The implemented RHCV also stores the train/test partitions, and the selected features at each train instance. The FRESA.CAD implementation of RHCV allows the comparison of different classifiers and feature selection algorithms under the same playground, hence removing variations due to differences in training/test sets and filter-filtering algorithms. In order to address class imbalance problems, we devised a procedure that creates balanced training folds using a combination of oversampling of the underrepresented class and undersampling of the overrepresented class. This procedure was inspired by the SMOTE algorithm [20].

**Figure 2.**
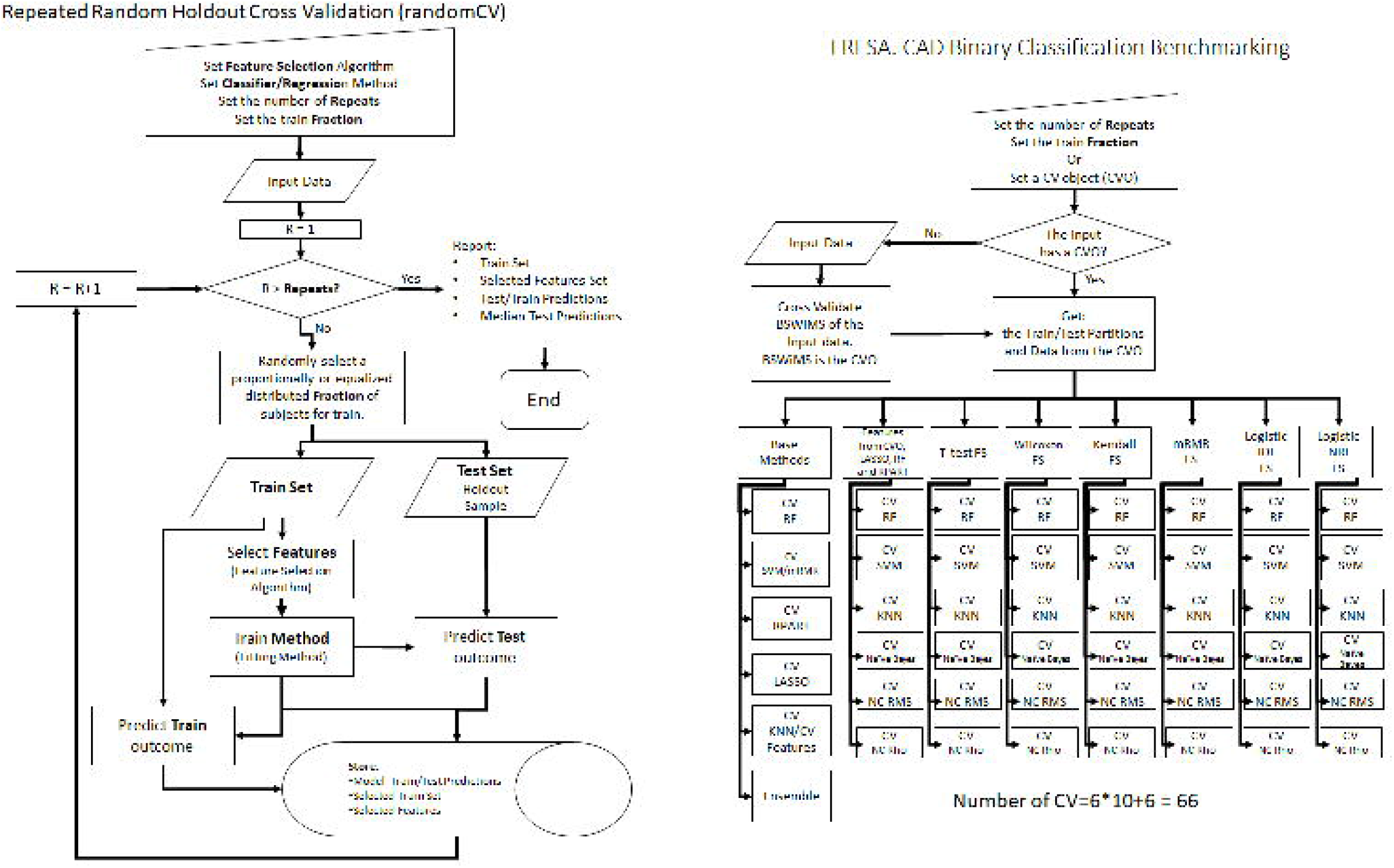
Repeatev Holdout Cross-Validation, and the Benchmark method.

The benchmark then performs the method comparison feature by takeing a user-supplied dataset and running a set of predetermined classifiers and feature selection algorithms on the data set. All the train/test results are stored and can easily be performance-ranked and visualized by FRESA.CAD provided functions. Furthermore, FRESA.CAD provides a set of functions that allows the simple exploration and comparison of the selected features of each filter method. The current implementation of FRESA.CAD benchmarking function evaluates the RHCV performance of the following algorithms: Bootstrap Stage-Wise Model Selection (BSWiMS) [21], Least Absolute Shrinkage and Selection Operator (LASSO) [22], Random Forest (RF) [23],Recursive Partitioning and Regression Trees (RPART)[24], K-Nearest Neighbors (KNN) with BSWiMS features, Support Vector Machine (SVM) [25] with minimum-Redundancy-Maximum-Relevance (mRMR)[26] feature selection filter, Naive Bayes (NB), and the Nearest Centroid (NC) algorithms. Finally, FRESA.CAD also performs the ensemble of all the above methods.

Each classification method is run using their default parameters or in combination with the following feature selection methods: BSWiMS, LASSO, RPART and RF; furthermore, the following filters are also mixed with the classification algorithms and the feature selection methods: integrated discrimination improvement (IDI), net reclassification improvement (NRI)[27], t student test, Wilcoxon test, Kendall correlation, and mRMR. These different variations give a total of 66 Cross-Validation instances. Figure 2 shows the workflow of the benchmark procedure. The default parameters of the Binary Benchmark function uses BSWiMS, but the user has the freedom to change the BSWiMS method for any data classifier.

The simplest way to visualize the results of the 66 Cross Validation instances executed by the FRESA.CAD Binary Benchmark is by using the provided FRESA.CAD plot function. The plot function compares the performance statistics and ranks them by comparing each CV instance’s 95% confidence interval (CI). The ranking method accumulates a positive score each time the lower CI of a performance metric is superior to the mean of the other methods and loses a point each time the mean is inferior to the top 95%CI of the other methods. The ranked data is visualized by bar plots that include the 95%CI. The plot function returns the accuracy, the sensitivity, the specificity, the balanced error rate and the receiver operating characteristic area under the curve (ROCAUC) with their corresponding 95% CI. The following code snippet shows a simple execution and visualization of the randomCV for the CV evaluation of the quadratic linear discriminant (MASS::qda) and benchmarks its performance to the methods evaluated by FRESA.CAD BinaryBenchmark functions.

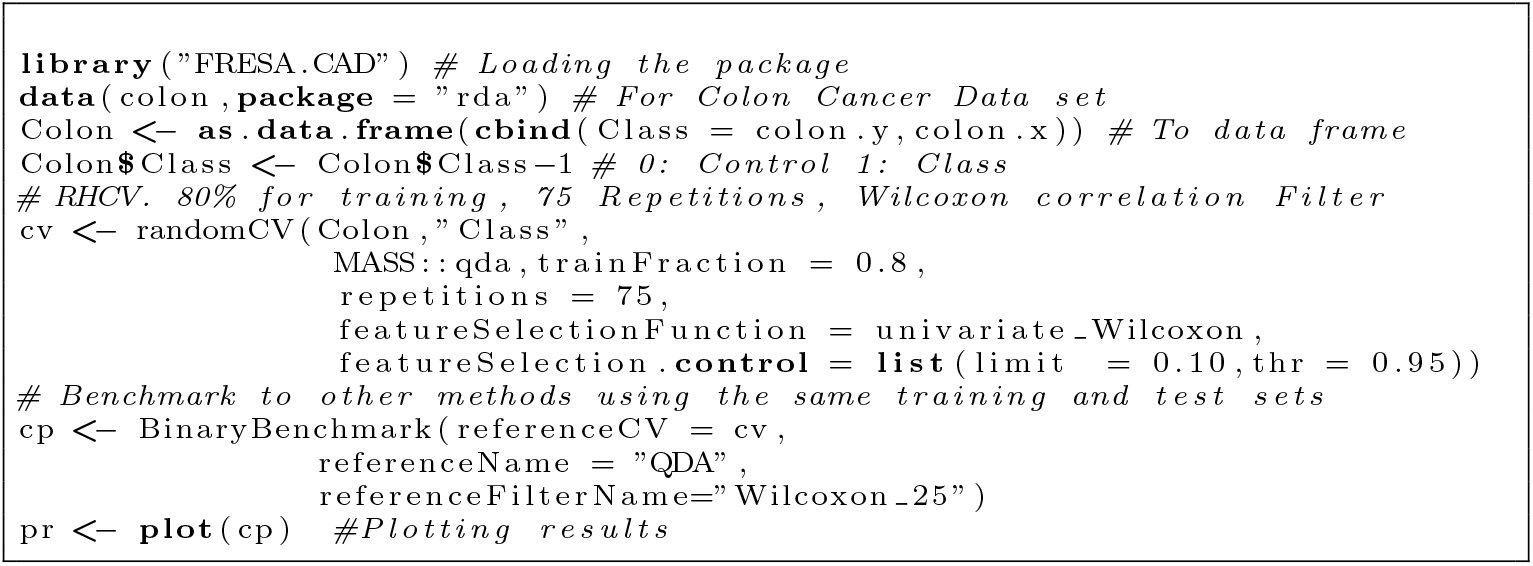

The result of the above code is provided in the supplementary material. (FRESA_DEMO.pdf) For demonstration purposes on SNP datasets, we will build a Type 2 diabetes data sets. We created the SNPs dataset using the Open-SNP(https://opensnp.org/) service. The OpenSNP query found 14 Type 2 subjects. We added 114 normal controls for a total of 128 individuals for our benchmarking study. The extracted data was stored into two different folders: one folder for case subjects and the second for control subjects. After that, we configured a Python and PLINK script to generate BED formatted files [28]. The BED files contained the phenotypes and the SNPs in a single dataset. The generated data set gives us the opportunity to study the behavior of classifiers and feature selection filters in a highly unbalanced dataset.

### Preprocessing the Genetic Data

The correct use of the FRESA.CAD benchmarking function on a specific dataset may require data conditioning and pre-processing, where the type of re-processing depends on the type of data. Here, we will show how to process the OpenSNP genotyping data. We followed the gene-data quality control methodologies [29]. The pre-process BED files were analyzed for marker call rate, sample call rate, minor allele frequency (MAF), and Linkage Disequilibrium (LD) [30]. Therefore, we removed the intrinsic factors in genetics that may cause methodological errors or inconsistencies between results. Sample call rate filtering threshold was set to 70%, marker call rate filtering threshold was set to 95%, MAF threshold was set to 1%, and LD was addressed by clumping [31]. Lastly, the final dataset, required by FRESA.CAD was generated using only the top 1000 SNPs ranked by their univariate p-values. This simplifies the task computationally for FRESA.CAD as well as eliminating those SNPs which have a very minor statistical correlation which could be due to random effects. The adjustments to the dataset with the Quality Control Pipeline can be analyzed in Table 1.

**Table 1.** Quality control procedures

| Processing Step | Cases | Controls | Features |
| --- | --- | --- | --- |
| Starting Dataset | 14 | 114 | 1,458,419 |
| SNP Quality Controls | 14 | 114 | 487,464 |
| LD-Based Clumping | 14 | 114 | 30,598 |
| Sample Filtering and SNP reduction | 11 | 110 | 1,000 |

### Running the Benchmark

Once the processed data was ready, the FRESA.CAD Benchmarking was run on the genetic dataset. Supplementary material: `Type2Diabetes.pdf` reports all the results of the experiment. In this section, we will present the main results. Figure 3 and 4 show the ROC Curve for the different classifiers. The AUC varied from 0.59 to 0.95. The Random Forest did not return good results. On the other hand, BSWiMS, LASSO and KNN models showed nice AUC performance. The Ensemble of the methods also shows great AUC performance.

**Figure 3.**
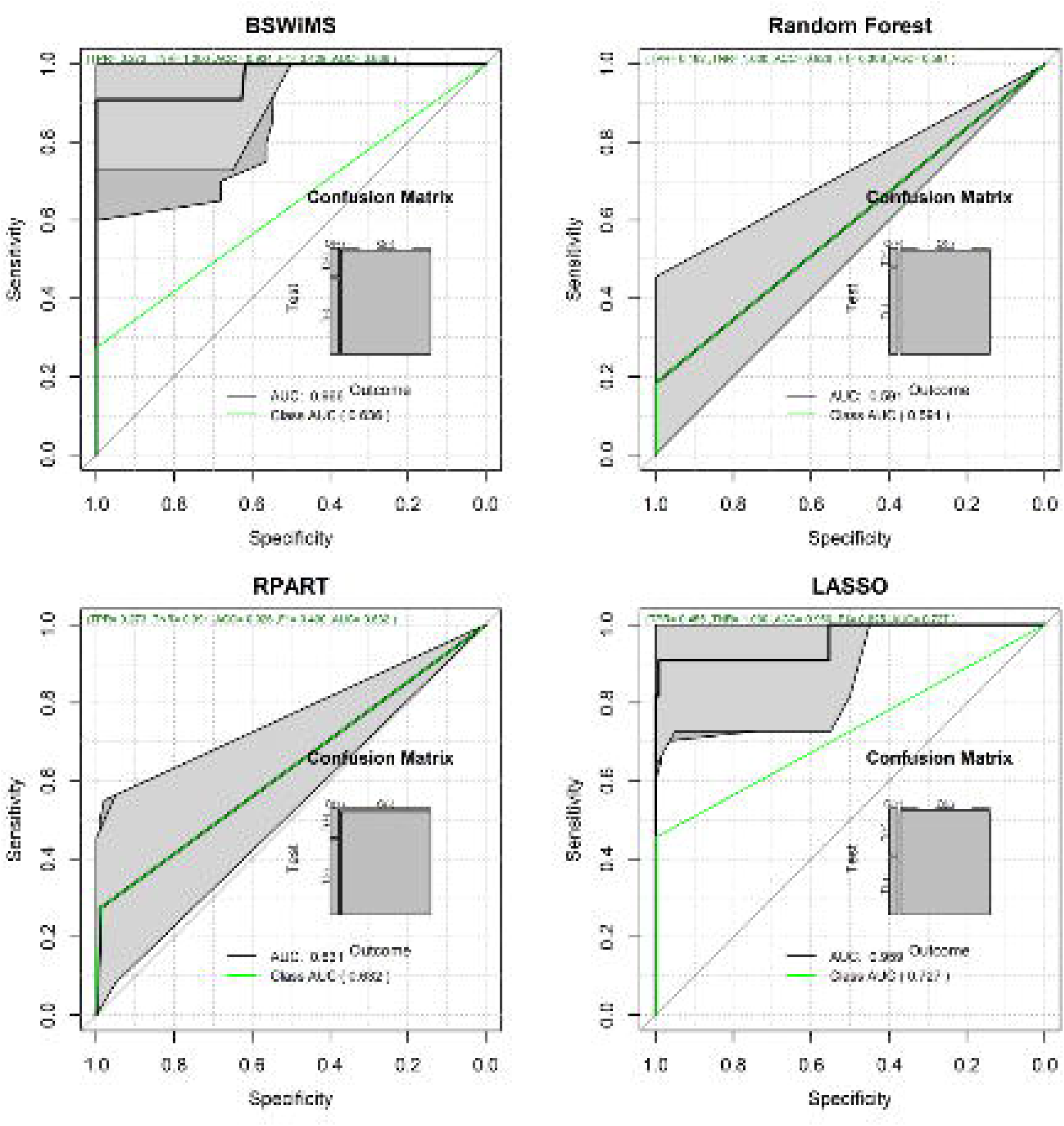
ROC Curves for the Benchmarking of the BSWiMS, Random Forest, RPART and LASSO Classifiers.

**Figure 4.**
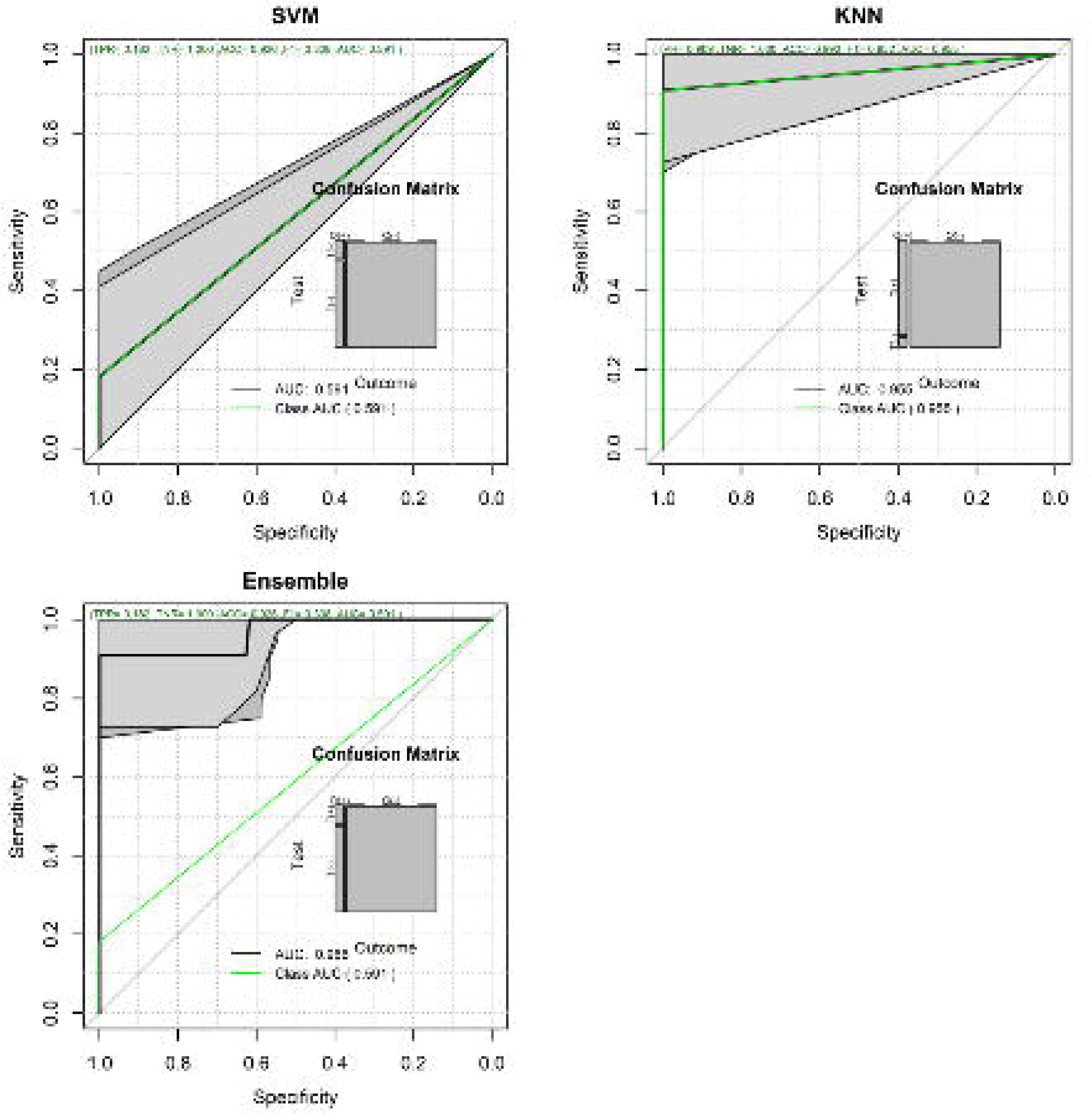
ROC Curves for the Benchmarking of the SVM, KNN Classifiers as well as the Ensemble.

### Analyzing the Results

Figure 5 shows the heat map of the prediction for every single classifier on every test subject and the relationship between classifiers. It can be observed that for cases the results of the classifiers differ. On one side we have RPART and the other extreme we have BSWiMS and Filtered RF. RF, SVM and Ensemble models have similar predictions, the figure also shows that the methods with more false positives were LASSO and RPART.

**Figure 5.**
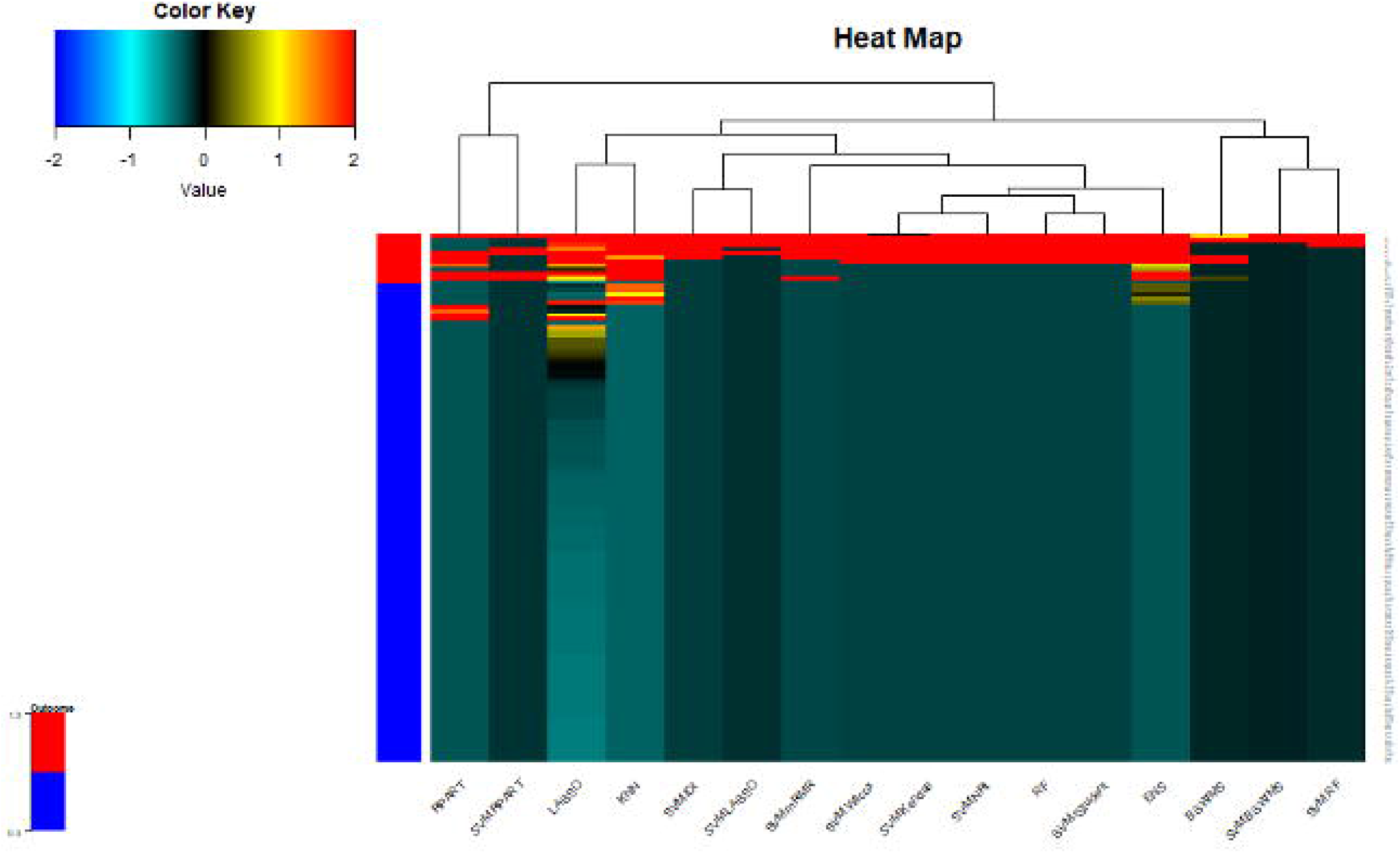
Heat map to compare the similarity between predictions.

Figures 6 shows three outputs of the `plot.BinaryBenchmark()` FRESA.CAD function. The Balanced Error Rate (BER) of the classifiers indicates that the KNN classifier was superior to the rest of the classifiers. The ROC AUC indicates that BSWiMS, LASSO, KNN, and Method Ensemble were superior to RF, RPART, and SVM. The ROC AUC of the combination of filters and classifiers shows that the nearest centroid classifiers (KNN, NC-Spearman, and NC-RSS) are superior to other methods when classifying SNPs data sets.

**Figure 6.**
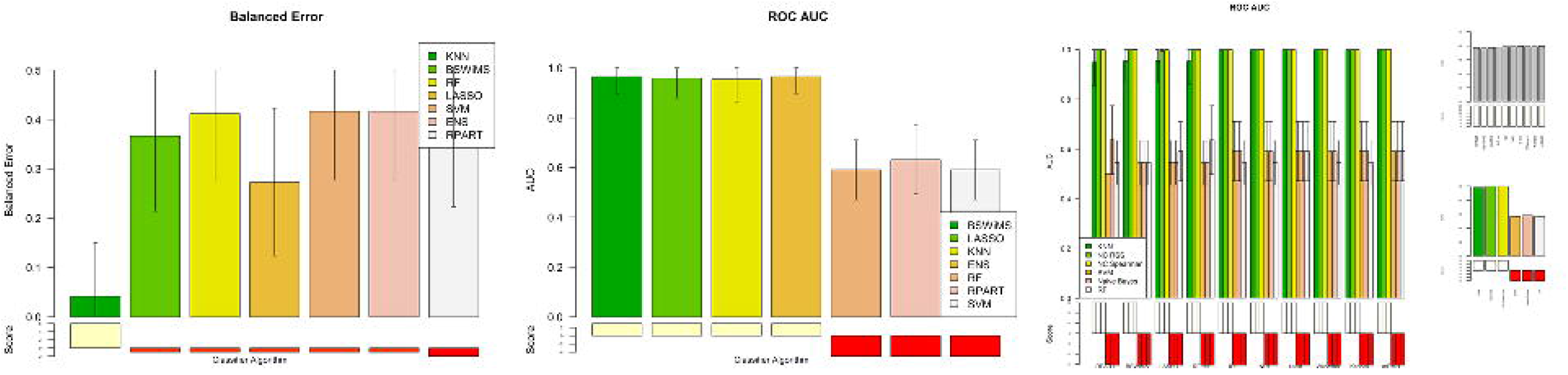
Balanced error of the classifiers for the Diabetes SNP dataset, ROC AUC of the classifiers for the Diabetes SNP dataset, ROC AUC for combinations of filters and classifiers.

### Meta-Analysis: Feature selection

FRESA.CAD provides a comprehensive analysis of the performance of the classifiers. If the main goal of the analysis is to explore relevant features, the object returned by the benchmarking stores the feature selection frequency of all filter methods, or model selection methods. Figure 7 shows the heat-map of the top SNPs and their association to cases and controls. Table 2 shows the univariate analysis provided by the FRESA.CAD univariate ranking function for each SNP. The table shows the univariate ROC AUC given by the SNP marker, the result of the Wilcoxon test and the corresponding p-value associated with type-2 diabetes. It is interesting that SNP rs2410284 was a top selected SNP even though, its univariate p-value was not significant. Furthermore, rs13136503 was also found to be associated with type-2 diabetes confirming previous reports [32].

**Figure 7.**
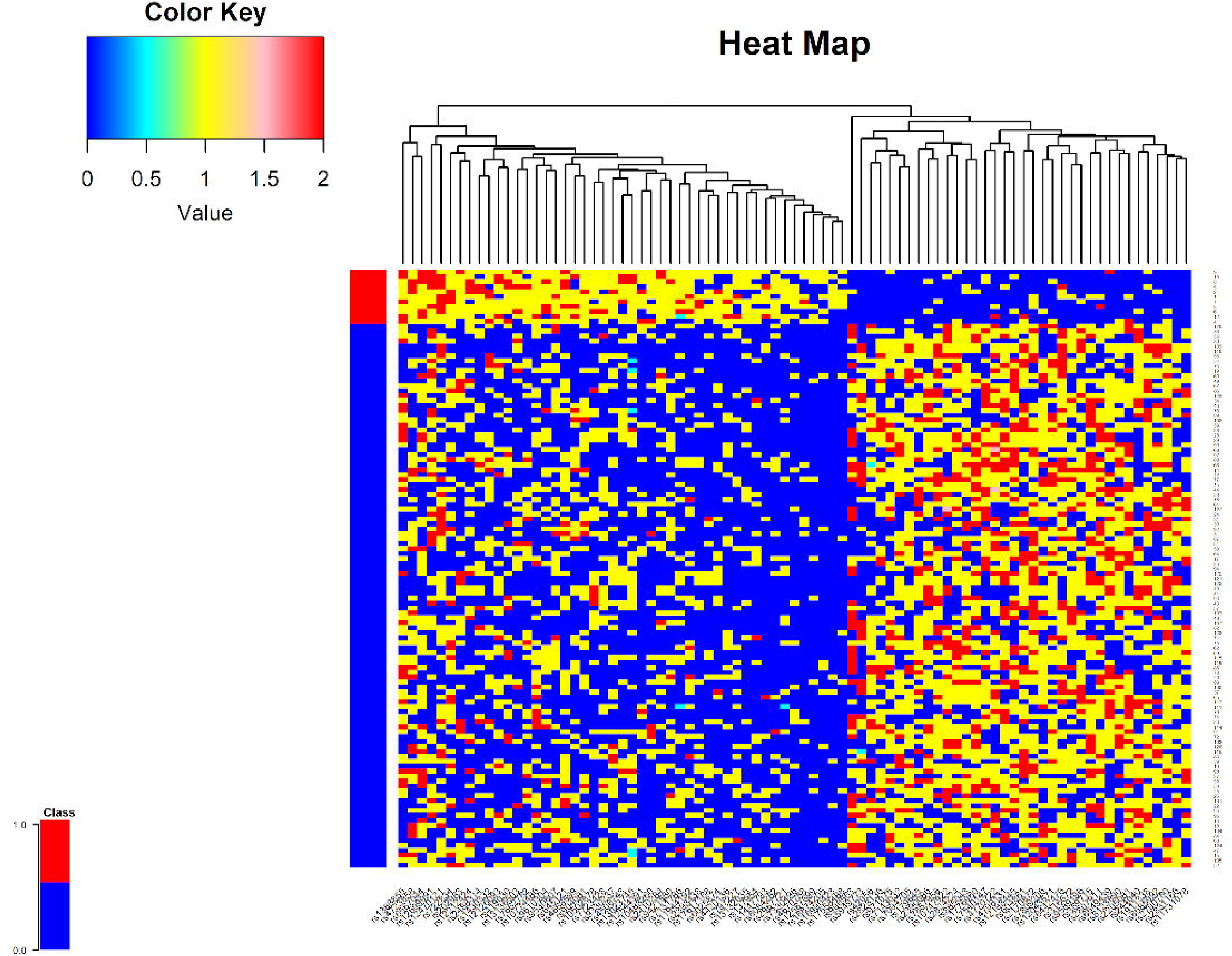
Values for the top features for the Diabetes Dataset.

**Table 2.** Top 20 SNPs

| SNP | ROCAUC | WILCOX | FREQ |
| --- | --- | --- | --- |
| rs2410284 | 0.85 | 0.1572 | 0.964 |
| rs7131786 | 0.8227 | 0.0479 | 0.746 |
| rs7857956 | 0.8182 | 0.0654 | 0.754 |
| rs6549439 | 0.8182 | 0.0688 | 0.732 |
| rs904081 | 0.8182 | 0.7038 | 0.898 |
| rs179645 | 0.8182 | 0.6765 | 0.904 |
| rs7143032 | 0.8182 | 0.6957 | 0.872 |
| rs10488260 | 0.8136 | 2.00E-04 | 0.68 |
| rs13132035 | 0.8091 | 8.00E-04 | 0.622 |
| rs13136503 | 0.8091 | 0.1641 | 0.675 |
| rs1776960 | 0.8091 | 0.1427 | 0.728 |
| rs13023745 | 0.8045 | 0.8428 | 0.868 |
| rs369715 | 0.8045 | 0.2106 | 0.655 |
| rs4420136 | 0.8045 | 0.0014 | 0.619 |
| rs10774486 | 0.8 | 0.898 | 0.829 |
| rs7094705 | 0.8 | 0.2682 | 0.613 |
| rs8123890 | 0.8 | 0.2962 | 0.631 |
| rs9589196 | 0.8 | 0.2742 | 0.645 |
| rs4798131 | 0.8 | 0.2751 | 0.682 |
| rs202983 | 0.7955 | 0.3385 | 0.615 |

### Comparing Benchmarking on Different Datasets

The binary classification benchmark was run on a set of different datasets: Leukemia [33], Colon [34], BRCA.REC [35], Lymphoma [36], ARCENE [37], Prostate [38], BRCA [39], Diabetes [15], and Depression [15]. In this section, we present the summary analysis of the results, while the supplementary material shows the detailed outputs and analysis of the results. Table 3 shows the characteristics of the nine data sets (before quality controls) used to evaluate the functionality of the binary benchmarking. As it can be seen multiple datasets are highly imbalanced, yet FRESA.CAD achieves good results thanks to the intrinsic methods implemented in the package. Figure 8 summaries the test-performance results of the main classifiers for each data set. We built Radar plots from the FRESA.CAD results by using a simple script as described in the demo document. The figure 8 shows those plots which rank each method on each one of the following performances metric: Sensitivity, Specificity, Accuracy, Balanced Error Rate, ROC AUC and CPU time. Figure 9 shows the average effect of filtering on the classification results. The plots report the following performance metrics: Accuracy, Sensitivity, Specificity, Balanced Error Rate, ROC AUC, plus the average size and Jaccard index. These figures clearly show that the best classifier is dataset dependent. While KNN overperformed all methods in the Type-2 diabetes data set, it was the worst classifier in the BRCA data set. We also see that RF is the best classifier in the BRCA recurrence study, and depression SNP; but, it did not well in the BRCA data set.

**Figure 8.**
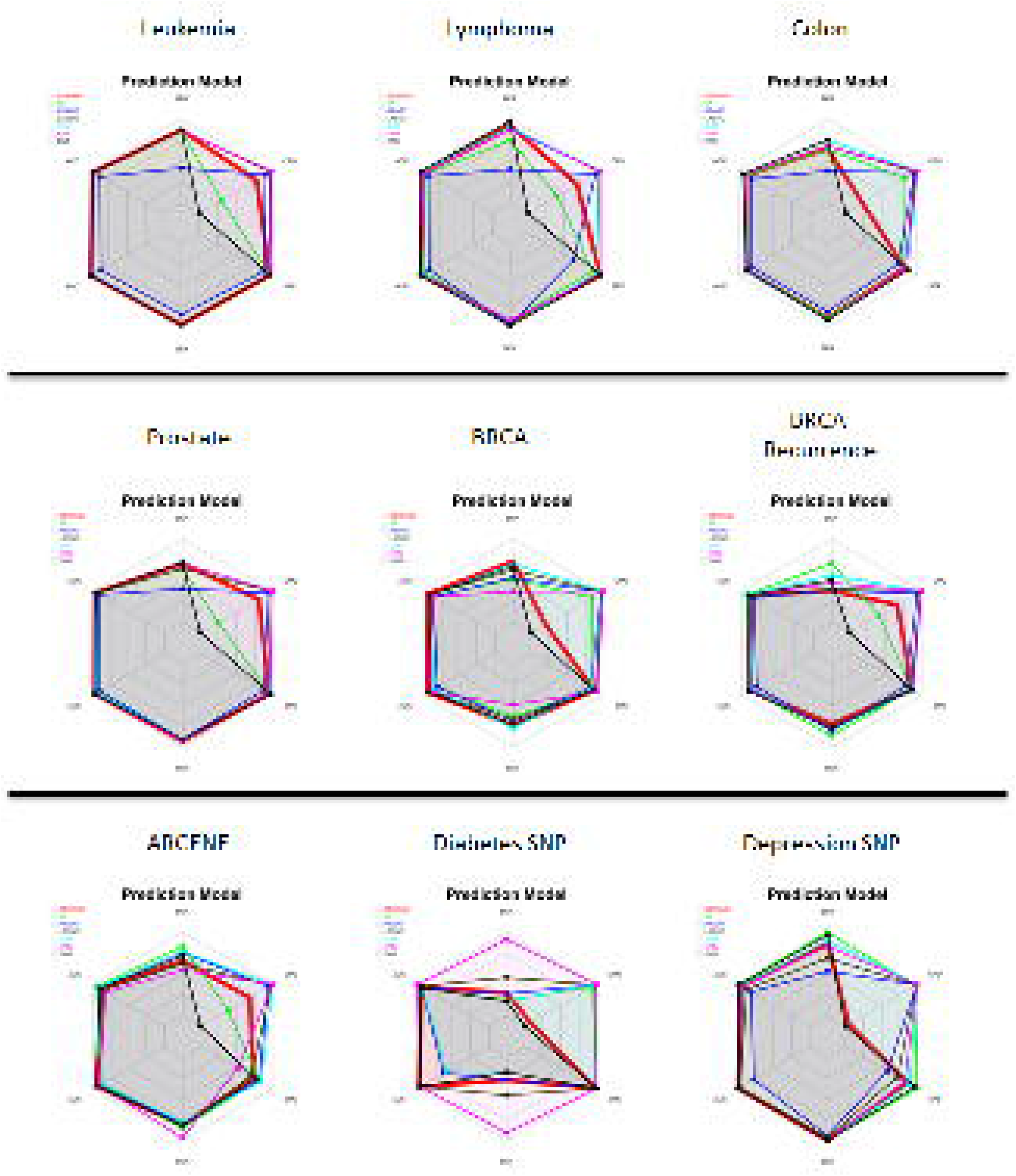
Radar plots of benchmarking classification methods on different data sets.

**Figure 9.**
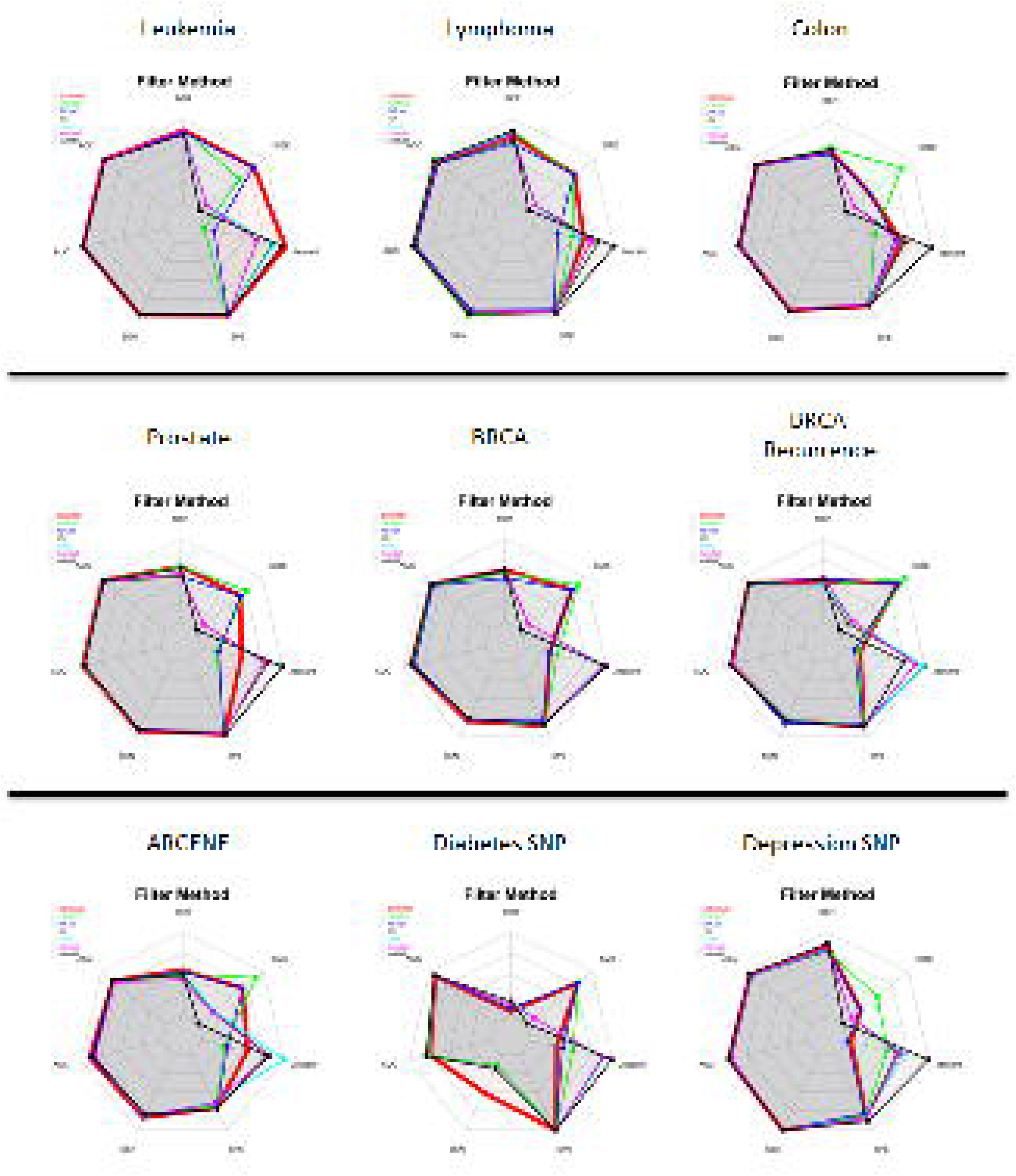
Radar plots showing the average effect of filters on the classification methods.

**Table 3.** Data set characteristics before QC

| Dataset | Cases | Controls | Number of Features |
| --- | --- | --- | --- |
| Leukemia | 25 | 47 | 7130 |
| Colon | 40 | 22 | 2000 |
| BRCA.REC | 80 | 308 | 10350 |
| Lymphoma | 58 | 19 | 7129 |
| ARCENE | 88 | 112 | 10000 |
| Prostate | 52 | 50 | 12600 |
| BRCA | 57 | 111 | 2905 |
| Diabetes | 11 | 110 | 7220 |
| Depression | 54 | 29 | 8300 |

## Comparison with other Benchmarking Software

The FRESA.CAD benchmark functionality and outputs were compared to other widely used machine-learning software packages: Caret [17] and Weka [16]. Both software packages can be configured to evaluate the cross-validation performance of classifiers. We set up Weka and Caret to cross-validate Random Forest, Support Vector Machines, Naive Bayes and K-Nearest Neighbors classifiers and we compared the results to the ones provided by FRESA.CAD. Furthermore, the Logistic Regression outputs of Caret and Weka were compared to BSWiMS logistic regression. For comparison purposes, we did the tests on two different datasets: Type-2 diabetes and BRCA. We attempted to setups all software parameters in a similar way, and the cross-validation was repeated 50 times and summary statistics were produced. The ROC AUC score was used to compare software outputs.

Figure 10 shows the ROC AUC returned by all software packages tested on Type-2 diabetes, while figure 11 shows the results of BRCA. These results show differences in CV performance, especially the results of SVM and RF. These can be expected due to the fact that RF and SVM have several hyper-parameters and FRESA.CAD was run with their default values. These values are different from other software implementations. Hence user must be careful in the interpretation of classifiers that depend on hyper-parameters. Table 4 shows the run times of the different software/classifiers. These times varied among software packages, but overall the amount of time required to run all 50 repetitions across different classifiers was similar. These run times were all performed in the same computer running on a i5-8400 CPU, on Windows 10 in Rstudio. In general, the results of FRESA.CAD were slightly superior compared to the other two toolkits, especially on the more unbalanced dataset of Type-2 Diabetes. The ease of use as well as the direct comparison graphs are two other areas where Weka and especially Caret were found lacking and where FRESA.CAD excelled at.

**Figure 10.**
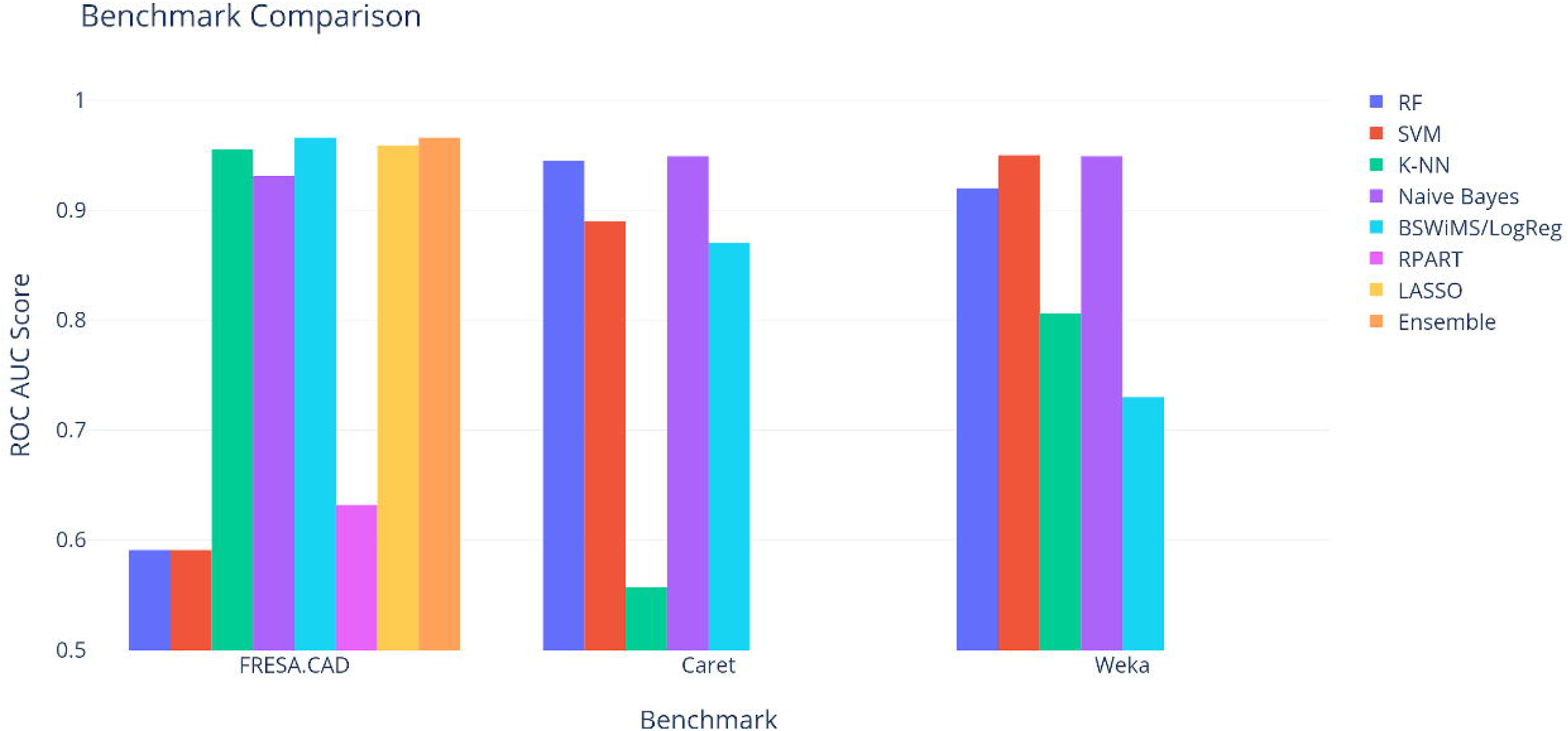
Benchmark comparisons for the Type-2 Diabetes dataset.

**Figure 11.**
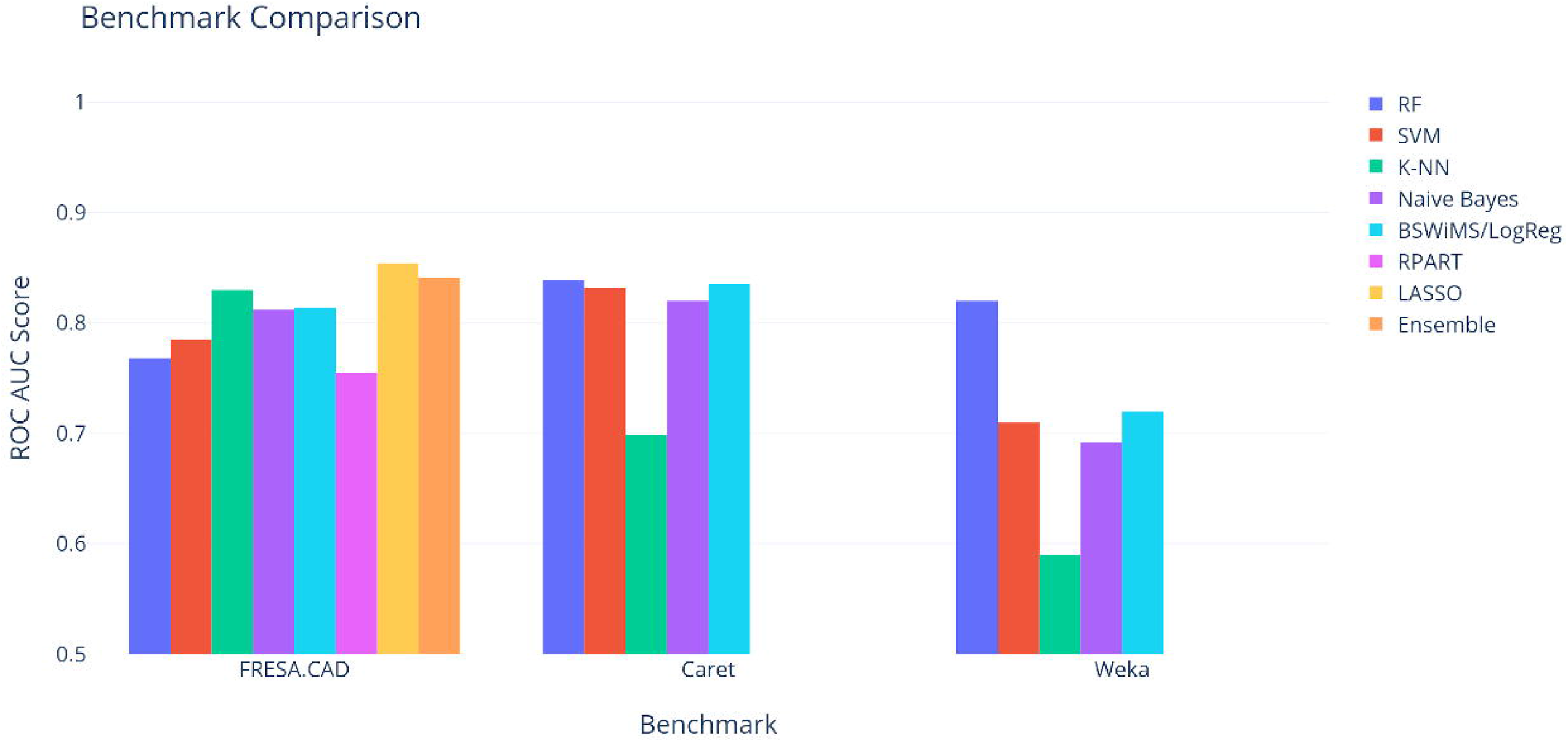
Benchmark comparisons for the BRCA dataset.

**Table 4.** Runtimes for one instance of each model (sec.)

| Benchmark | RF | SVM | K-NN | Naive Bayes | BSWiMS/ Log- Reg | RPART | LASSO | Ensemble | Total (s) |
| --- | --- | --- | --- | --- | --- | --- | --- | --- | --- |
| FRESA.CAD | 0.94 | 0.15 | 0.02 | 0.03 | 15.5 | 0.15 | 0.25 | 16.9 | 1,697 |
| Caret | 1.28 | 0.25 | 0.06 | 0.08 | 0.08 | / | / | / | 88 |
| Weka | 0.06 | 0.02 | 0.03 | 0.02 | 27.9 | / | / | / | 1,402 |

## Discussion

There is no panacea in ML: there is no superior model that performs best overall binary classification problems. Therefore, tools that allow for rapid experimentation on the performance of representative ML models intended to address binary classification problems are becoming increasingly useful for the analysis of genetic data. Genetic data typically consists of a large number of predictors and a much smaller number of samples. For example, genetic variation datasets often consist of millions of SNPs and thousands of samples. Similarly, gene expression datasets often consist of thousands of predictors and dozens of samples. These datasets are prone to high variance and overfitting when using ML models. Therefore, the systematic use of resampling methods such as k-fold cross-validation and bootstrapping are necessary. To evaluate these datasets, metrics that account for class imbalance like AUC and balanced error are typically required. FRESA.CAD Binary Classification Benchmarking uses these metrics and provides a collection of graphical results presented by ROC curves and heatmaps in order to simplify the interpretation of the results and compare the efficiency in balanced or imbalanced cases. FRESA.CAD Binary Classification Benchmarking carefully implements these methods for reducing the uncertainty associated with the analysis of genetic data and for the estimation of test errors of the adjusted ML models.

The FRESA.CAD Benchmark tool has some benefits over the other software available for benchmarking; it requires just one line to do the fitting and repetition with multiple models, as well as the comparison between them. On the other hand, Weka and Caret, offer greater flexibility, and more powerful tools to compare methods, but their usage is not straightforward. The main advantage FRESA.CAD has over the two other benchmarks is the feature selection system is directly implemented and how easily the resulting features can be obtained for analysis. The results also show directly the confidence intervals of each given metric to give. Furthermore, graphing is already included with the most useful types of graphics displayed with little code in a professional manner. This makes it so that FRESA.CAD is the best benchmarking tool out of the three for a researcher that wants to compare common methods easily on their dataset and wants to gain insight into the feature selection process required to further refine the investigation, as we were able to show in the sample type-2 diabetes experiment: benchmarking methods and meta-feature analysis identified key SNPs required to get good classification performance.

FRESA.CAD Binary Classification Benchmarking is designed for rapid experimentation with minimal effort, comparing multiple representative ML models and quickly exploring any type of binary classification problem. To compare multiple models just 3 steps are needed: process the dataset with the desired features and the binary labels to provide to the benchmark (if needed), run the FRESA.CAD Benchmarking and then analyze the different results and tables.

There are some limitations that have to be taken into account when using FRESA.CAD Benchmarking. First, exhaustive model optimization is not implemented due to computational considerations, second, SNP studies require preprocessing to mitigate biased results, but these preprocessing steps have to be evaluated carefully to avoid overfitting, hence independent validation may be required in all SNP analysis. However, FRESA.CAD binary benchmarking is sufficiently flexible and available as open source. Therefore, this R package can be extended to incorporate hyperparameters optimization for each of the considered ML models, and it may be extended to incorporate SNP preprocessing to mitigate false-positive discovery. We expect to incorporate this functionality in future versions of this package. Although, FRESA.CAD benchmarking capabilities can be used to evaluate continuous and ordinal regression models, it still lacks the ability to properly handle multiclass and multilabel classification problems. We are actively working on adding this capability because we believe these extensions would enhance the usability of the FRESA.CAD R package.

## Conclusions

We presented FRESA.CAD Binary Classification Benchmarking for supervised binary classification problems using genetic data. We presented a working problem related to the prediction of complex phenotypes from genetic variation data. We contrasted the functionality of FRESA.CAD benchmarking with those provided by WEKA and Caret packages. Overall, we believe that FRESA.CAD benchmarking is a promising alternative to allow for the rapid experimentation of ML models.

## Supporting information

Example for FRESA.CAD

Type 2 Diabetes Analysis

## Availability and requirements

Project name: FRESA.CAD Project home page: https://cran.r-project.org/web/packages/FRESA.CAD/index.html Operating system(s): Platform independent Programming language: R, C+—+ Other requirements: R 2.3 or higher Dependencies: Rcpp (≥ 0.10.0), stringr, miscTools, Hmisc, pROC. Suggest: nlme, rpart, gplots, RColorBrewer, class, cvTools, glmnet, randomForest, survival, e1071, MASS, naivebayes, mRMRe, epiR, DescTools, irr License: GNU GPL. Any restrictions to use by non-academics: none

## Declarations

**Ethics approval and consent to participate**

Not applicable.

## Consent for publication

Not applicable.

## Availability of data and material

The programs are available on CRAN. Datasets from examples are included as part of the R package.

## Authors’ contributions

JVO: Implementing and testing SNP and Genetic scripts, initial manuscript writing and careful review of the final version of the paper. AMT: Contributed to the implementation and optimization, FRESA.CAD manual writing, and careful review of final version of the paper. IA: Implementation of the C++ code, code optimization and parallelization of key functions. VT: Suggested many methods implemented in FRESA.CAD, careful review of final version of the code. EV: Critical review of ML algorithms, testing of FRESA.CAD benchmarking code, initial writing of manuscript, and critical review of the final version of the manuscript. JTP: FRESA.CAD software concept, implementation of the first versions of the code and Benchmark scripts, manuscript writing and review of the final version of the manuscript.

## Acknowledgments

We thank our colleagues from the Bioinformatics for Clinical Diagnosis research program, Escuela de Medicina y Ciencias de la Salud, Tecnologico de Monterrey for their valuable comments during the coding of the FRESA.CAD package and the preparation of this manuscript.

## Tables

### Additional Files

Additional file 1 — Supplementary material

Pdf file containing additional implementation details and additional figures and tables.

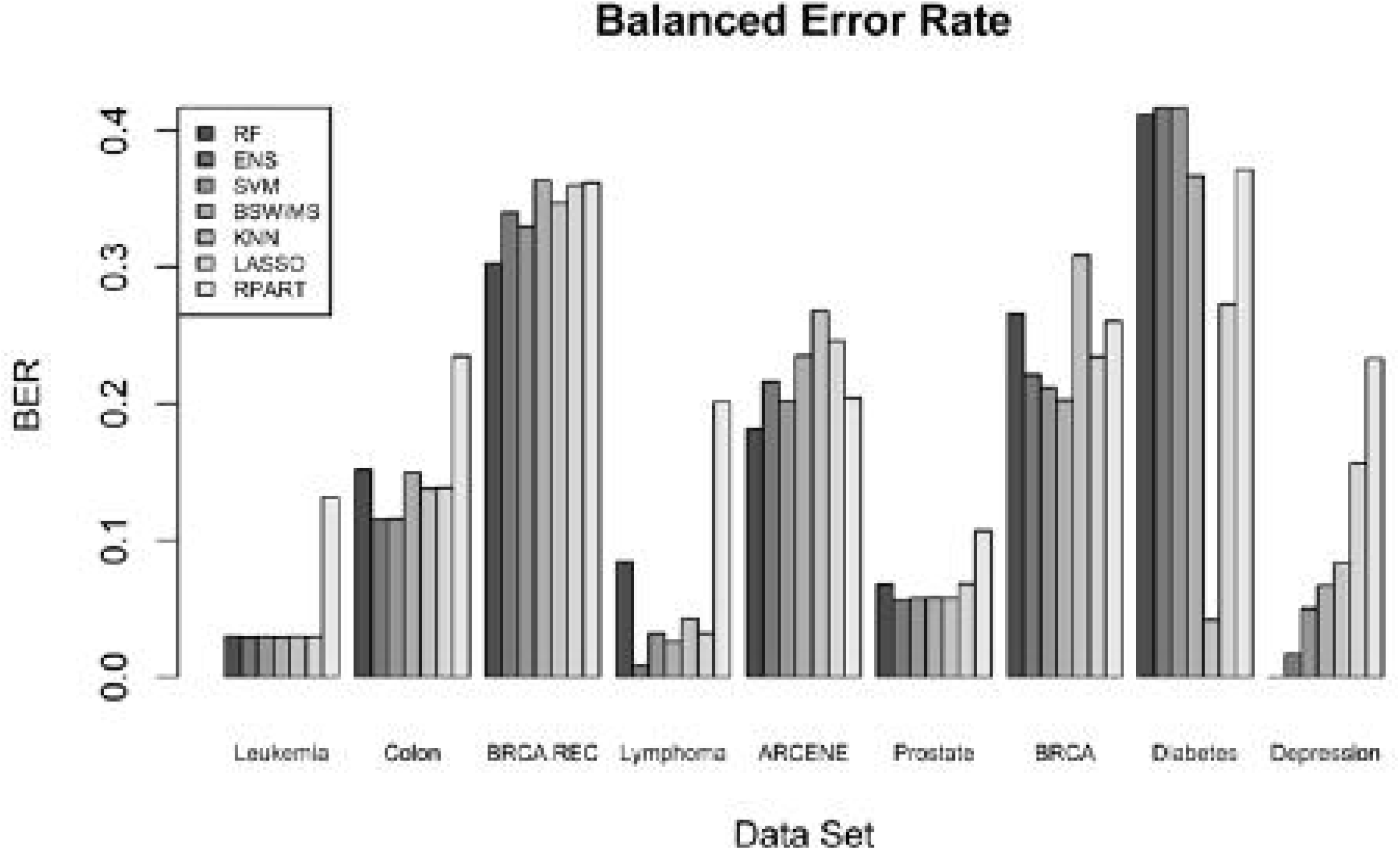

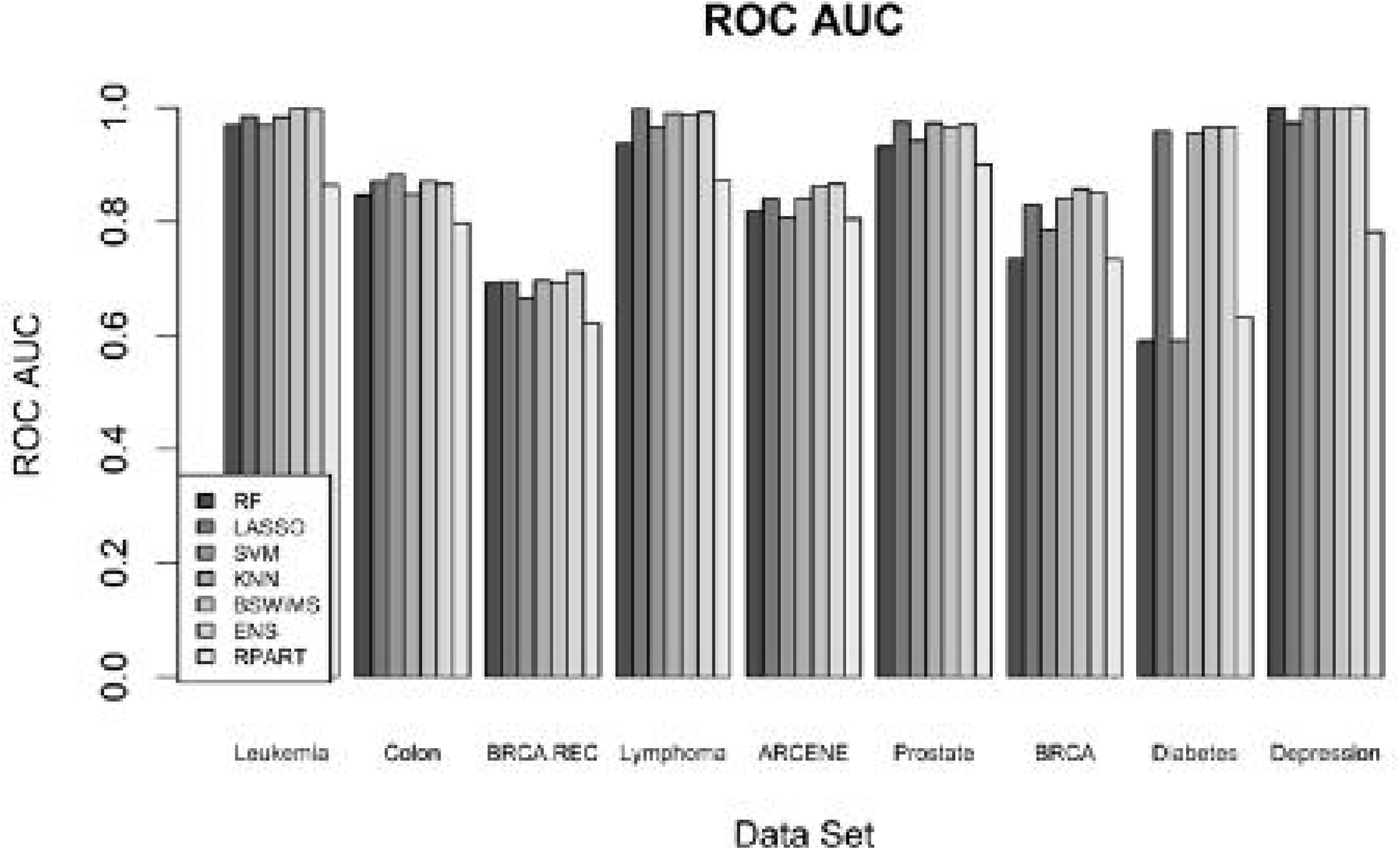

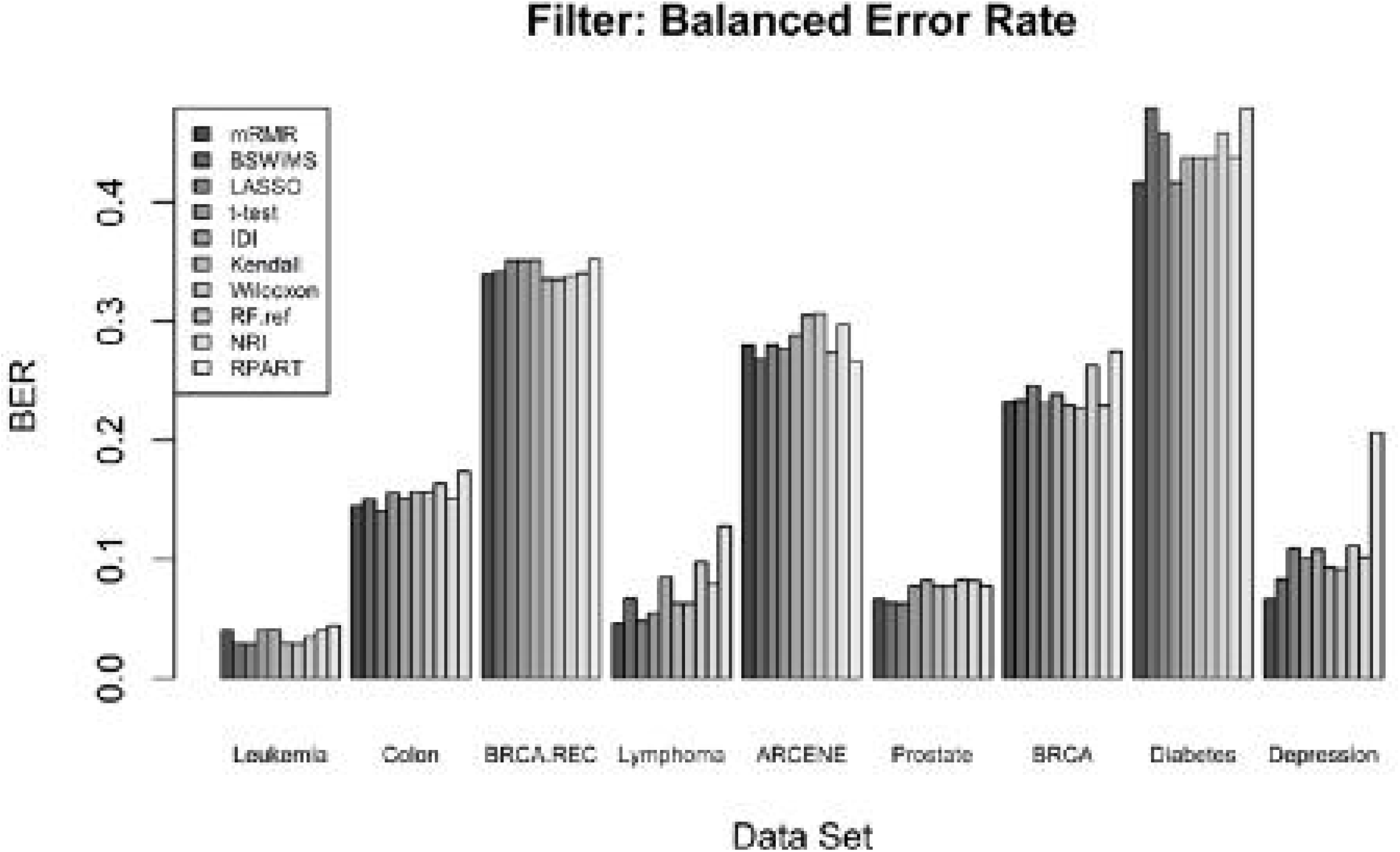

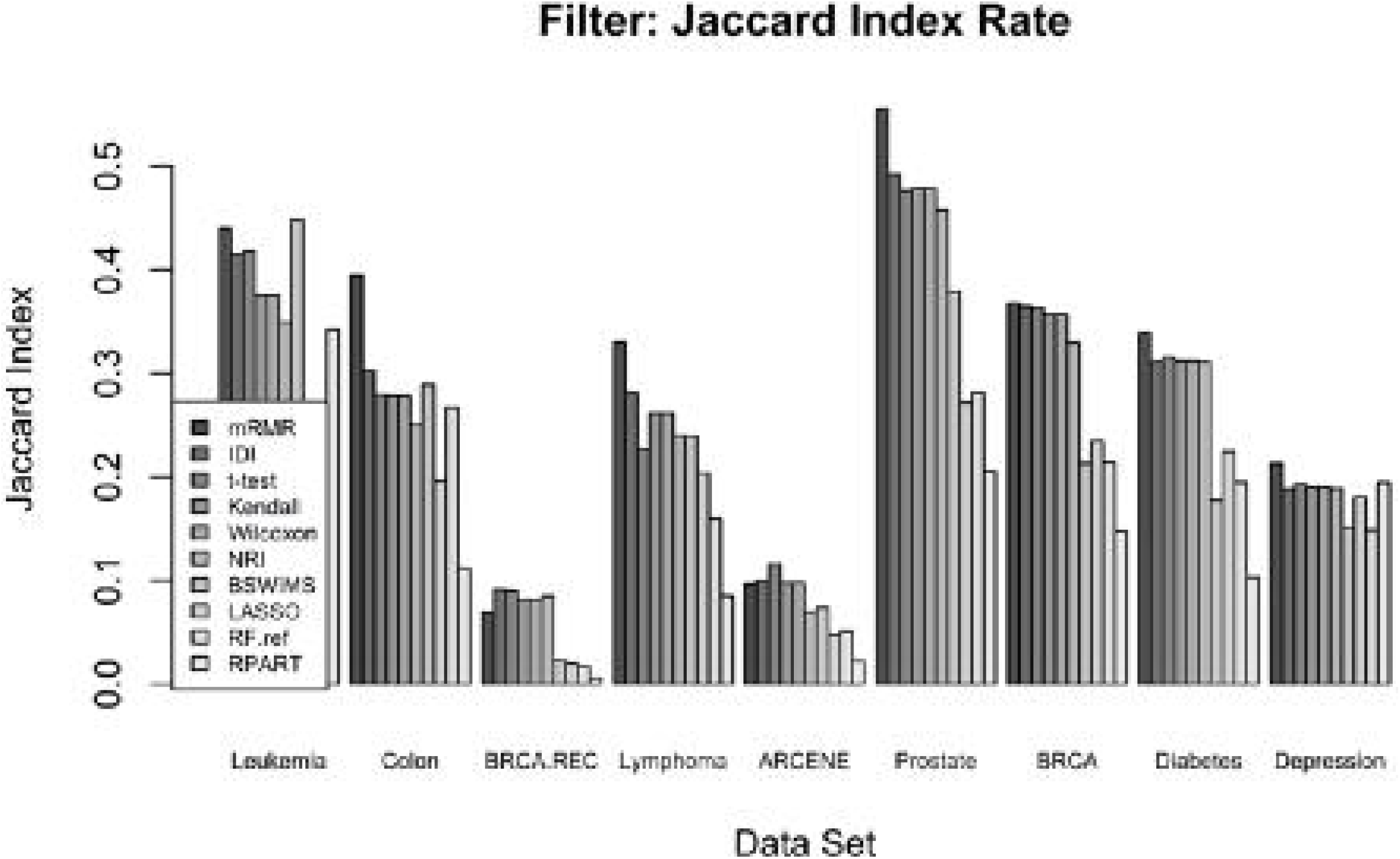

