## Supplementary material for "Benchmarking machine learning models for the analysis of genetic data using FRESA.CAD Binary Classification Benchmarking": Example for FRESA.CAD

### FRESA_DEMO

Jose Tamez-Pena

April 26, 2019

#Demo of CrossValidation and Binary Benchmarking

#### Data loading and preparation

In this demo, we will use the colon data set from the “rda” package

```
# Load the colon data form the rda package
data(colon,package = "rda")
#FRESA.CAD requires a data frame. One of the columns must be the class
Colon <- as.data.frame(cbind(Class = colon.y, colon.x))
#The class should be 0 for controls and 1 for cases
Colon$Class <- Colon$Class - 1
```

#### Cross-Validation of a QDA classifier

The colon cancer dataset has 62 observations and 2000 features. I will cross-validate (CV) a quadratic discriminant analysis (QDA) classifier that predicts the presence of cancer. Before estimating the QDA parameters a univariate filter based on the Wilcoxon-test will select the top 12 features (25% of 80% of the 62 samples, with a Pearson correlation lower than 0.95)

The CV will select 80% of the samples randomly for training, and the other 20% will be a holdout for validation.

The CV will be repeated 75 times; hence, on average each sample will have 15 estimations.

```
# Cross validate a QDA classifier using only the top ranked features
QDAcv <- randomCV(Colon,"Class",
                  MASS::qda,trainFraction = 0.8,
                  repetitions = 75,
                  featureSelectionFunction = univariate_Wilcoxon,
                  featureSelection.control = list(limit = 0.10,thr = 0.95))
```

#### Report the cross-validation performance

After CV, we can visuzlize the ROC and extract the test performance:

```
# Evaluating the test performance of the QDA classifier
par(mfrow = c(1,1),cex = 1.0);

QDAtestStats <- predictionStats_binary(QDAcv$medianTest,
                                        plotname = "QDA CV",cex = 0.9)
```

QDA CV

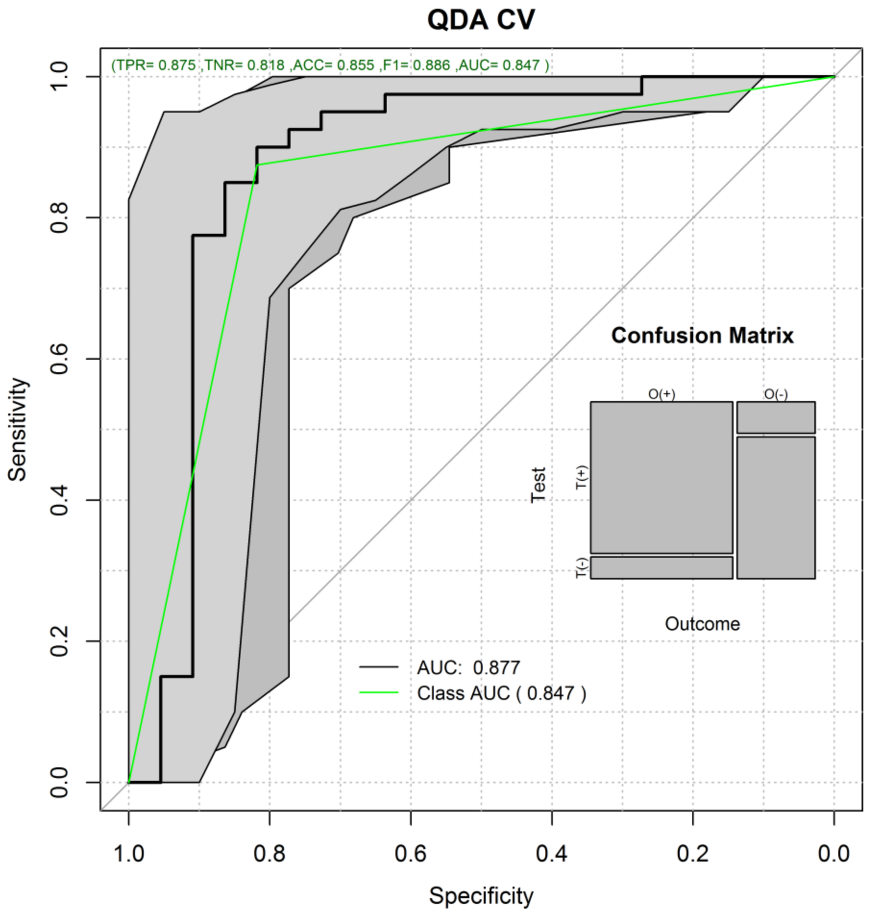

```
pander::pander(QDAtestStats$berror,caption = "Balanced Error",round = 3 )
```

| 50% | 2.5% | 97.5% |
|---|---|---|
| 0.148 | 0.061 | 0.257 |

```
pander::pander(QDAtestStats$ClassMetrics,caption = "Classification Performance",round = 3)
```

- **accci**:

| 50% | 2.5% | 97.5% |
|---|---|---|
| 0.855 | 0.758 | 0.935 |

- **senci**:

| 50% | 2.5% | 97.5% |
|---|---|---|
| 0.852 | 0.743 | 0.939 |

- **aucci**:

| 50% | 2.5% | 97.5% |
|---|---|---|
| 0.852 | 0.743 | 0.939 |

- **berci**:

| 50% | 2.5% | 97.5% |
|---|---|---|
| 0.148 | 0.061 | 0.257 |

- **preci**:

| 50% | 2.5% | 97.5% |
|---|---|---|
| 0.844 | 0.737 | 0.933 |

- **F1ci**:

| 50% | 2.5% | 97.5% |
|---|---|---|
| 0.846 | 0.738 | 0.931 |

#### Compare the QDA CV with other classifiers

The QDA CV can be compared to ohter common classifiers using the FRESA::BinaryBenchmark() function.

The same training and test sets will be used in all classifiers.

```
#comparing the cross validation to standard classifiers
par(mfrow = c(2,2),cex = 0.45);
ClassBenchmark <- BinaryBenchmark(referenceCV = QDAcv,
                          referenceName = "QDA",
                          referenceFilterName = "Wilcoxon_10")
```

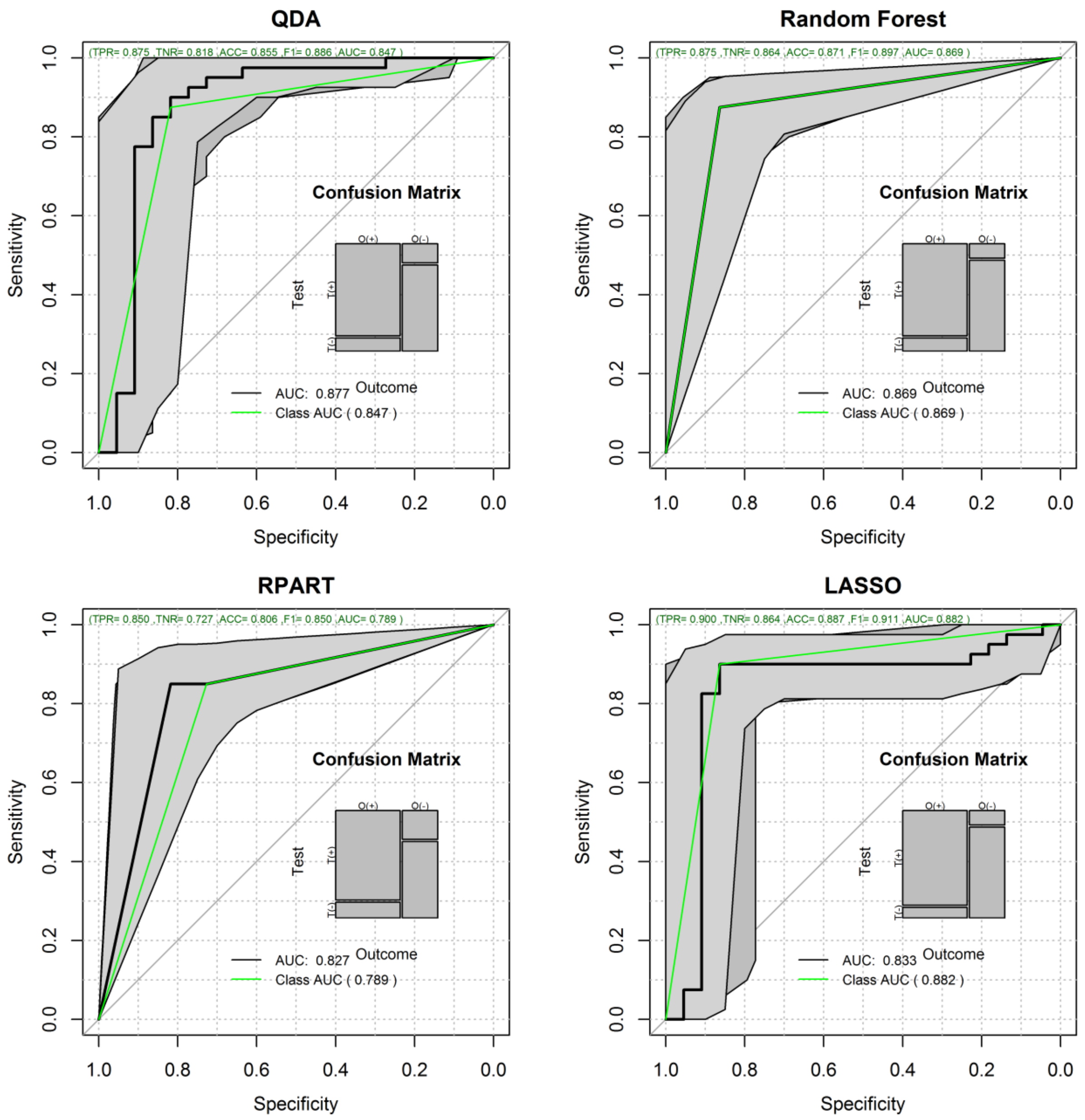

```
par(mfrow = c(1,1),cex = 1.0);
```

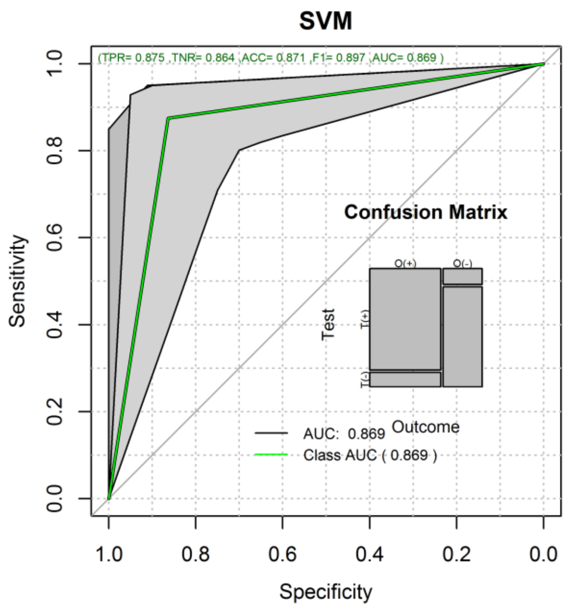

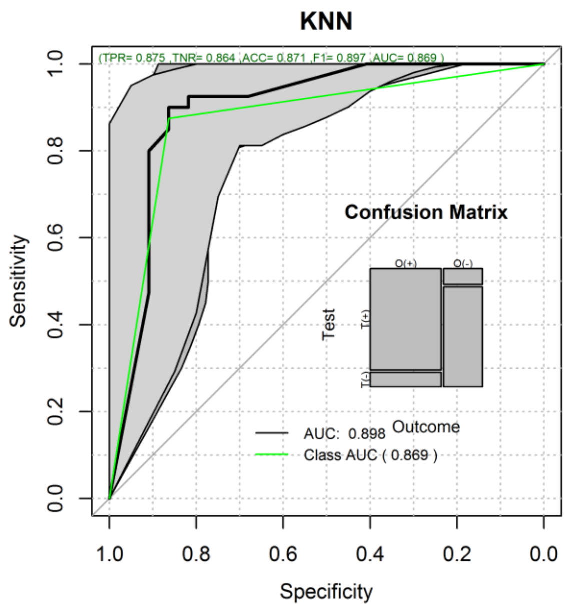

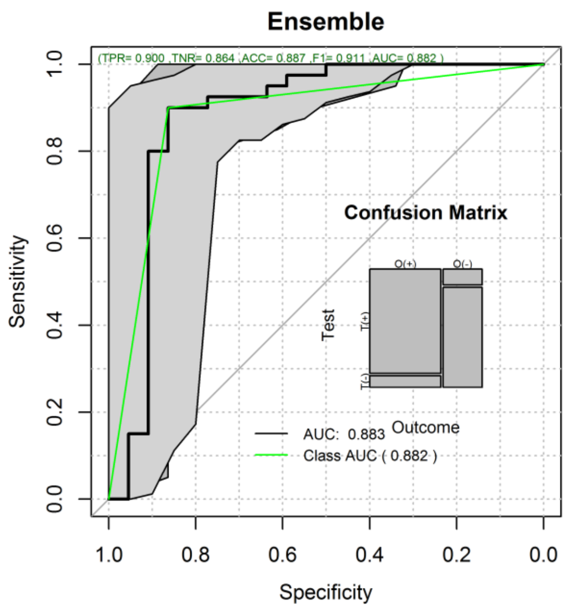

#### Reporting the results of the Benchmark procedure

Once done, we can compare CV test results using the plot() function. The plot function also generates summary tables of the CV results.

```
#ploting the results
par(mfrow = c(1,2),cex = 1.0)
prBenchmark <- plot(ClassBenchmark)
```

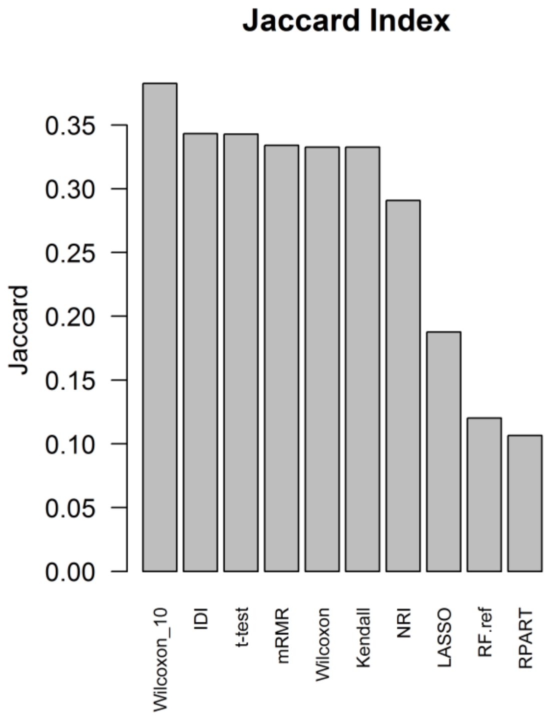

Jaccard Index
Jaccard
0.35
0.30
0.25
0.20
0.15
0.10
0.05
0.00
Wilcoxon_10
IDI
t-test
mRMR
Wilcoxon
Kendall
NRI
LASSO
RF.ref
RPART

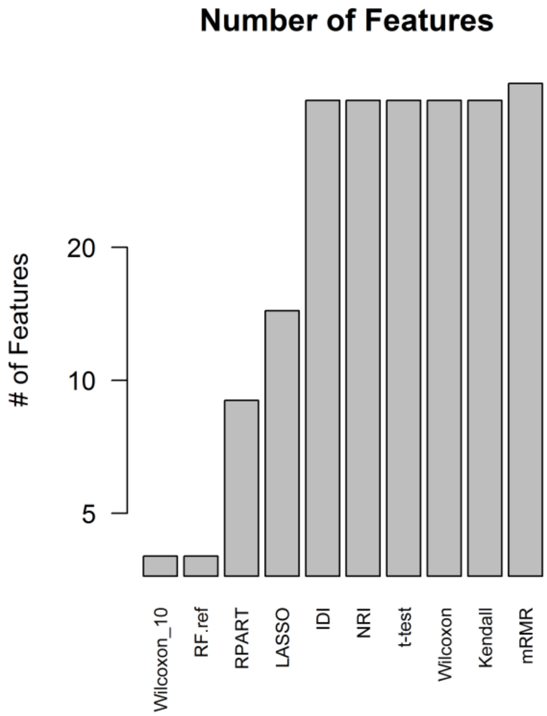

Number of Features
### of Features
20
10
5
Wilcoxon_10
RF.ref
RPART
LASSO
IDI
NRI
t-test
Wilcoxon
Kendall
mRMR

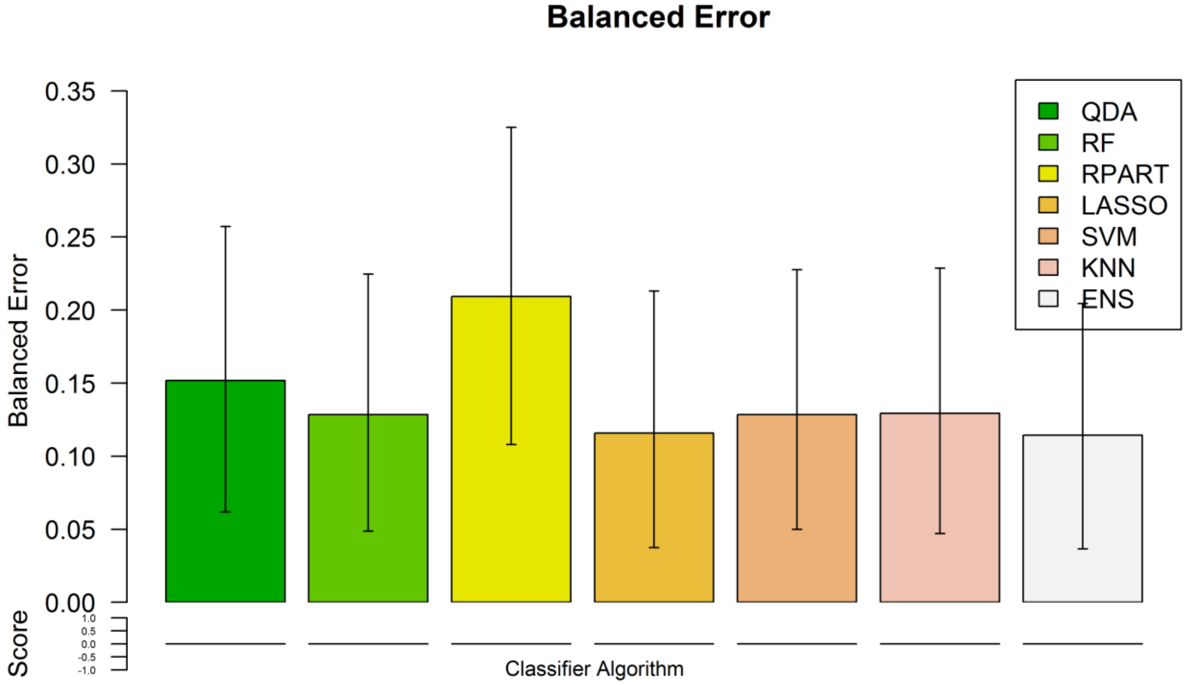

Balanced Error
Balanced Error
0.35
0.30
0.25
0.20
0.15
0.10
0.05
0.00
Score
1.0
0.5
0.0
-0.5
-1.0
Classifier Algorithm
QDA
RF
RPART
LASSO
SVM
KNN
ENS

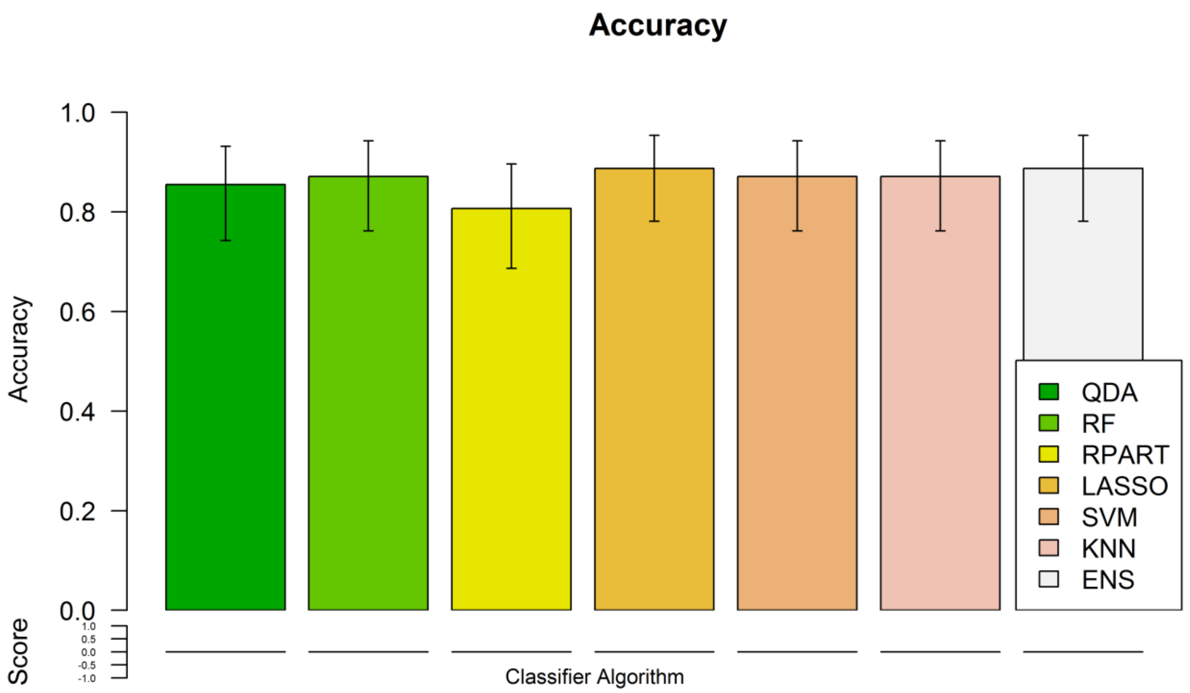
Accuracy
1.0
0.8
0.6
0.4
0.2
0.0
Accuracy
Score
1.0
0.5
0.0
-0.5
-1.0
Classifier Algorithm
QDA
RF
RPART
LASSO
SVM
KNN
ENS

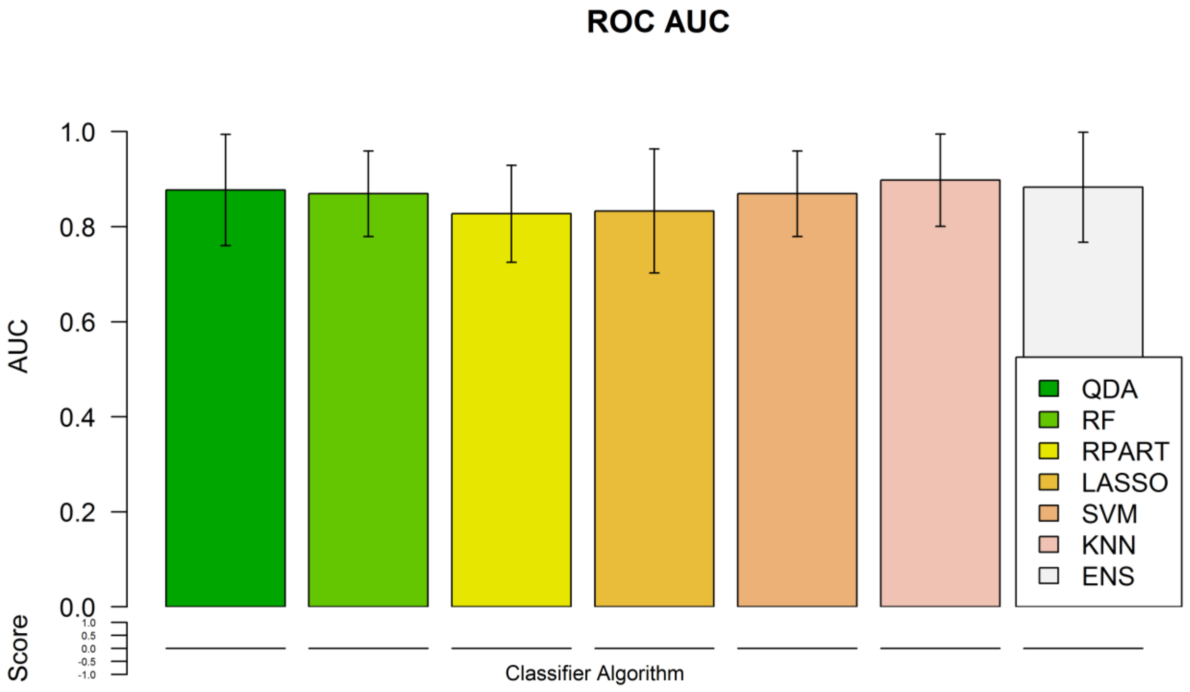
ROC AUC
1.0
0.8
0.6
0.4
0.2
0.0
AUC
Score
1.0
0.5
0.0
-0.5
-1.0
Classifier Algorithm
QDA
RF
RPART
LASSO
SVM
KNN
ENS

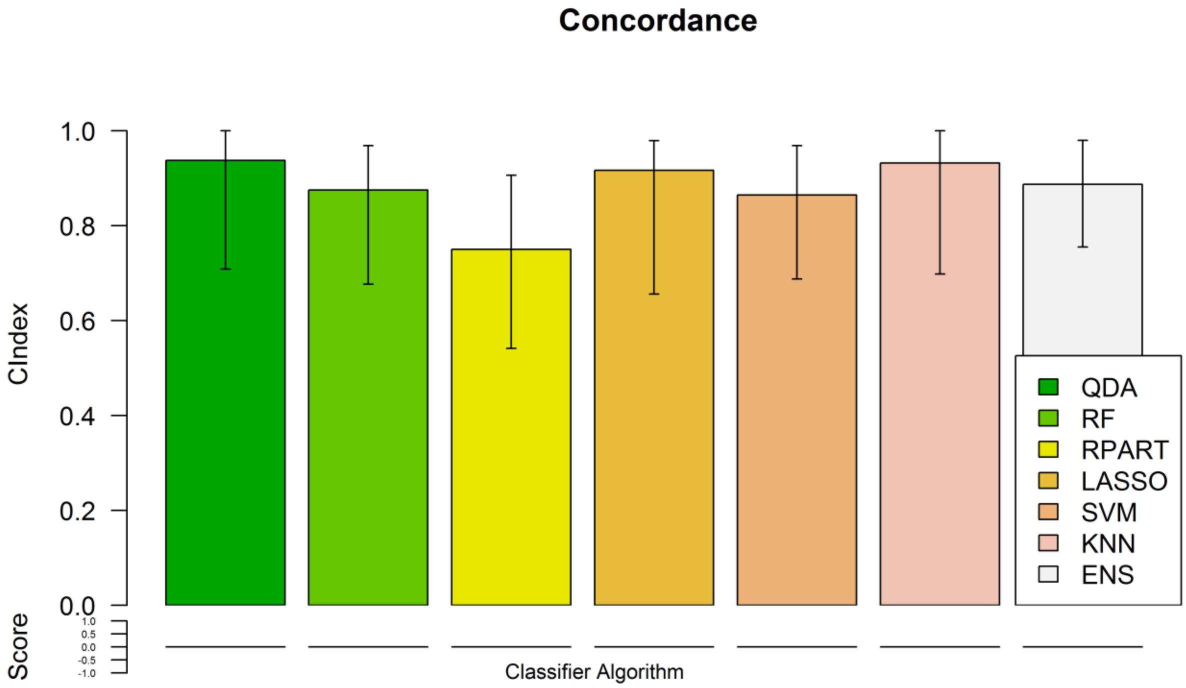
Concordance
1.0
0.8
0.6
0.4
0.2
0.0
CIndex
Score
1.0
0.5
0.0
-0.5
-1.0
Classifier Algorithm
QDA
RF
RPART
LASSO
SVM
KNN
ENS

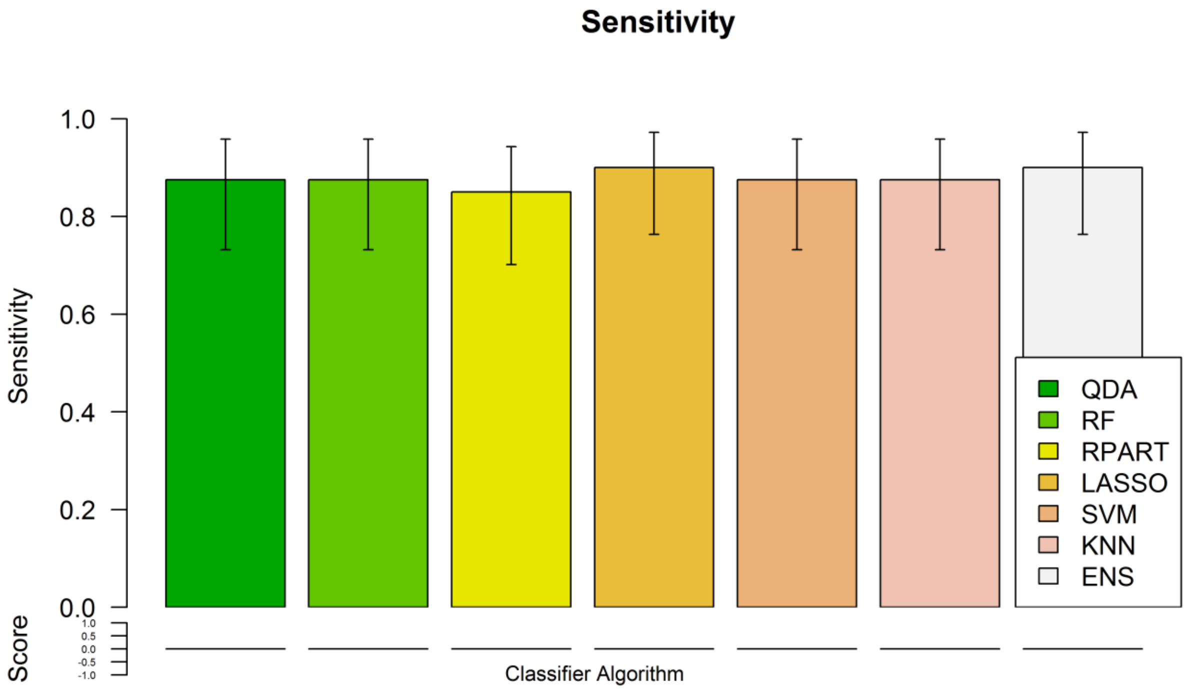
Sensitivity
1.0
0.8
0.6
0.4
0.2
0.0
Sensitivity
Score
1.0
0.5
0.0
-0.5
-1.0
Classifier Algorithm
QDA
RF
RPART
LASSO
SVM
KNN
ENS

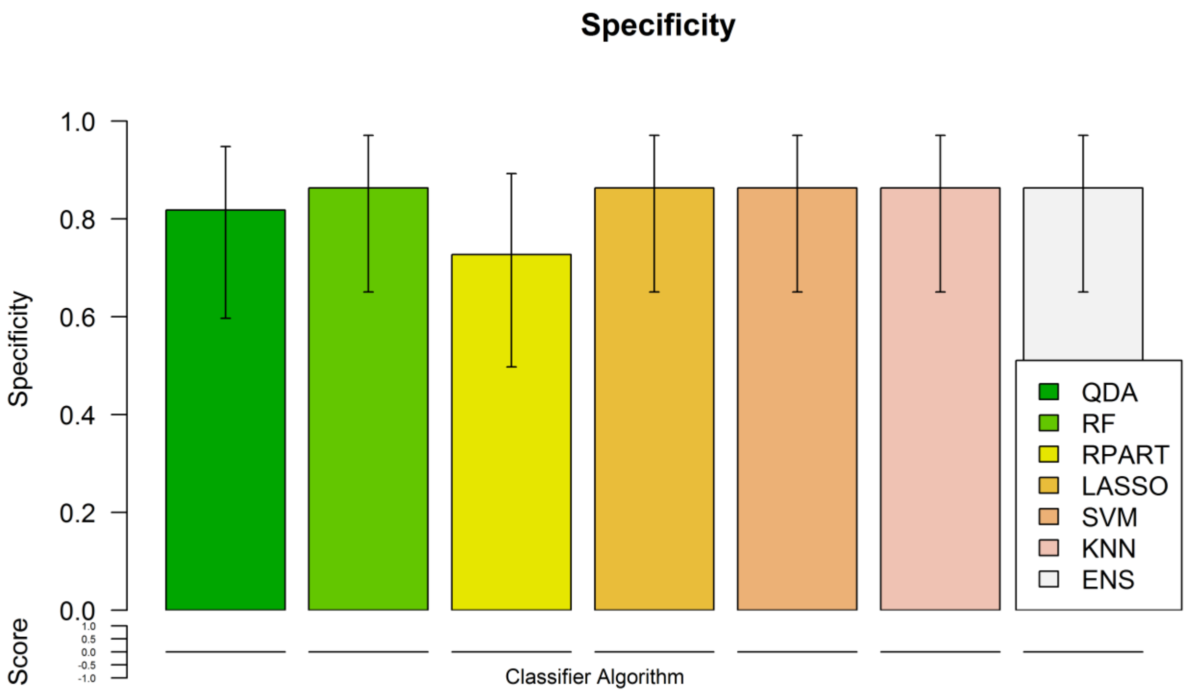

Specificity
1.0
0.8
0.6
0.4
0.2
0.0
Specificity
QDA
RF
RPART
LASSO
SVM
KNN
ENS
Score
Classifier Algorithm

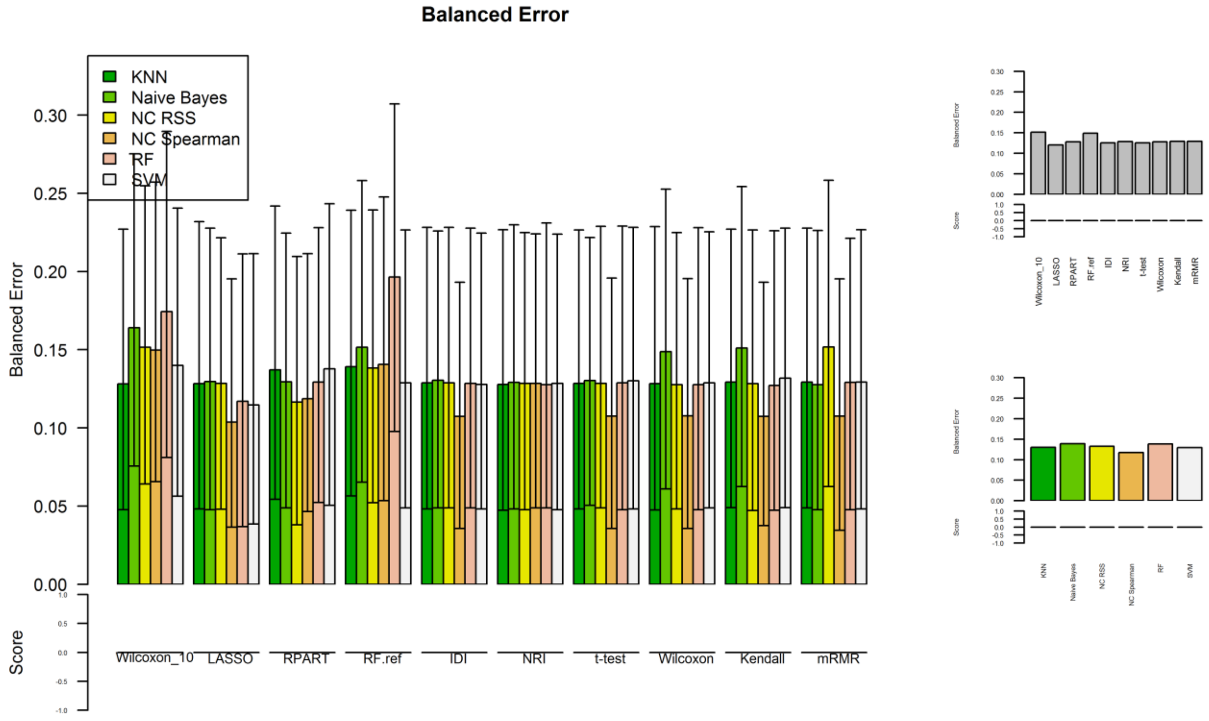

Balanced Error
KNN
Naive Bayes
NC RSS
NC Spearman
RF
SVM
0.30
0.25
0.20
0.15
0.10
0.05
0.00
Balanced Error
Score
Wilcoxon_10
LASSO
RPART
RF.ref
IDI
NRI
t-test
Wilcoxon
Kendall
mRMR

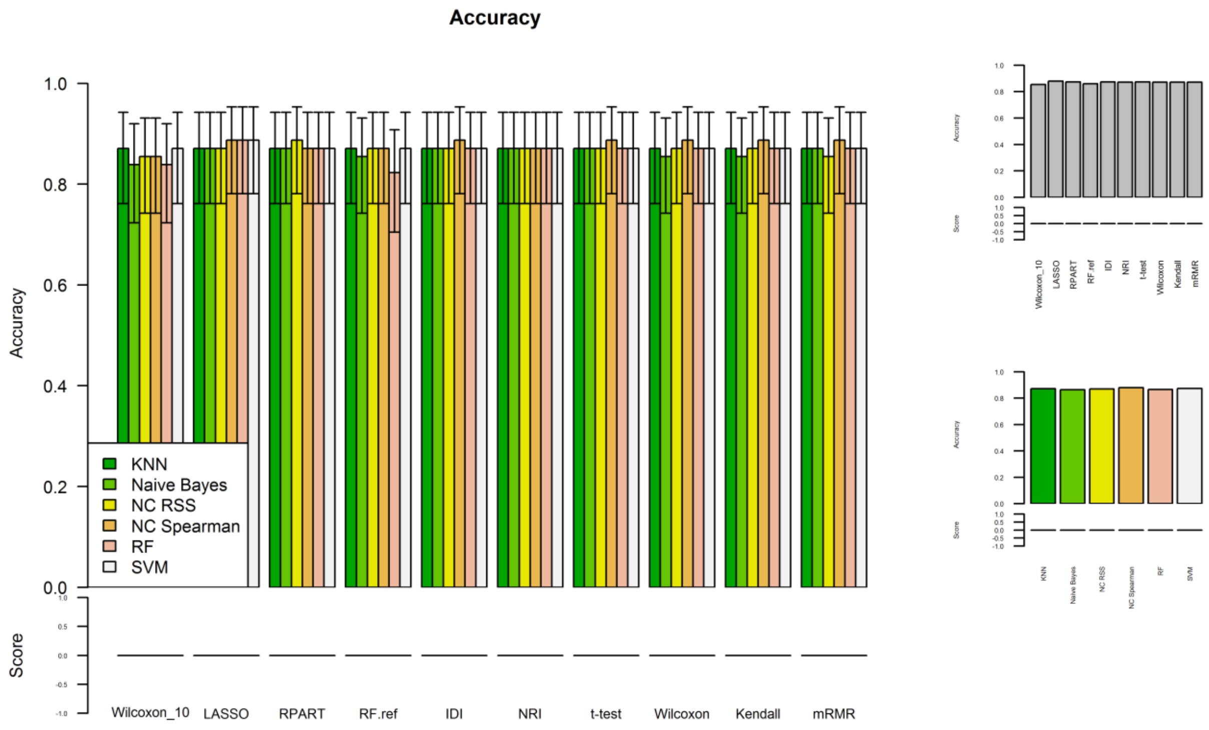
Accuracy
Accuracy
Score
1.0
0.8
0.6
0.4
0.2
0.0
KNN
Naive Bayes
NC RSS
NC Spearman
RF
SVM
Wilcoxon_10
LASSO
RPART
RF.ref
IDI
NRI
t-test
Wilcoxon
Kendall
mRMR

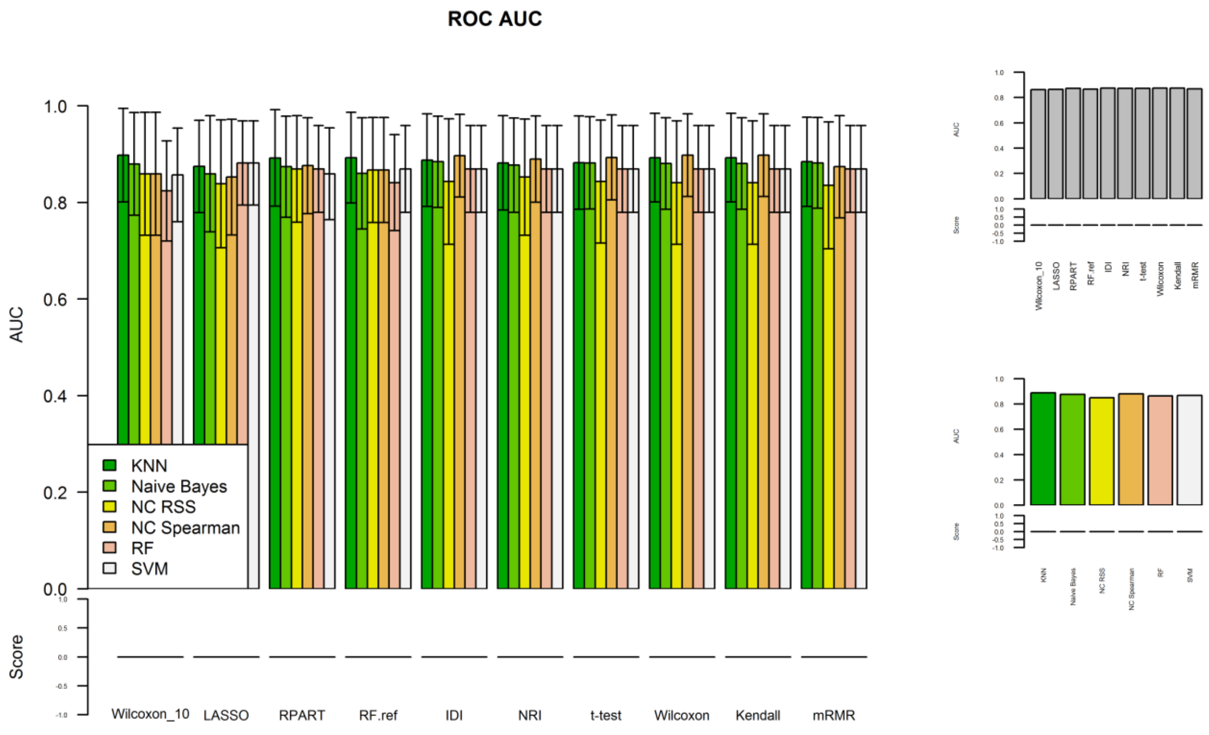
ROC AUC
AUC
Score
1.0
0.8
0.6
0.4
0.2
0.0
KNN
Naive Bayes
NC RSS
NC Spearman
RF
SVM
Wilcoxon_10
LASSO
RPART
RF.ref
IDI
NRI
t-test
Wilcoxon
Kendall
mRMR

#### Concordance

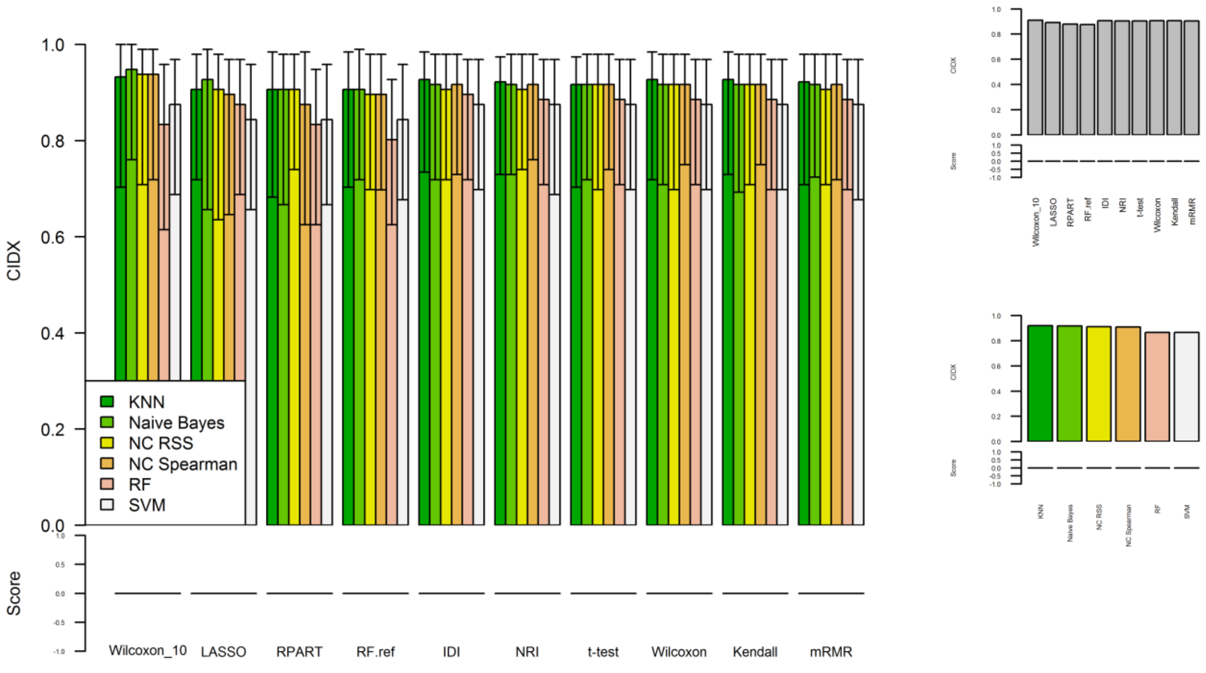

#### Sensitivity

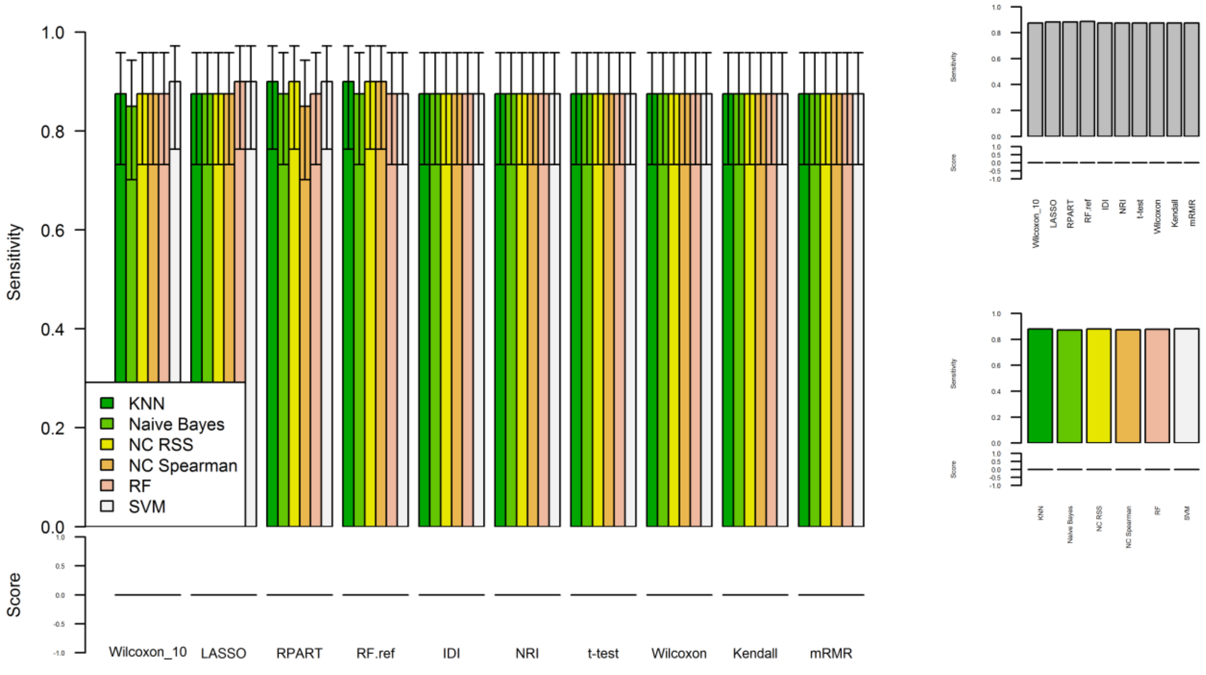

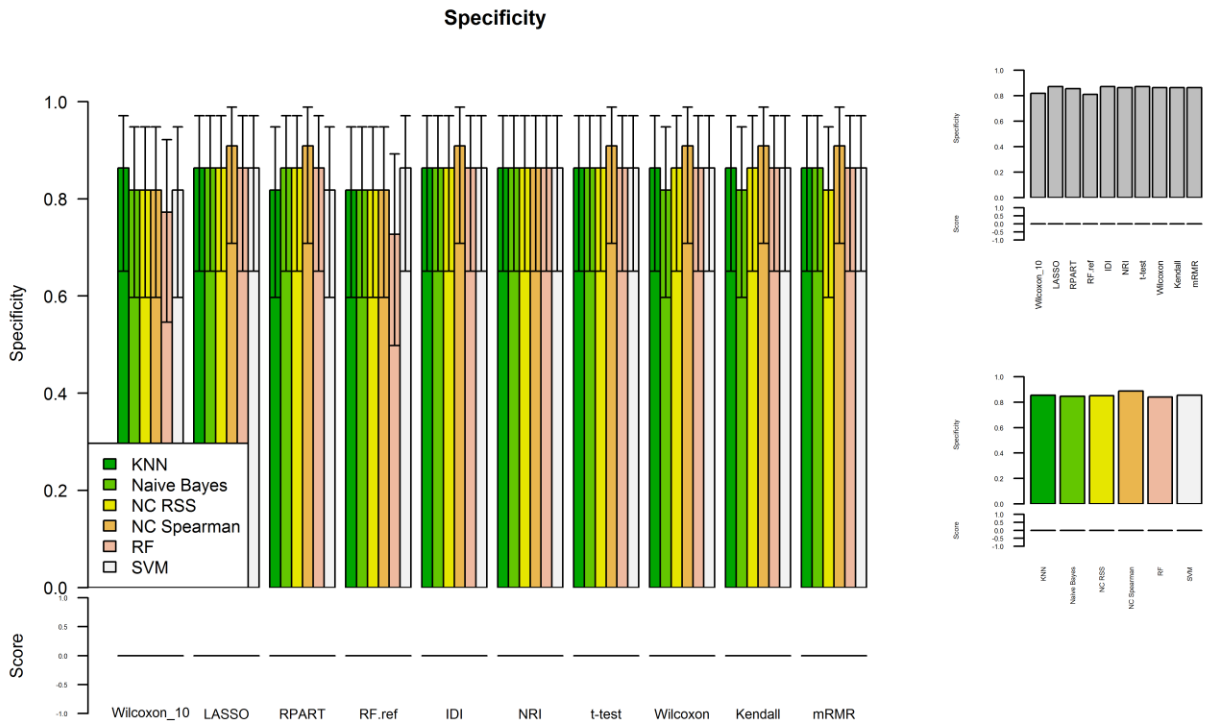

```r
par(mfrow = c(1,1),cex = 1.0)
pander::pander(prBenchmark$metrics,caption = "Clasifier Performance",round = 3)
```

*Clasifier Performance*

| | QDA | RF | RPART | LASSO | SVM | KNN | ENS |
|---|---|---|---|---|---|---|---|
| **BER** | 0.152 | 0.128 | 0.209 | 0.116 | 0.128 | 0.129 | 0.114 |
| **ACC** | 0.855 | 0.871 | 0.806 | 0.887 | 0.871 | 0.871 | 0.887 |
| **AUC** | 0.877 | 0.869 | 0.827 | 0.833 | 0.869 | 0.898 | 0.883 |
| **SEN** | 0.875 | 0.875 | 0.85 | 0.9 | 0.875 | 0.875 | 0.9 |
| **SPE** | 0.818 | 0.864 | 0.727 | 0.864 | 0.864 | 0.864 | 0.864 |
| **CIDX** | 0.938 | 0.875 | 0.75 | 0.917 | 0.865 | 0.932 | 0.887 |

```r
pander::pander(prBenchmark$metrics_filter,caption = "Average Filter Performance",round = 3)
```

*Average Filter Performance*

| | Wilcoxon_10 | LASSO | RPART | RF.ref | IDI | NRI | t-test | Wilcoxon | Kendall | mRMR |
|---|---|---|---|---|---|---|---|---|---|---|
| **BER** | 0.151 | 0.123 | 0.129 | 0.14 | 0.129 | 0.128 | 0.129 | 0.128 | 0.129 | 0.129 |
| **ACC** | 0.855 | 0.879 | 0.871 | 0.871 | 0.871 | 0.871 | 0.871 | 0.871 | 0.871 | 0.871 |
| **AUC** | 0.859 | 0.867 | 0.872 | 0.867 | 0.877 | 0.873 | 0.876 | 0.875 | 0.875 | 0.872 |
| **SEN** | 0.875 | 0.875 | 0.888 | 0.888 | 0.875 | 0.875 | 0.875 | 0.875 | 0.875 | 0.875 |
| **SPE** | 0.818 | 0.864 | 0.864 | 0.818 | 0.864 | 0.864 | 0.864 | 0.864 | 0.864 | 0.864 |
| **CIDX** | 0.935 | 0.901 | 0.891 | 0.896 | 0.911 | 0.911 | 0.917 | 0.917 | 0.917 | 0.911 |
