## Supplementary material for "Benchmarking machine learning models for the analysis of genetic data using FRESA.CAD Binary Classification Benchmarking": Type 2 Diabetes Analysis

### Benchmarks for Diabetes SNP Data Set

*Javier de Velasco*

*March 13, 2019*


- 1 FRESA.CAD Benchmark
  - 1.1 Diabetes SNP Data Set
  - 1.2 Benchmarking
  - 1.3 Results
    - 1.3.1 Classifier Results
    - 1.3.2 Radar Plots
    - 1.3.3 Feature Analysis


### 1 FRESA.CAD Benchmark

#### 1.1 Diabetes SNP Data Set

```
DiabetesData <- read.csv("FresaCAD/DiabSnps1000Fix.txt", sep = "", na.strings = "-1", str
ingsAsFactors=FALSE)

Diabetes <- as.data.frame(DiabetesData[,-1])
rownames(Diabetes) <- DiabetesData[,1]
Diabetes[,1:ncol(Diabetes)] <- sapply(Diabetes,as.numeric)

ExperimentName <- "Diabetes"
bswimsReps <- 20;
theData <- Diabetes;
theData <- as.data.frame(1.0*(theData > 0))


theData$Class <- 1.0*Diabetes$Class
theData <- nearestNeighborImpute(as.data.frame(theData))
theOutcome <- "Class";
reps <- 100;
fraction <- 0.9;

#sf <- univariate_Wilcoxon(data=theData, Outcome="Class",pvalue=0.5)
#theData <- theData[,c("Class",names(sf))]



BSWiMSFileName <- paste(ExperimentName,"BSWIMSMethod.RDATA",sep = "_")
CVFileName <- paste(ExperimentName,"CVMethod.RDATA",sep = "_")
BSCVFileName <- paste(ExperimentName,"BSCVMethod.RDATA",sep = "_")
```

#### 1.2 Benchmarking

```
BSWiMSMODEL <- BSWiMS.model(formula = paste(theOutcome," ~ 1"),data = theData,NumberofRep
eats = bswimsReps)
save(BSWiMSMODEL,file = BSWiMSFileName)


load(file = BSWiMSFileName)

par(mfrow = c(2,2),cex=0.6);
cp <- BinaryBenchmark(theData,theOutcome,reps,fraction)
```

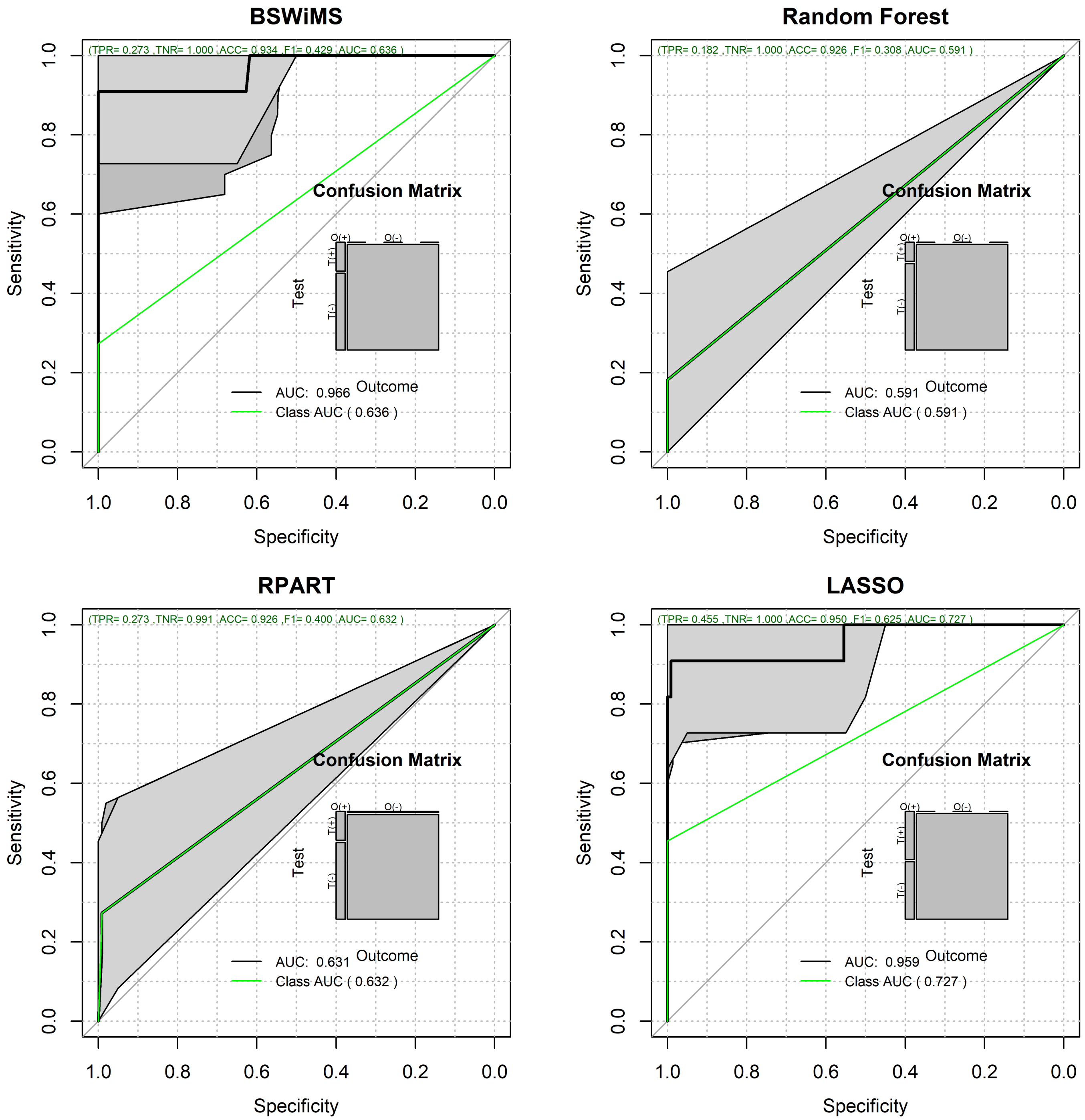


```
save(cp,file = CVFileName)
par(mfrow = c(1,1),cex=1.0);
```

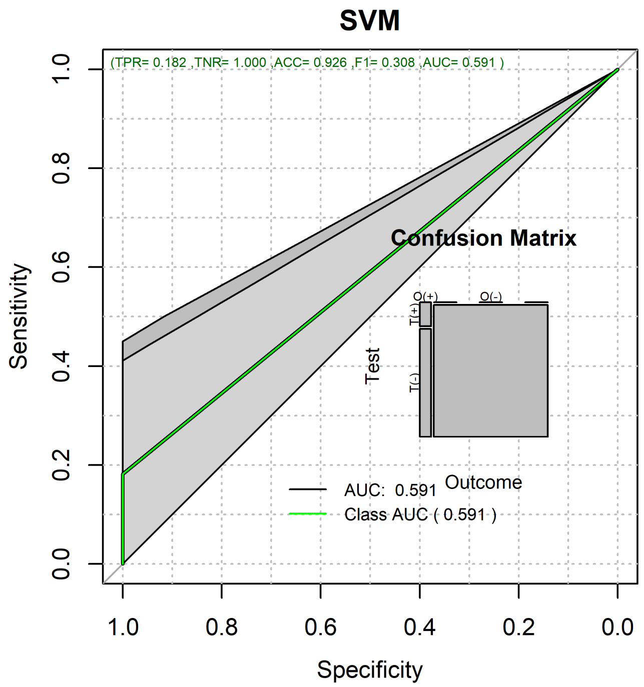


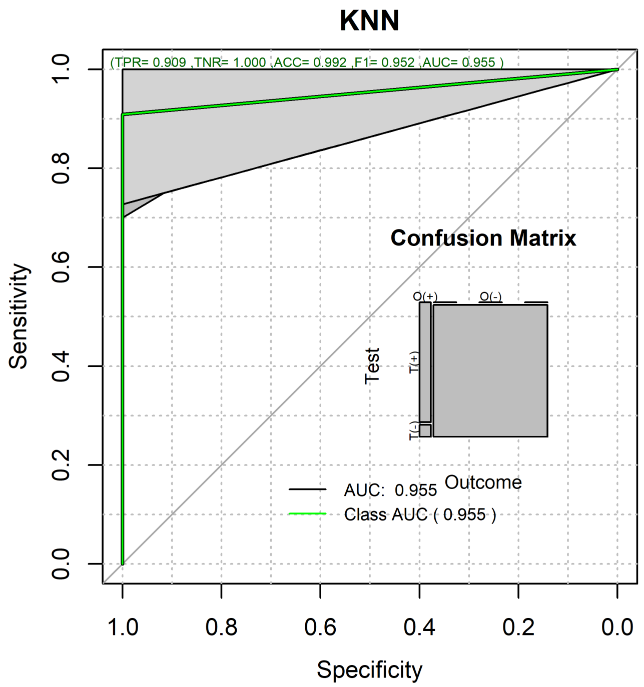


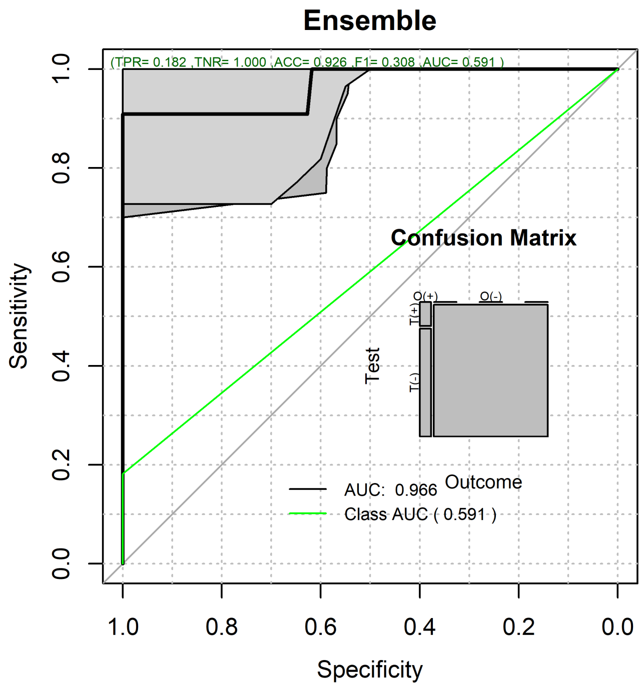


```
load(file = CVFileName)
```

### 1.3 Results

#### 1.3.1 Classifier Results

```
#hm <- heatMaps(Outcome = "Outcome",data = cp$testPredictions,title = "Heat Map",Scale =
 TRUE,hCluster = "col",cexRow = 0.25,cexCol = 0.75,srtCol = 45)

#The Times
par(mfrow = c(2,1),cex=1.0);
pander::pander(cp$cpuElapsedTimes)
```

| BSWiMS | RF | RPART | LASSO | SVM | KNN | ENS |
|---|---|---|---|---|---|---|
| 16.11 | 0.9882 | 0.1316 | 0.2693 | 0.015 | 0.0064 | 17.52 |

```
par(mfrow = c(1,1),cex=1.0);
learningTime <- -1*cp$cpuElapsedTimes
pr <- plot(cp)
```

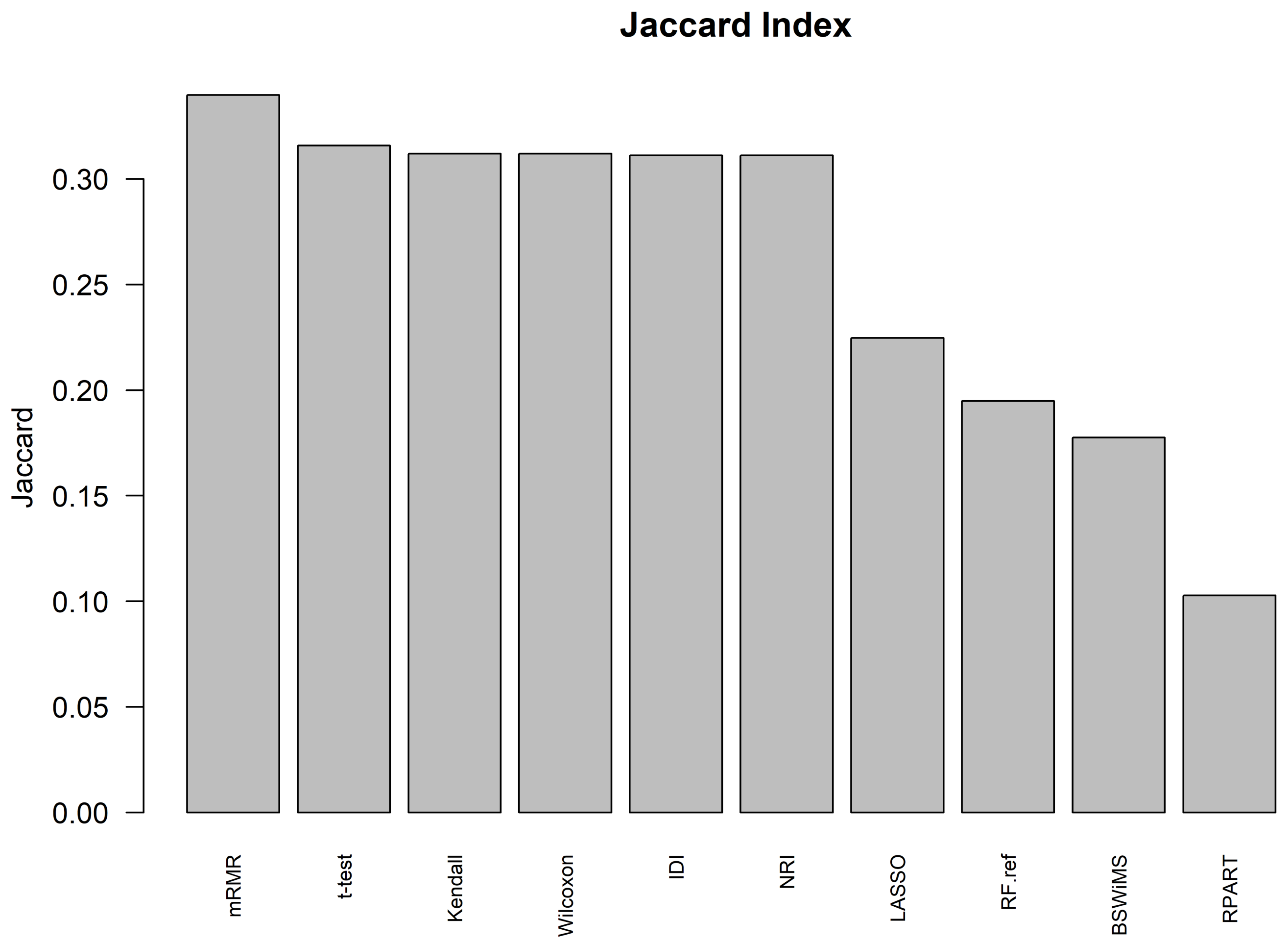
Jaccard Index
Jaccard
0.00
0.05
0.10
0.15
0.20
0.25
0.30
mRMR
t-test
Kendall
Wilcoxon
IDI
NRI
LASSO
RF.ref
BSWiMS
RPART

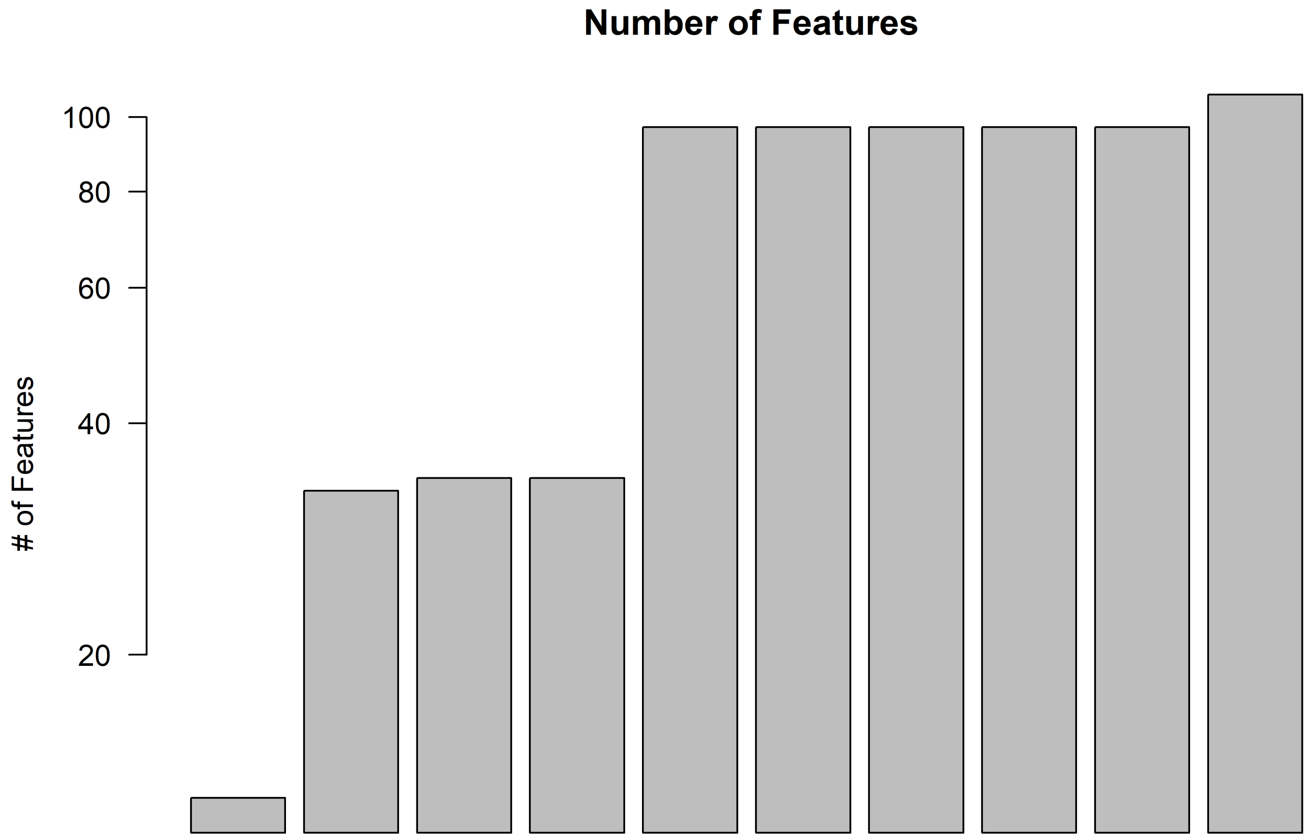
Number of Features
### of Features
20
40
60
80
100

RPART LASSO BSWiMS RF.ref IDI NRI t-test Wilcoxon Kendall mRMR

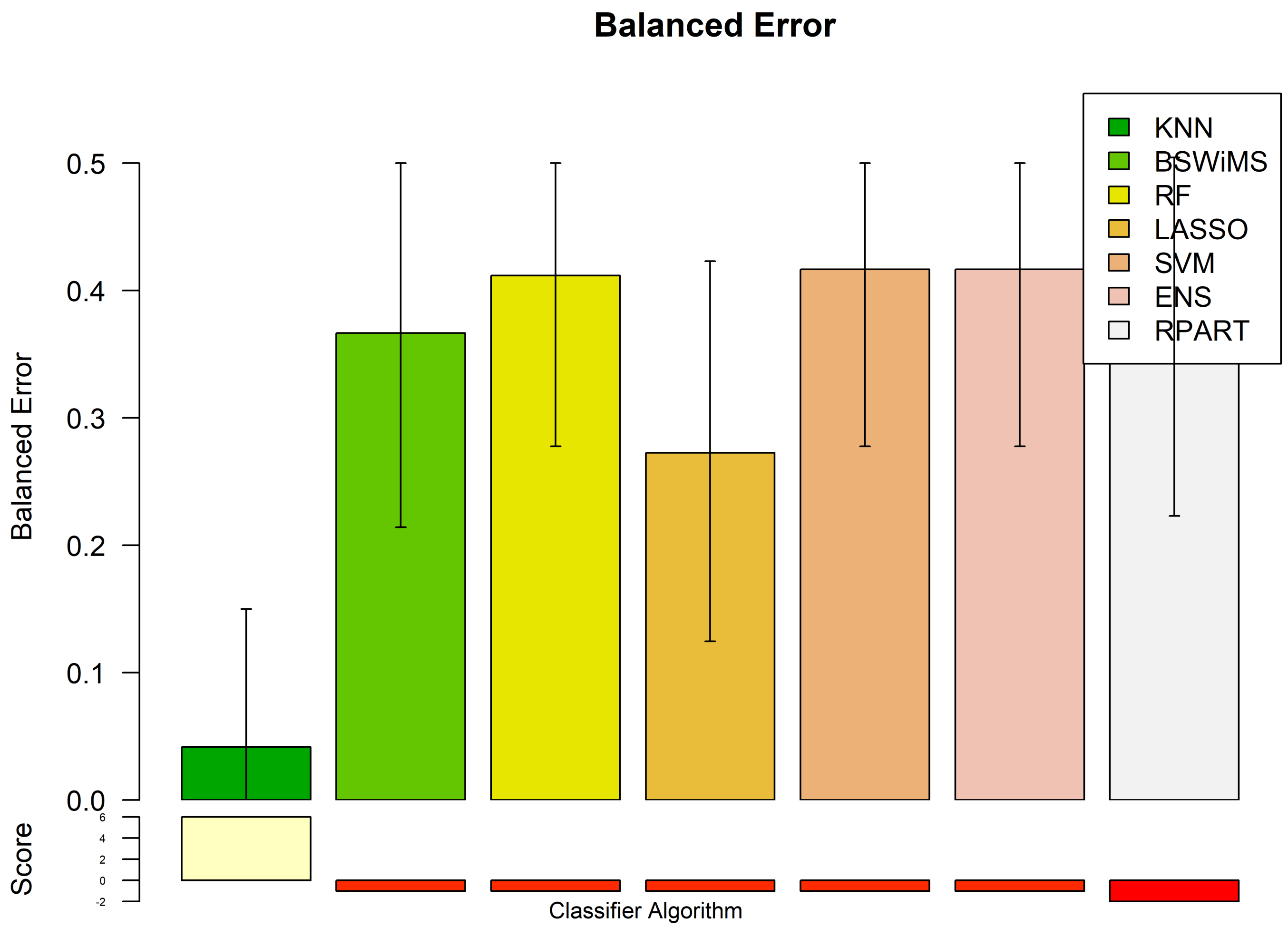

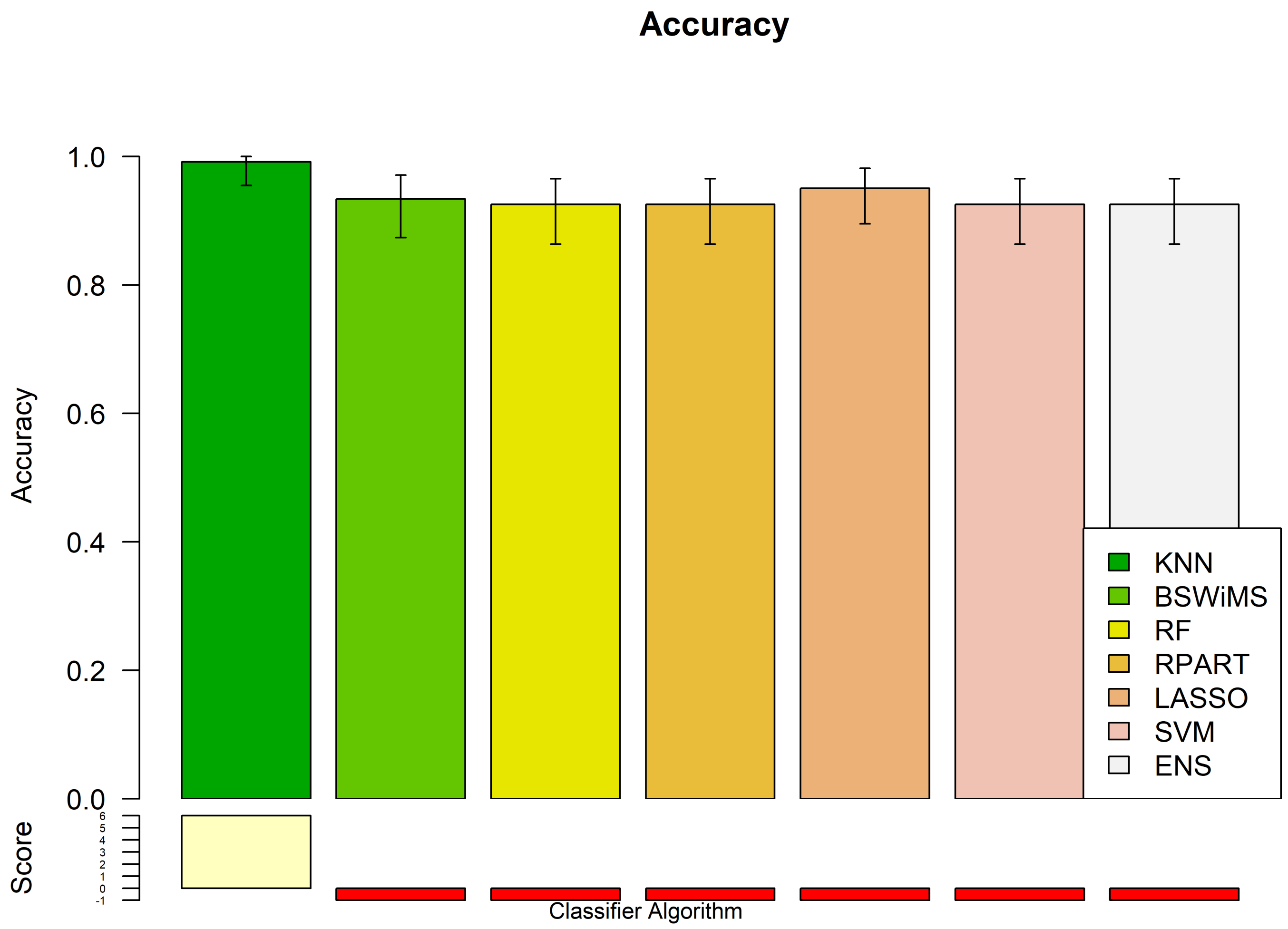
Accuracy
1.0
0.8
0.6
0.4
0.2
0.0
Accuracy
Score
KNN
BSWiMS
RF
RPART
LASSO
SVM
ENS
Classifier Algorithm

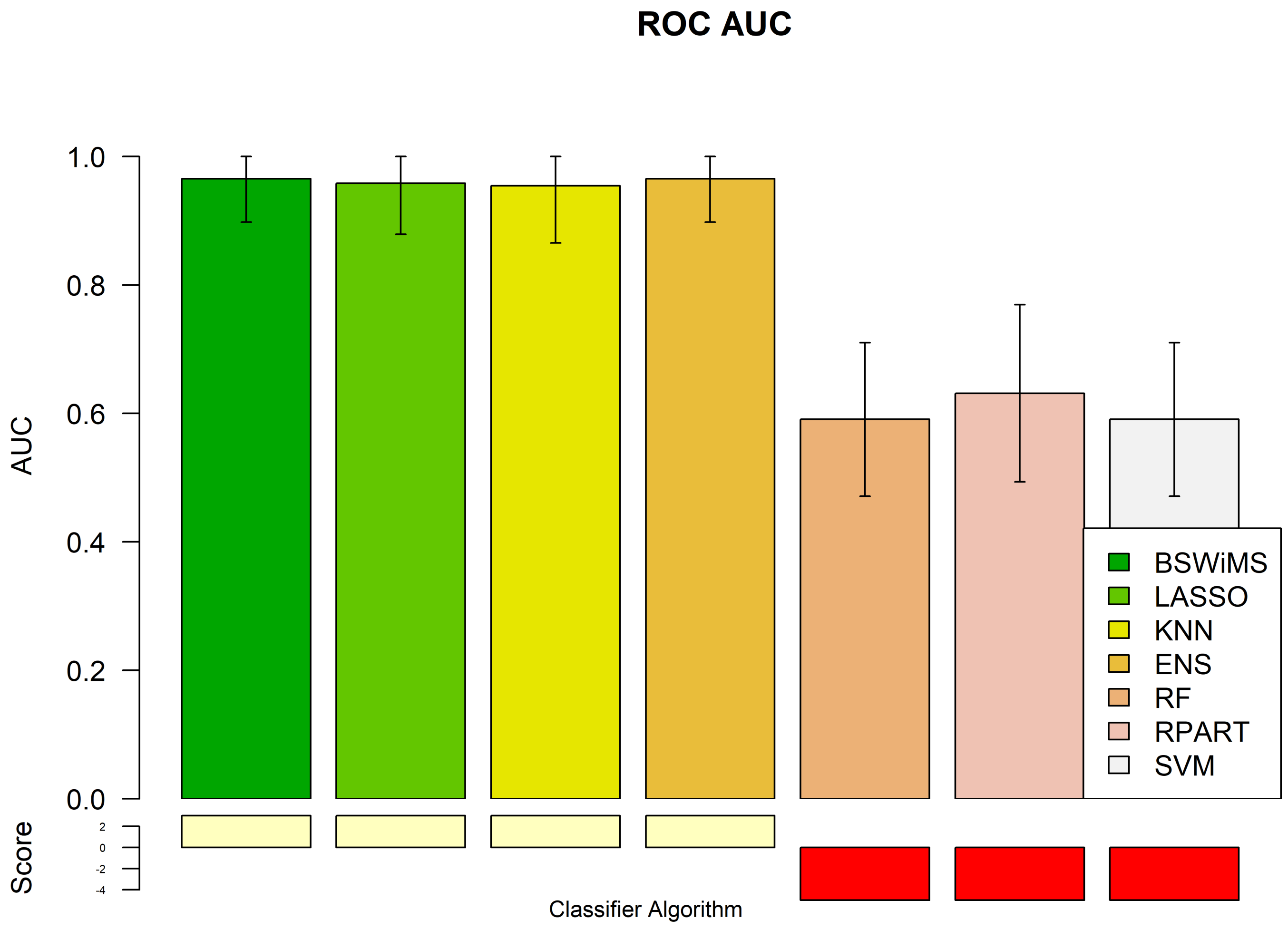
ROC AUC
AUC
1.0
0.8
0.6
0.4
0.2
0.0
Score
2
0
-2
-4
Classifier Algorithm
BSWiMS
LASSO
KNN
ENS
RF
RPART
SVM

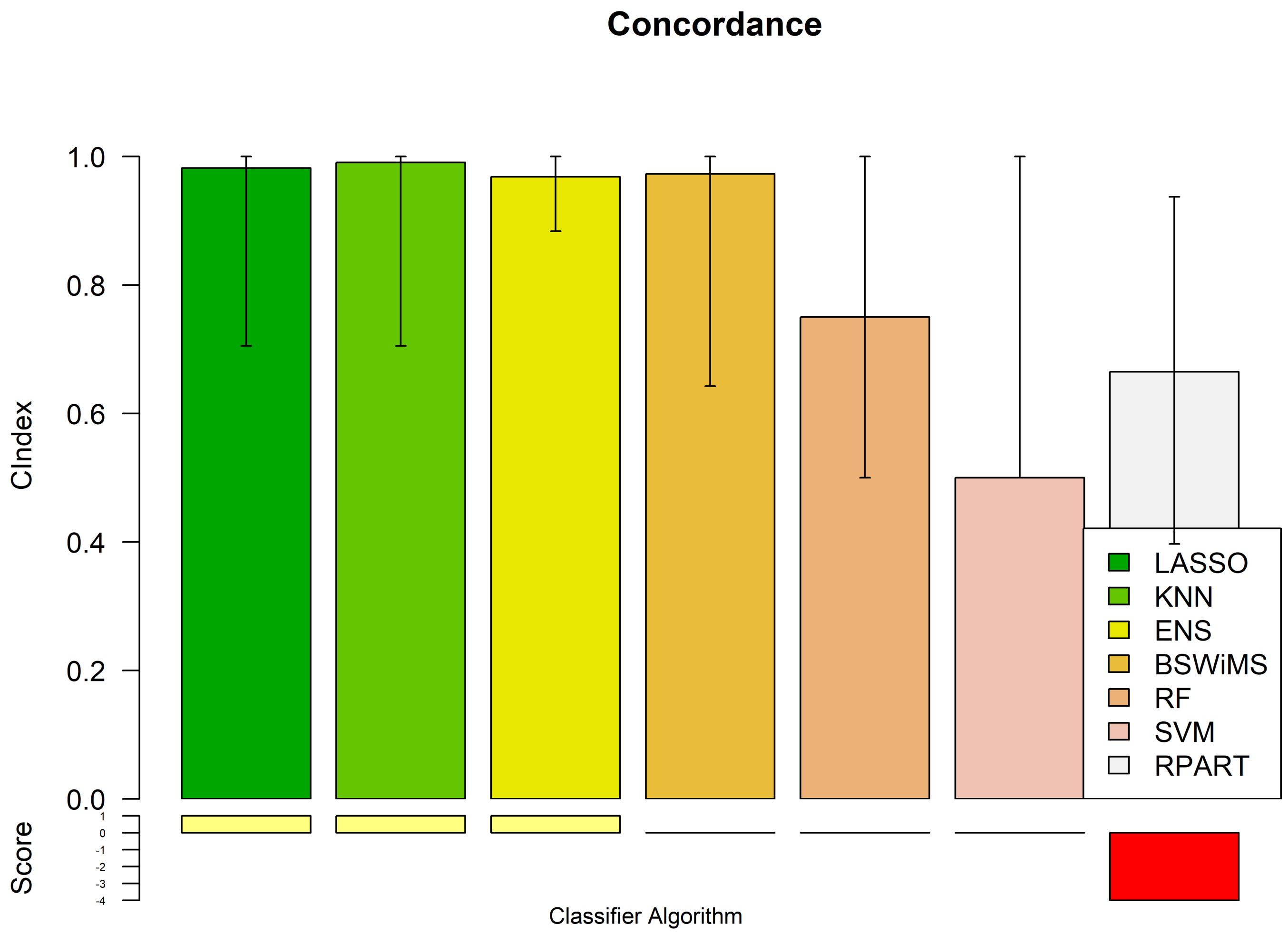
Concordance
1.0
0.8
0.6
0.4
0.2
0.0
CIndex
Score
1
0
-1
-2
-3
-4
LASSO
KNN
ENS
BSWiMS
RF
SVM
RPART
Classifier Algorithm

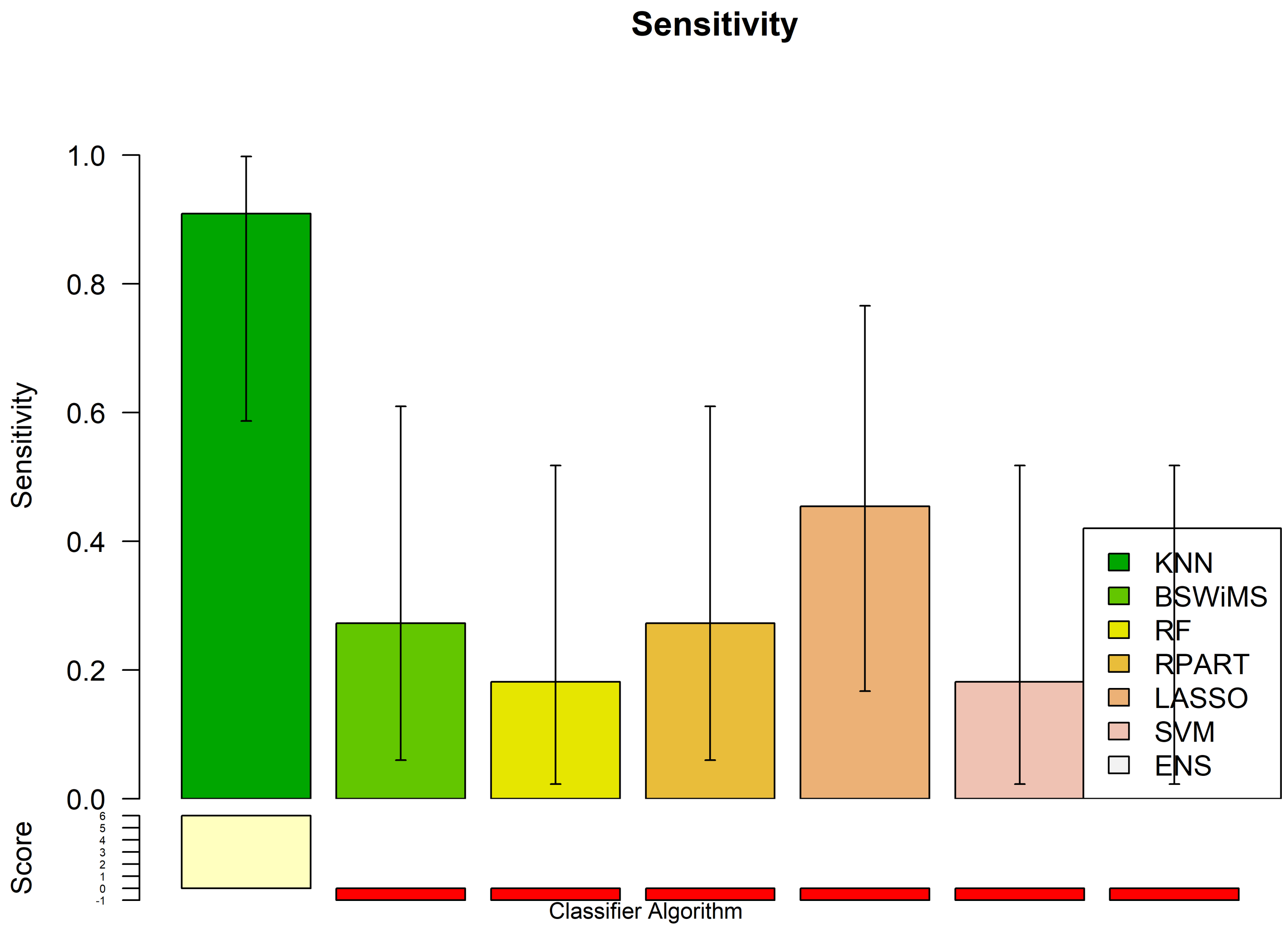
Sensitivity
1.0
0.8
0.6
0.4
0.2
0.0
Sensitivity
Score
6
5
4
3
2
1
0
-1
KNN
BSWiMS
RF
RPART
LASSO
SVM
ENS
Classifier Algorithm

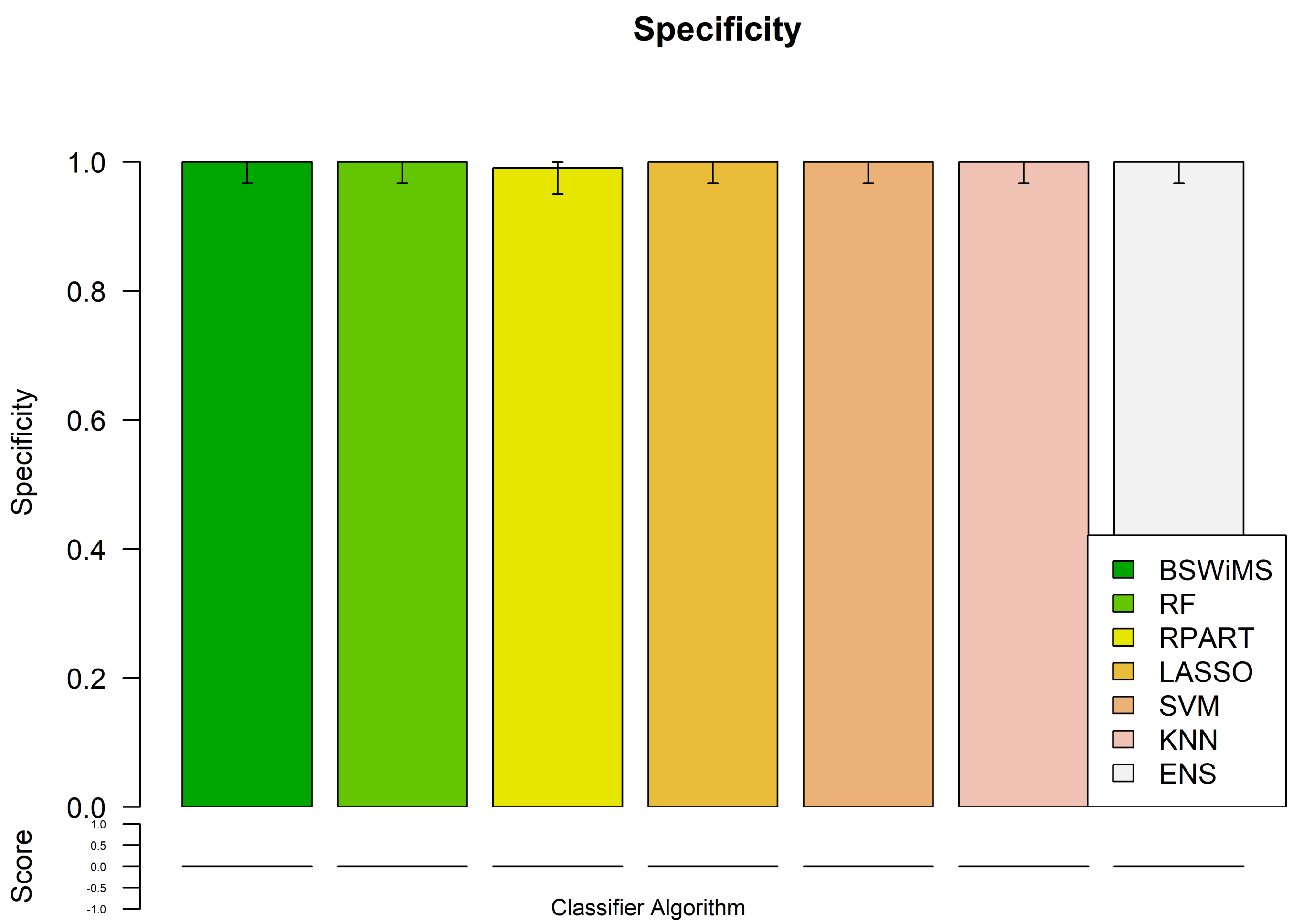
Specificity
Specificity
1.0
0.8
0.6
0.4
0.2
0.0
Score
1.0
0.5
0.0
-0.5
-1.0
BSWiMS
RF
RPART
LASSO
SVM
KNN
ENS
Classifier Algorithm

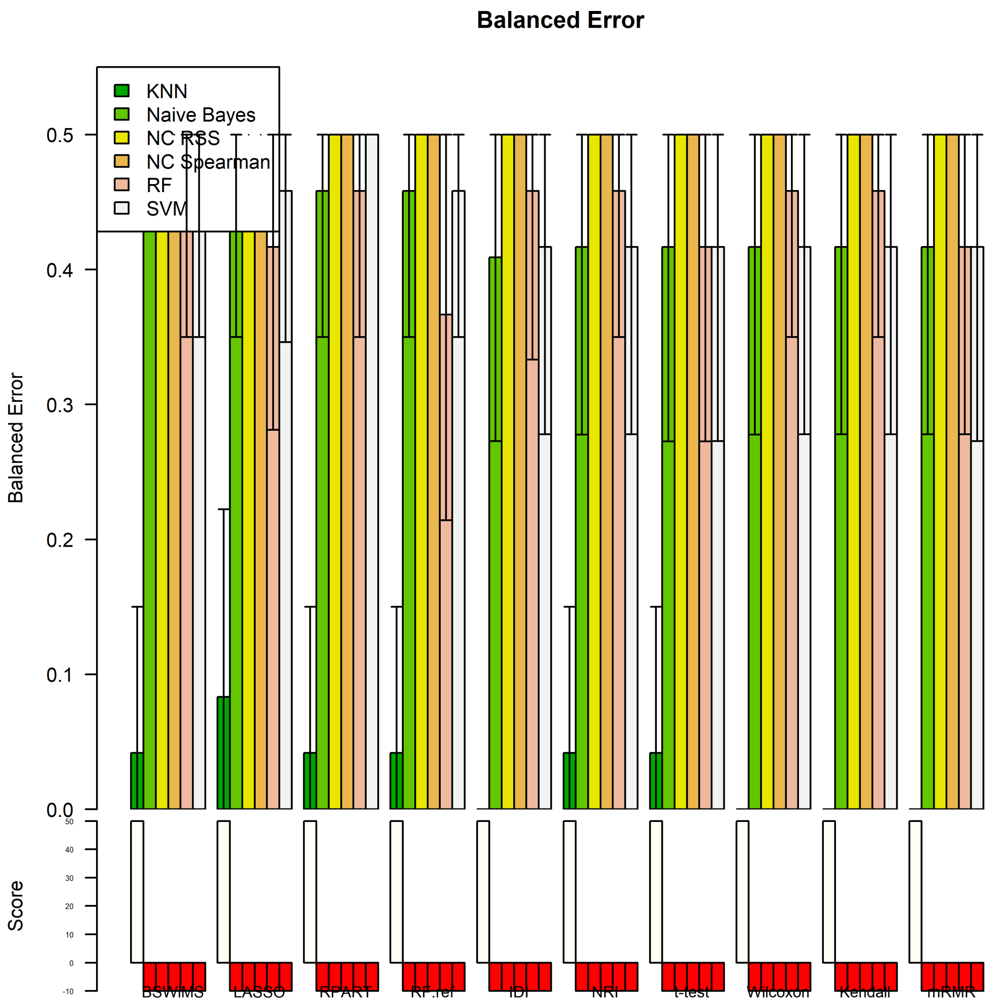
Balanced Error
KNN
Naive Bayes
NC RSS
NC Spearman
RF
SVM
Balanced Error
Score
BSWiMS
LASSO
RPART
RF.ref
IDI
NRI
t-test
Wilcoxon
Kendall
mRMR

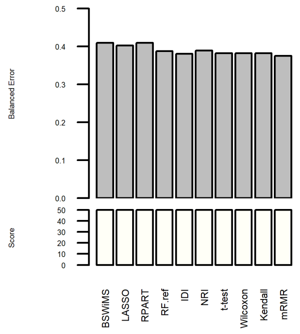
Balanced Error
Score
BSWiMS
LASSO
RPART
RF.ref
IDI
NRI
t-test
Wilcoxon
Kendall
mRMR

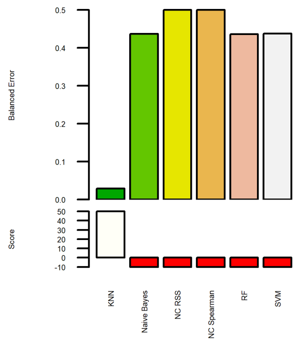
Balanced Error
Score
KNN
Naive Bayes
NC RSS
NC Spearman
RF
SVM

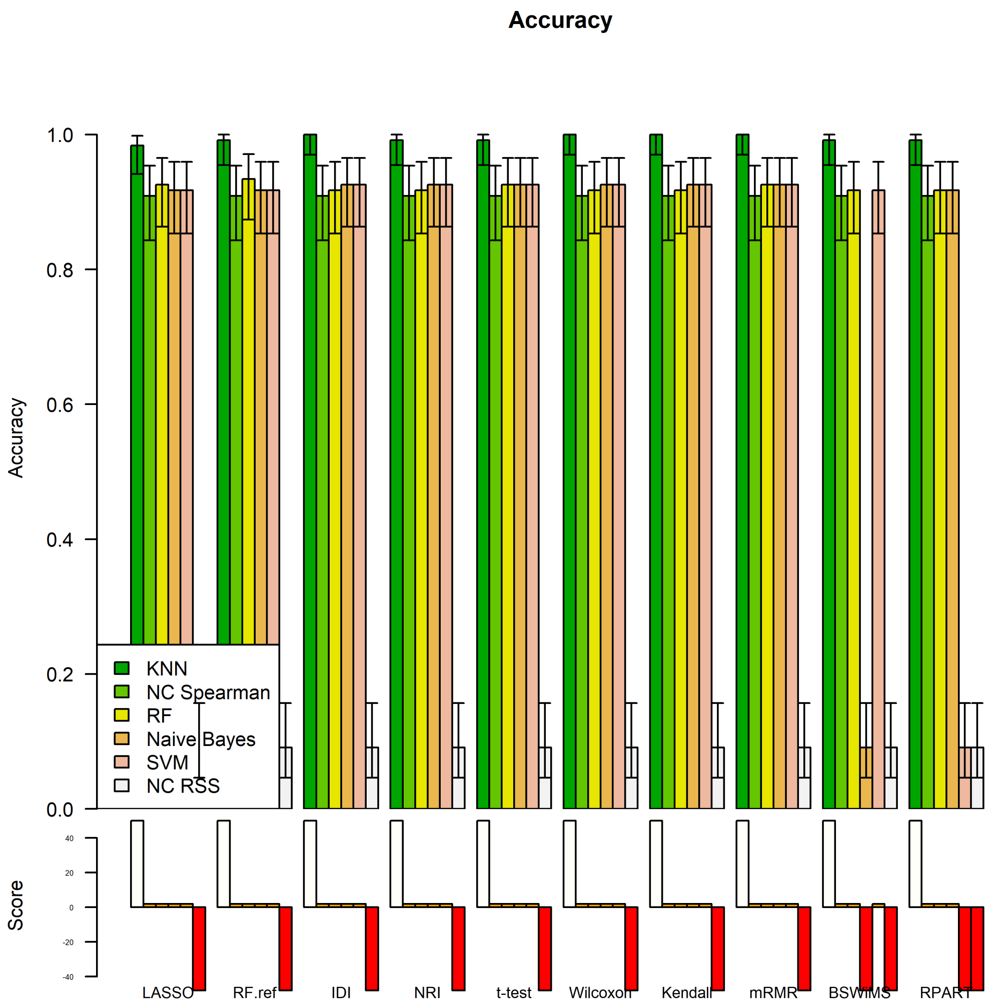
Accuracy
1.0
0.8
0.6
0.4
0.2
0.0
Accuracy
KNN
NC Spearman
RF
Naive Bayes
SVM
NC RSS
Score
LASSO
RF.ref
IDI
NRI
t-test
Wilcoxon
Kendall
mRMR
BSWiMS
RPART

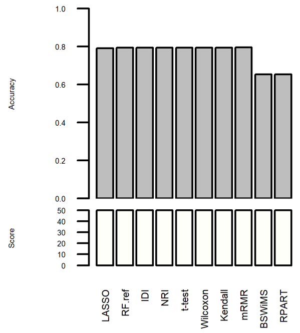
Accuracy
Score
LASSO
RF.ref
IDI
NRI
t-test
Wilcoxon
Kendall
mRMR
BSWiMS
RPART

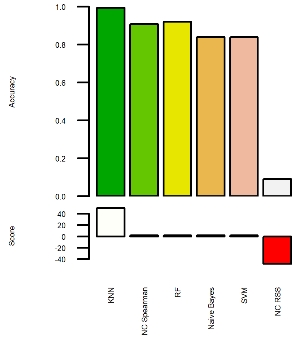
Accuracy
Score
KNN
NC Spearman
RF
Naive Bayes
SVM
NC RSS

**ROC AUC**

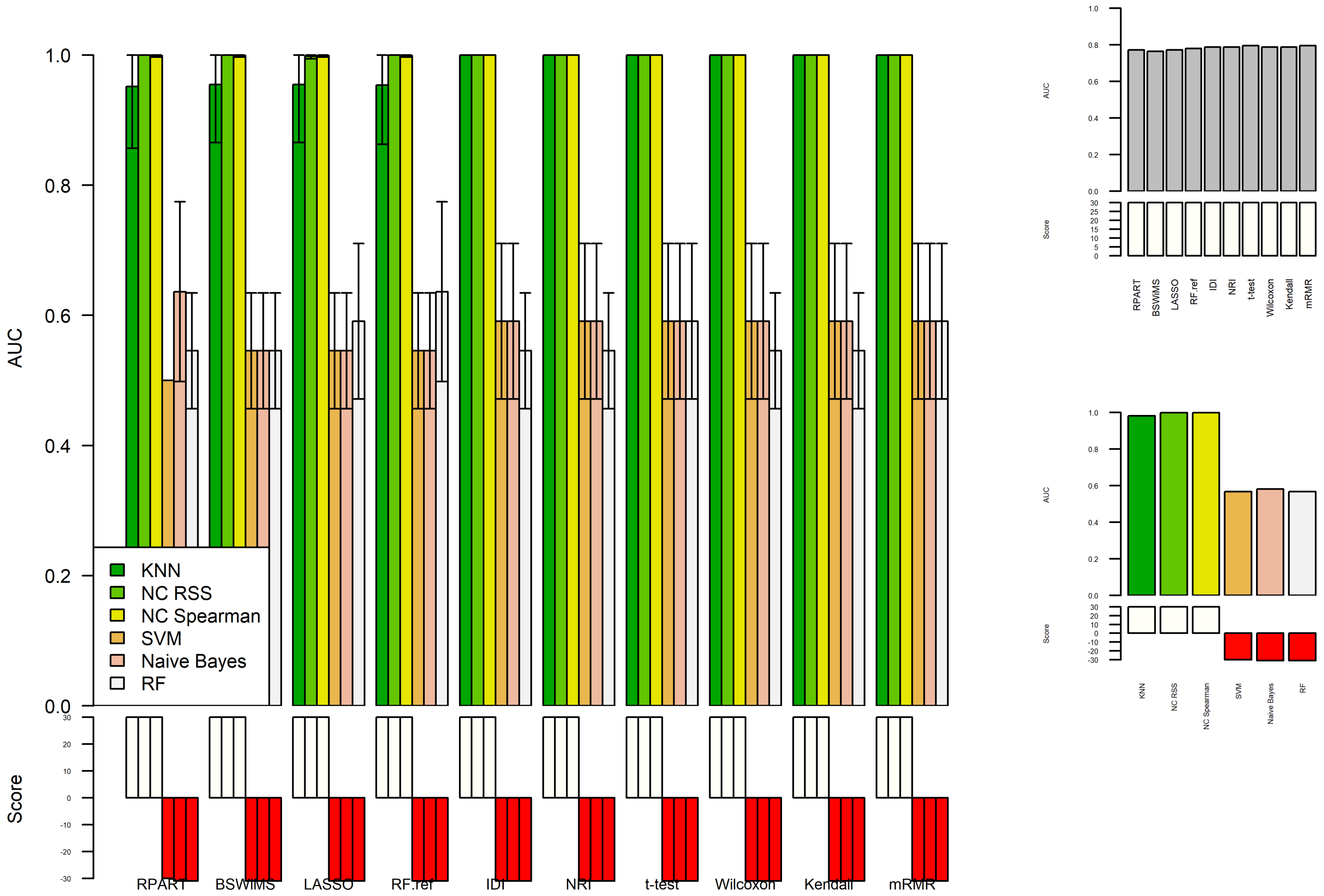
AUC
Score
KNN
NC RSS
NC Spearman
SVM
Naive Bayes
RF
RPART
BSWiMS
LASSO
RF.ref
IDI
NRI
t-test
Wilcoxon
Kendall
mRMR

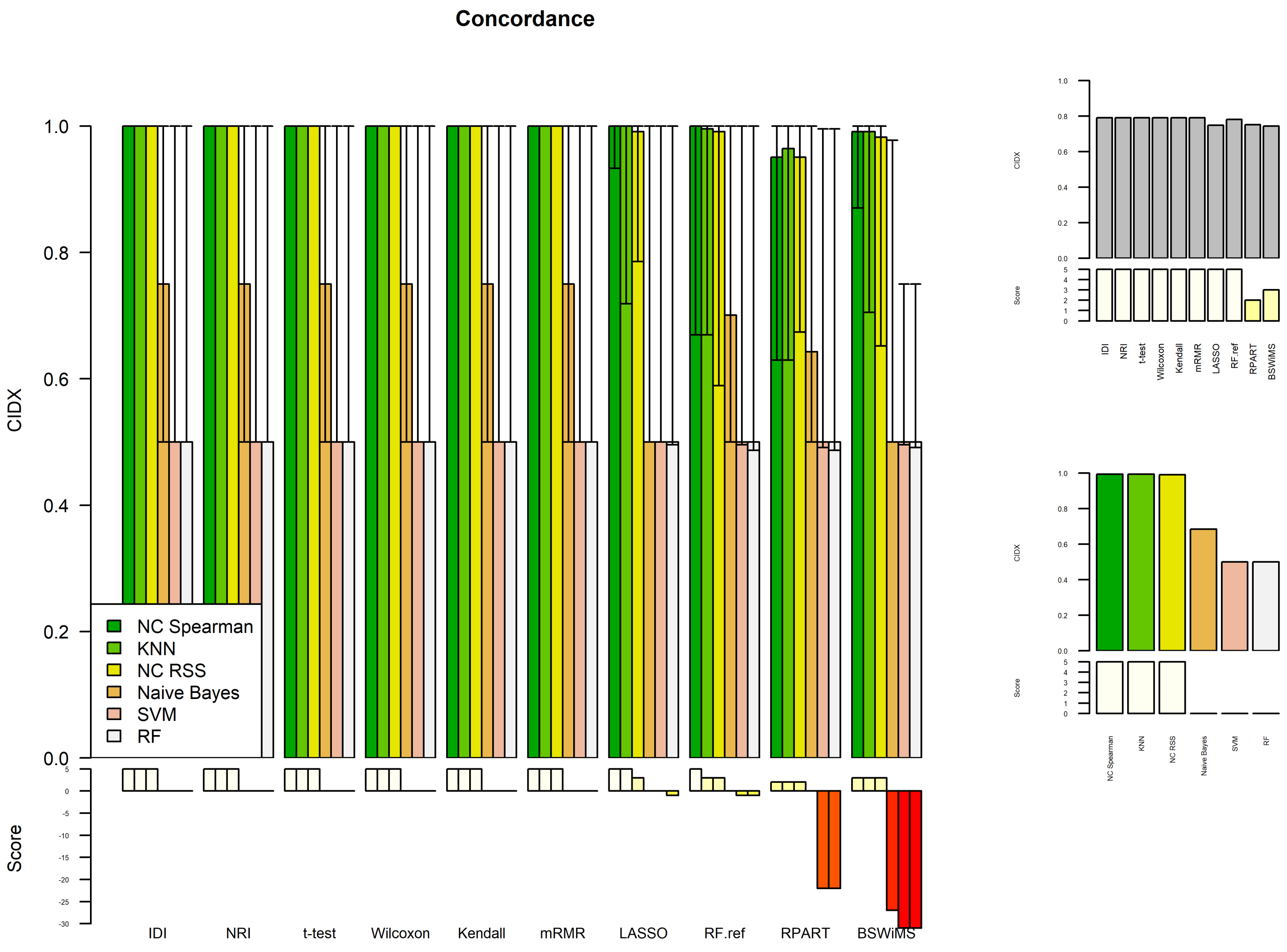
Concordance
CIDX
Score
NC Spearman
KNN
NC RSS
Naive Bayes
SVM
RF
IDI
NRI
t-test
Wilcoxon
Kendall
mRMR
LASSO
RF.ref
RPART
BSWiMS

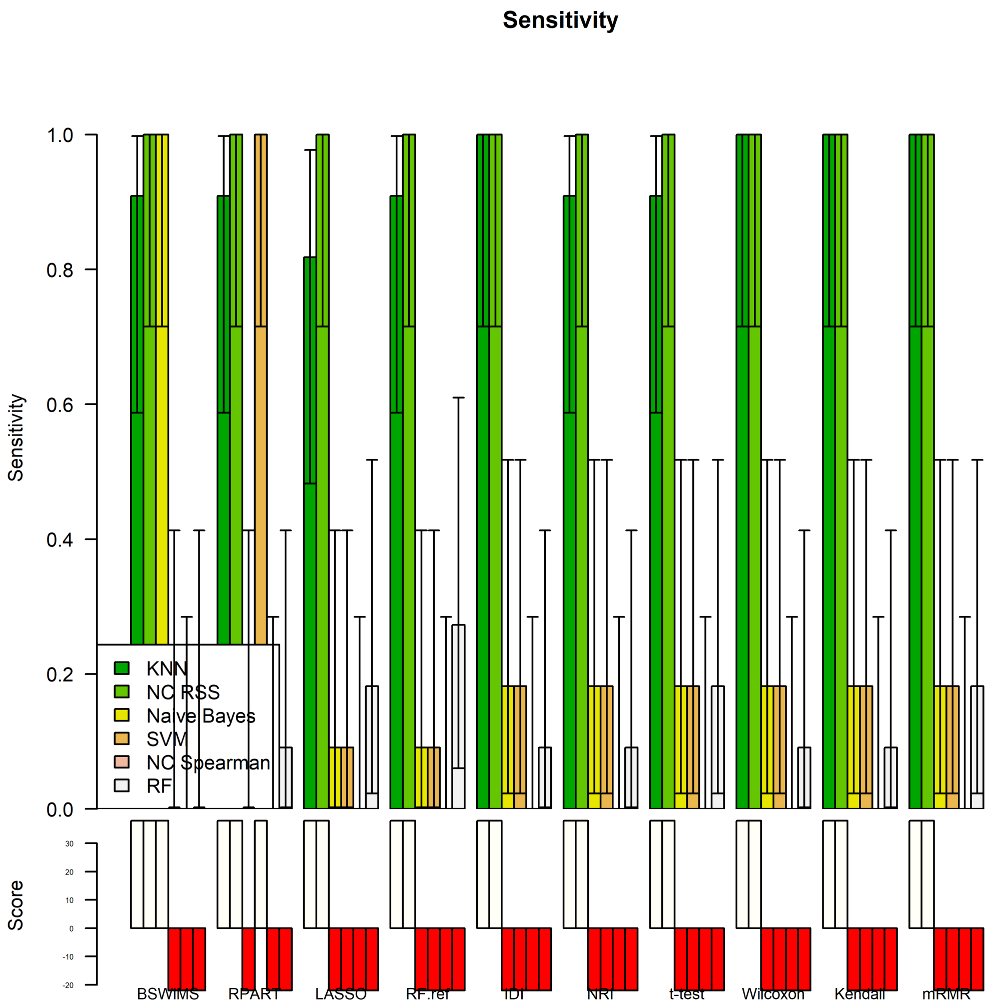
Sensitivity
Sensitivity
Score
KNN
NC RSS
Naive Bayes
SVM
NC Spearman
RF
BSWiMS
RPART
LASSO
RF.ref
IDI
NRI
t-test
Wilcoxon
Kendall
mRMR

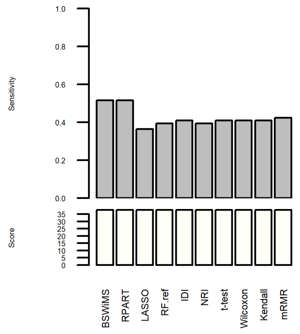
Sensitivity
Score
BSWiMS
RPART
LASSO
RF.ref
IDI
NRI
t-test
Wilcoxon
Kendall
mRMR

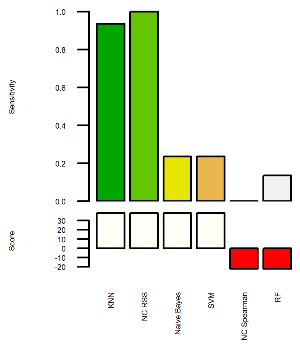
Sensitivity
Score
KNN
NC RSS
Naive Bayes
SVM
NC Spearman
RF

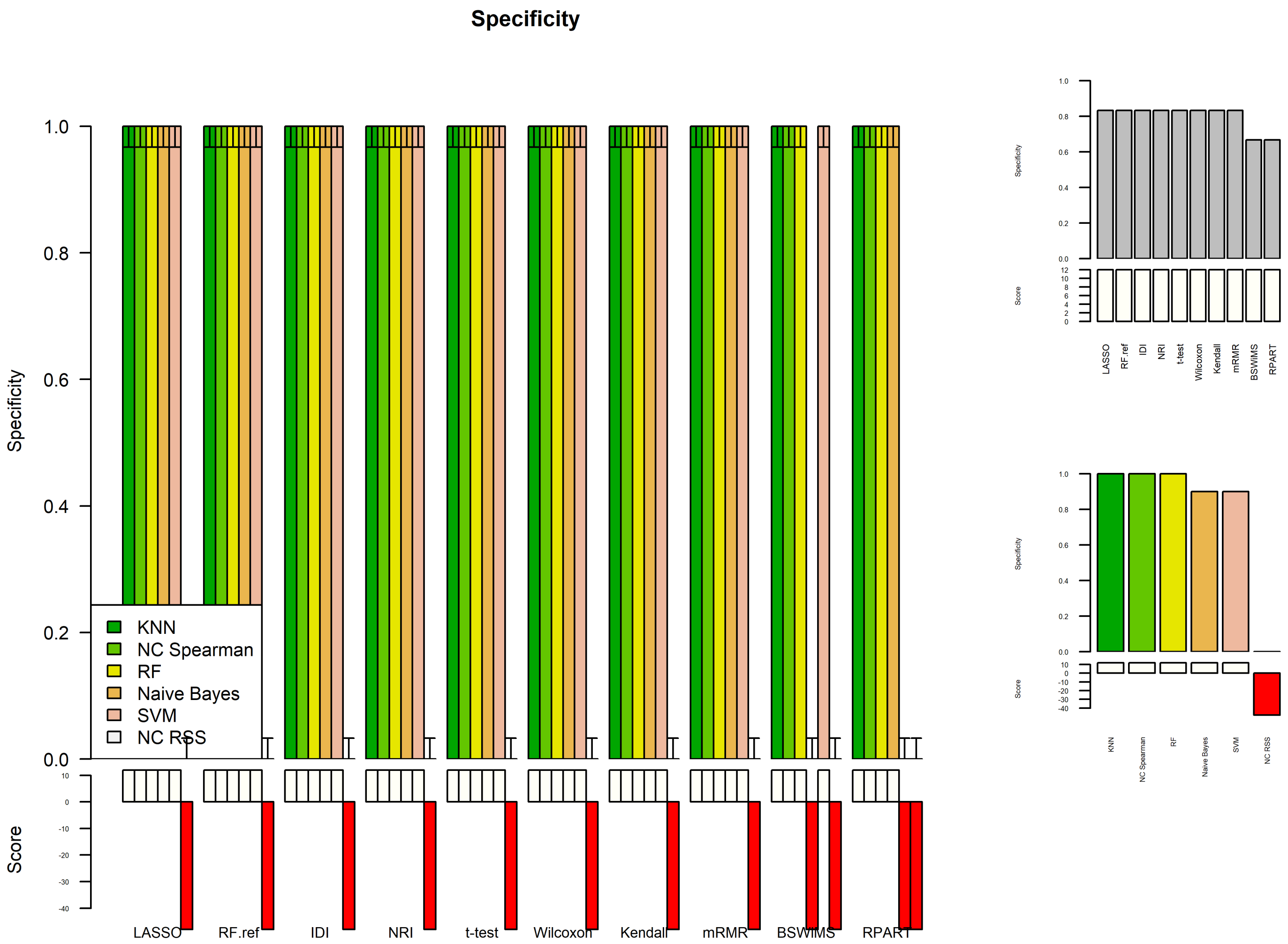


```
pr$metrics_filter
```

```
    BSWiMS     LASSO     RPART    RF.ref       IDI       NRI    t-test
```

BER 0.4791667 0.4583333 0.4791667 0.4583333 0.4375000 0.4375000 0.4166667 ACC 0.9132231 0.9173554 0.9132231 0.9173554 0.9214876 0.9214876 0.9256198 AUC 0.7500000 0.7727273 0.7940083 0.7950413 0.7954545 0.7954545 0.7954545 SEN 0.5000000 0.1363636 0.5000000 0.1818182 0.1818182 0.1818182 0.1818182 SPE 1.0000000 1.0000000 1.0000000 1.0000000 1.0000000 1.0000000 1.0000000 CIDX 0.7410714 0.7455357 0.7968750 0.8459821 0.8750000 0.8750000 0.8750000 Wilcoxon Kendall mRMR BER 0.4375000 0.4375000 0.4166667 ACC 0.9214876 0.9214876 0.9256198 AUC 0.7954545 0.7954545 0.7954545 SEN 0.1818182 0.1818182 0.1818182 SPE 1.0000000 1.0000000 1.0000000 CIDX 0.8750000 0.8750000 0.8750000

#### 1.3.2 Radar Plots

```
op <- par(no.readonly = TRUE)

library(fmsb)
par(mfrow = c(1,2),xpd = TRUE,pty = "s",mar = c(1,1,1,1))

mNames <- names(cp$cpuElapsedTimes)

classRanks <- c(pr$minMaxMetrics$BER[1],pr$minMaxMetrics$ACC[2],pr$minMaxMetrics$AUC[2],p
r$minMaxMetrics$CIDX[2],pr$minMaxMetrics$SEN[2],pr$minMaxMetrics$SPE[2],min(cp$cpuElapsed
Times))
classRanks <- rbind(classRanks,c(pr$minMaxMetrics$BER[2],0,0,0,0,0,max(cp$cpuElapsedTime
s)))
classRanks <- as.data.frame(rbind(classRanks,cbind(t(pr$metrics[c("BER","ACC","AUC","CID
X","SEN","SPE"),mNames]),cp$cpuElapsedTimes)))
colnames(classRanks) <- c("BER","ACC","AUC","CIDX","SEN","SPE","CPU")

classRanks$BER <- -classRanks$BER
classRanks$CPU <- -classRanks$CPU

colors_border = c( rgb(1.0,0.0,0.0,1.0), rgb(0.0,1.0,0.0,1.0) , rgb(0.0,0.0,1.0,1.0), rgb
(0.2,0.2,0.0,1.0), rgb(0.0,1.0,1.0,1.0), rgb(1.0,0.0,1.0,1.0), rgb(0.0,0.0,0.0,1.0) )
colors_in = c( rgb(1.0,0.0,0.0,0.05), rgb(0.0,1.0,0.0,0.05) , rgb(0.0,0.0,1.0,0.05),rgb(
1.0,1.0,0.0,0.05), rgb(0.0,1.0,1.0,0.05) , rgb(1.0,0.0,1.0,0.05), rgb(0.0,0.0,0.0,0.05) )
radarchart(classRanks,axistype = 0,maxmin = T,pcol = colors_border,pfcol = colors_in,plwd
= c(6,2,2,2,2,2,2),plty = 1, cglcol = "grey", cglty = 1,axislabcol = "black",cglwd = 0.8,
vlcex  = 0.5 ,title = "Prediction Model")

legend("topleft",legend = rownames(classRanks[-c(1,2),]),bty = "n",pch = 20,col = colors_
in,text.col = colors_border,cex = 0.5,pt.cex = 2)



filnames <- c("BSWiMS","LASSO","RF.ref","IDI","t-test","Kendall","mRMR")

filterRanks <- c(pr$minMaxMetrics$BER[1],pr$minMaxMetrics$ACC[2],pr$minMaxMetrics$AUC[2],
pr$minMaxMetrics$CIDX[2],pr$minMaxMetrics$SEN[2],pr$minMaxMetrics$SPE[2],max(cp$jaccard),
min(cp$featsize));

filterRanks <- rbind(filterRanks,c(pr$minMaxMetrics$BER[2],0,0,0,0,0,min(cp$jaccard),max
(cp$featsize)));

filterRanks <- as.data.frame(rbind(filterRanks,cbind(t(pr$metrics_filter[c("BER","ACC","A
UC","CIDX","SEN","SPE"),filnames]),cp$jaccard[filnames],cp$featsize[filnames])));
colnames(filterRanks) <- c("BER","ACC","AUC","CIDX","SEN","SPE","Jaccard","SIZE")
filterRanks$BER <- -filterRanks$BER
filterRanks$SIZE <- -filterRanks$SIZE

colors_border = c( rgb(1.0,0.0,0.0,1.0), rgb(0.0,1.0,0.0,1.0) , rgb(0.0,0.0,1.0,1.0), rgb
(0.2,0.2,0.0,1.0), rgb(0.0,1.0,1.0,1.0), rgb(1.0,0.0,1.0,1.0), rgb(0.0,0.0,0.0,1.0) )
```

```
colors_in = c( rgb(1.0,0.0,0.0,0.05), rgb(0.0,1.0,0.0,0.05) , rgb(0.0,0.0,1.0,0.05),rgb(
1.0,1.0,0.0,0.05), rgb(0.0,1.0,1.0,0.05) , rgb(1.0,0.0,1.0,0.05), rgb(0.0,0.0,0.0,0.05) )
radarchart(filterRanks,axistype = 0,maxmin = T,pcol = colors_border,pfcol = colors_in,plw
d = c(6,2,2,2,2,2,2,2),plty = 1, cglcol = "grey", cglty = 1,axislabcol = "black",cglwd = 0.
8, vlcex  = 0.6,title = "Filter Method" )


legend("topleft",legend = rownames(filterRanks[-c(1,2),]),bty = "n",pch = 20,col = colors
_in,text.col = colors_border,cex = 0.5,pt.cex = 2)
```

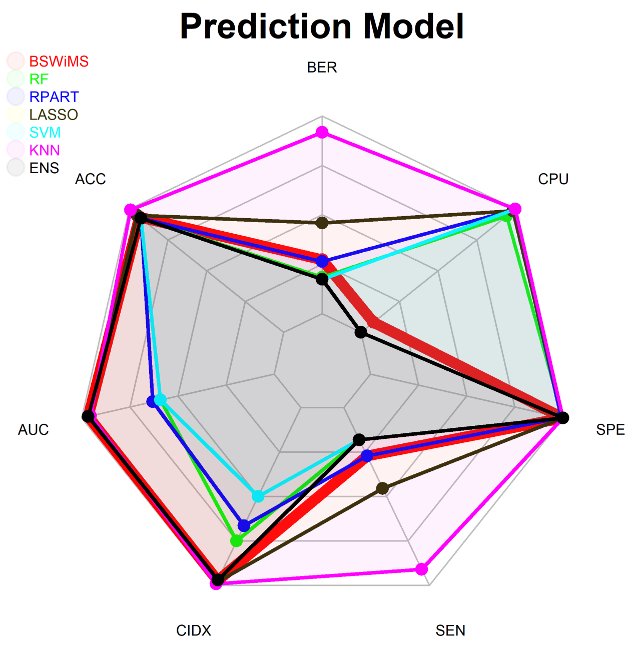


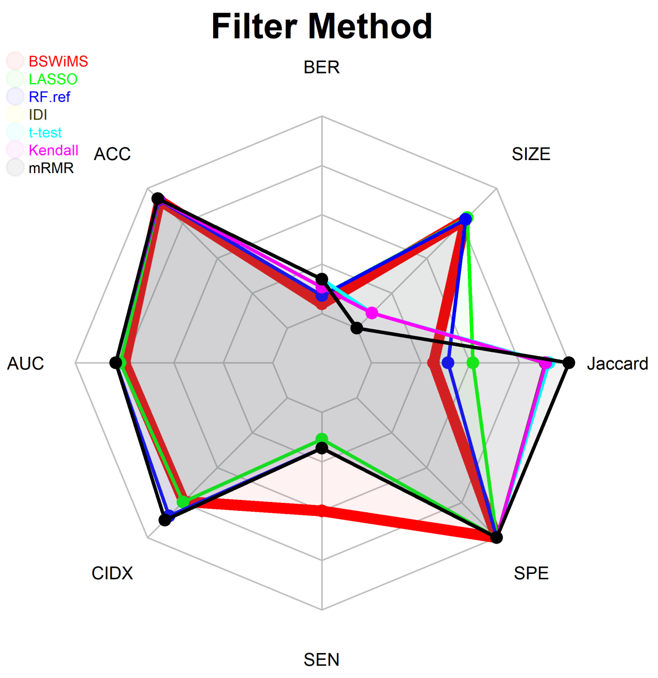


```
detach("package:fmsb", unload=TRUE)

par(mfrow = c(1,1))
par(op)
```

#### 1.3.3 Feature Analysis

```
rm <- rowMeans(cp$featureSelectionFrequency)
selFrequency <- cp$featureSelectionFrequency[rm > 0.25,]
gplots::heatmap.2(selFrequency,trace = "none",mar = c(10,10),main = "Features",cexRow =
0.25,cexCol = 0.5)
```

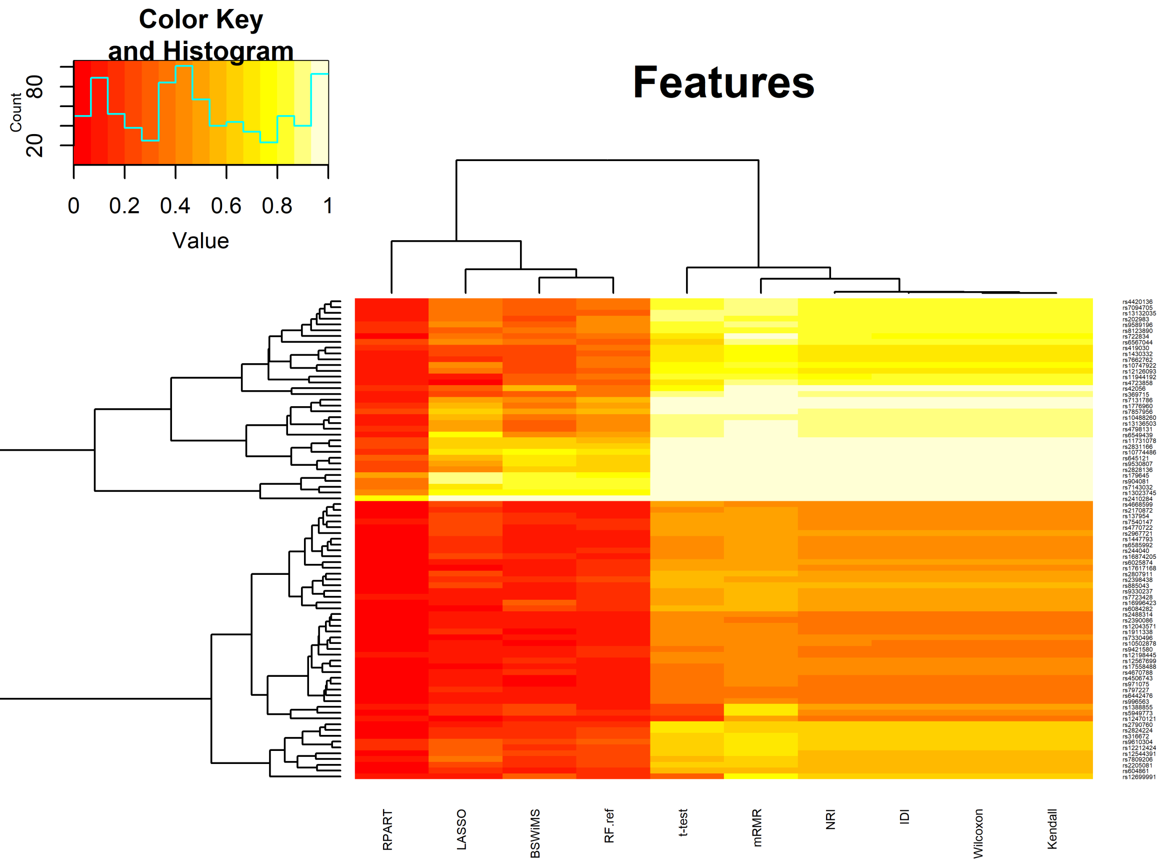


```
topFeat <- min(ncol(BSWiMSMODEL$bagging$formulaNetwork),30);
gplots::heatmap.2(log(100*BSWiMSMODEL$bagging$formulaNetwork[1:topFeat,1:topFeat] + 1),tr
ace="none",mar = c(10,10),main = "B:SWiMS Formula Network",cexRow = 0.5,cexCol = 0.5)
```

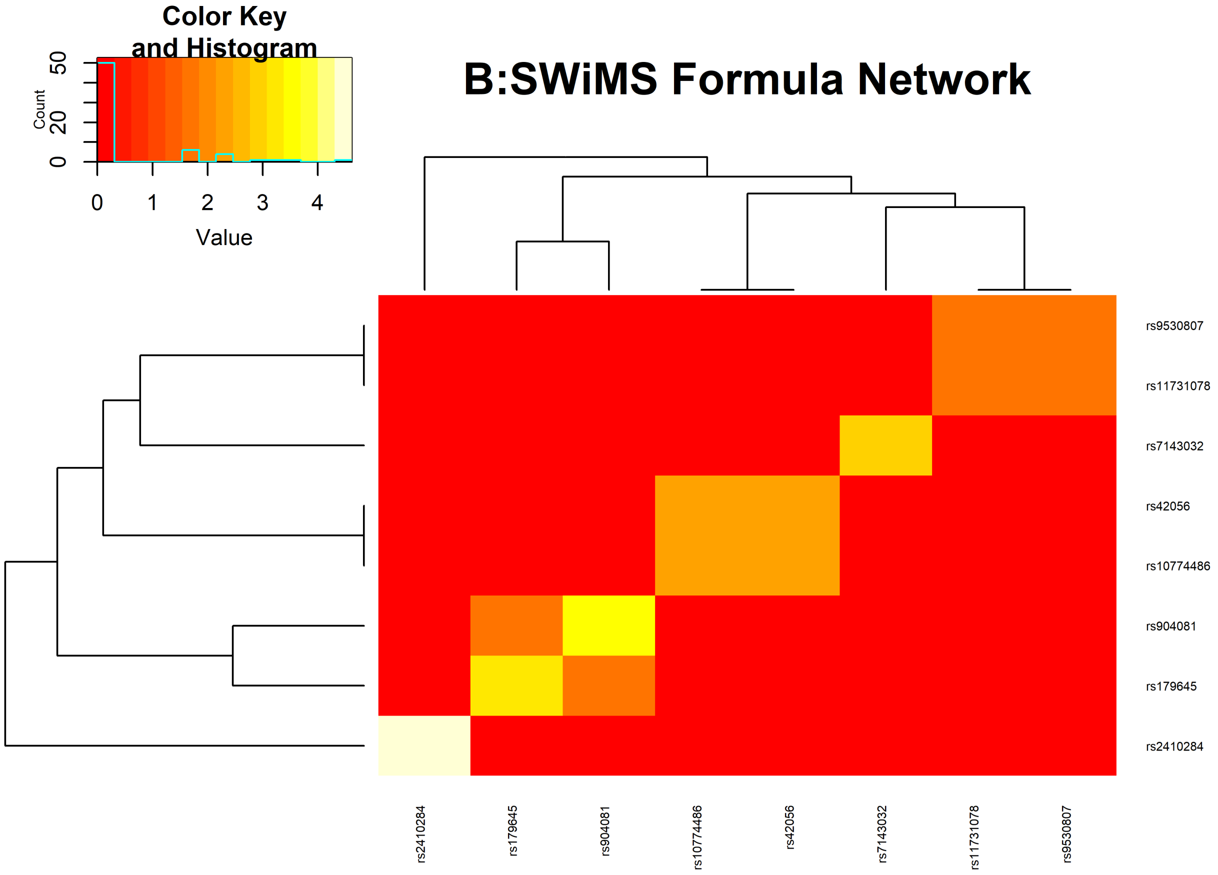


```
pander::pander(summary(BSWiMSMODEL$bagging$bagged.model,caption="Diabetes",round = 3))
```

- **coefficients**:

  Table continues below

| | Estimate | lower | OR | upper | u.Accuracy |
|---|---|---|---|---|---|
| **rs2410284** | 8.738 | 2143 | 6236 | 18147 | 0.7273 |
| **rs7143032** | -1.137 | 0.2735 | 0.3208 | 0.3762 | 0.6694 |
| **rs904081** | 4.371 | 41.91 | 79.13 | 149.4 | 0.6694 |
| **rs179645** | -3.969 | 0.01049 | 0.0189 | 0.03404 | 0.6694 |
| **rs10774486** | 5.492 | 105.1 | 242.8 | 561 | 0.6364 |
| **rs42056** | -5.444 | 0.001629 | 0.004324 | 0.01148 | 0.5785 |

  Table continues below

| | r.Accuracy | full.Accuracy | u.AUC | r.AUC | full.AUC |
|---|---|---|---|---|---|

| | r.Accuracy | full.Accuracy | u.AUC | r.AUC | full.AUC |
|---|---|---|---|---|---|
| **rs2410284** | 0.09091 | 0.7273 | 0.85 | 0.5 | 0.85 |
| **rs7143032** | 0.09091 | 0.6694 | 0.8182 | 0.5 | 0.8182 |
| **rs904081** | 0.1873 | 0.708 | 0.8182 | 0.553 | 0.8394 |
| **rs179645** | 0.2066 | 0.7157 | 0.8182 | 0.5636 | 0.8436 |
| **rs10774486** | 0.5785 | 0.8926 | 0.8 | 0.7682 | 0.9409 |
| **rs42056** | 0.6364 | 0.8926 | 0.7682 | 0.8 | 0.9409 |

| | IDI | NRI | z.IDI | z.NRI | Frequency |
|---|---|---|---|---|---|
| **rs2410284** | 0.5381 | 1.397 | 16.03 | 16.11 | 1 |
| **rs7143032** | 0.471 | 1.278 | 13.99 | 14.05 | 0.15 |
| **rs904081** | 0.4506 | 1.345 | 13.48 | 16.24 | 0.3 |
| **rs179645** | 0.4398 | 1.341 | 13.22 | 16.28 | 0.25 |
| **rs10774486** | 0.4298 | 1.771 | 12.85 | 29.98 | 0.1 |
| **rs42056** | 0.3529 | 1.708 | 10.93 | 28.55 | 0.1 |

- **Accuracy**: *0.9339*
- **tAUC**: *0.9636*
- **sensitivity**: *1*
- **specificity**: *0.9273*
- **bootstrap**:

```
sdata <- nearestNeighborImpute(Diabetes)[,c(theOutcome,rownames(selFrequency))]
```

…………

```
hm <- heatMaps(Outcome = theOutcome,data =sdata ,title = "Heat Map",theFiveColors=c("blu
e","cyan","Yellow","pink","red"),Scale = FALSE,hCluster = "col",cexRow = 0.25,cexCol = 0.
5,srtCol = 45)
```

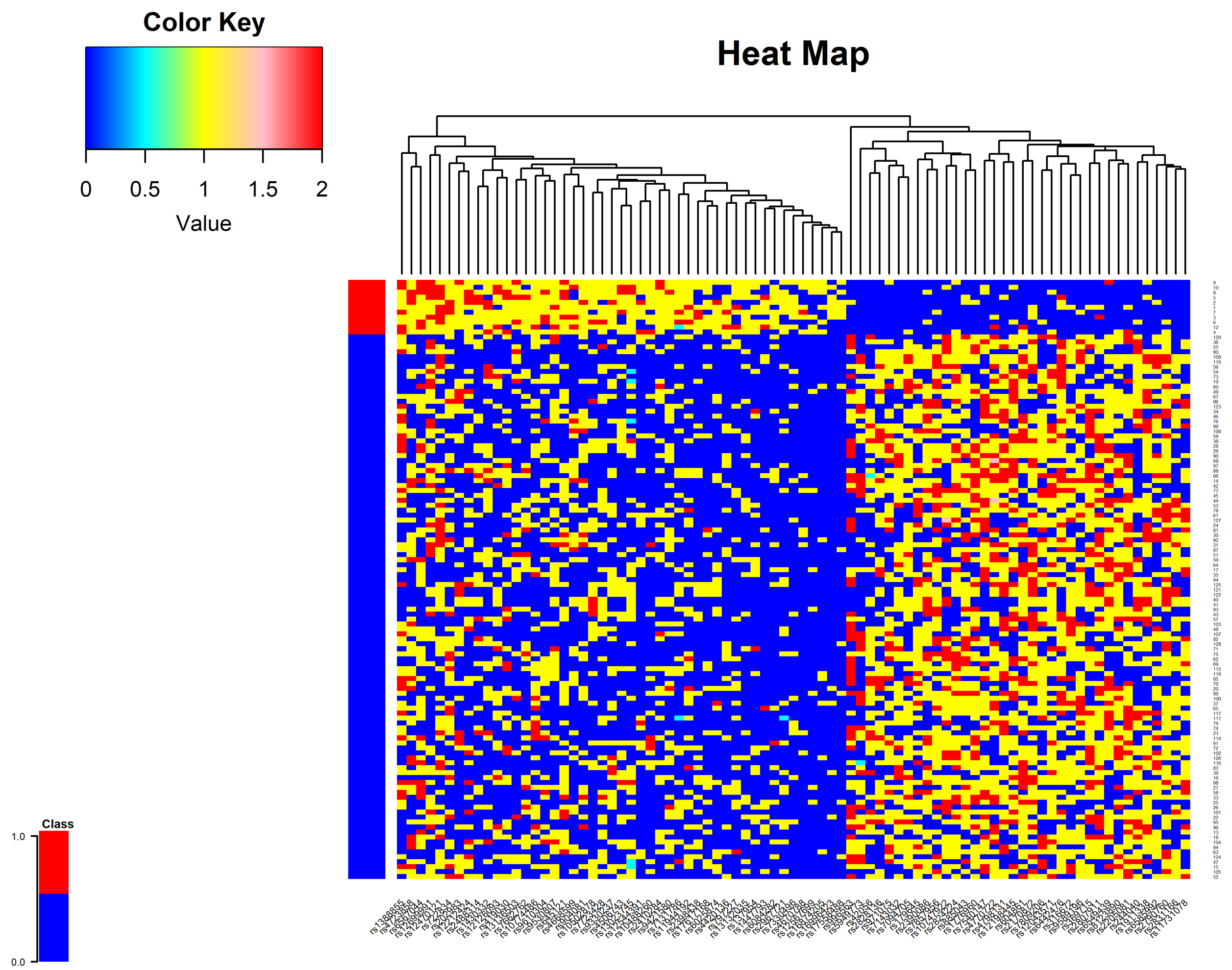


```
vlist <- rownames(selFrequency)
vlist <- cbind(vlist,vlist)
univ <- univariateRankVariables(variableList = vlist,formula = paste(theOutcome,"~1"),Out
come = theOutcome,data = theData,type = "LOGIT",rankingTest = "zIDI",uniType = "Binary")
[,c("controlMean","controlStd","caseMean","caseStd","ROCAUC","WilcoxRes.p")]

cnames <- colnames(univ);
univ <- cbind(univ,rm[rownames(univ)])
colnames(univ) <- c(cnames,"Frequency")
univ <- univ[order(-univ[,5]),]
pander::pander(univ,caption = "Features",round = 4)
```

Features (continued below)

| | controlMean | controlStd | caseMean | caseStd | ROCAUC |
|---|---|---|---|---|---|
| **rs2410284** | 77 | 33 | 11 | NA | 0.85 |
| **rs7131786** | 81 | 29 | 1 | 10 | 0.8227 |

| | controlMean | controlStd | caseMean | caseStd | ROCAUC |
|---|---|---|---|---|---|
| **rs7857956** | 30 | 80 | 10 | 1 | 0.8182 |
| **rs6549439** | 30 | 80 | 10 | 1 | 0.8182 |
| **rs904081** | 70 | 40 | 11 | NA | 0.8182 |
| **rs179645** | 40 | 70 | 11 | NA | 0.8182 |
| **rs7143032** | 40 | 70 | 11 | NA | 0.8182 |
| **rs10488260** | 89 | 21 | 2 | 9 | 0.8136 |
| **rs13132035** | 88 | 22 | 2 | 9 | 0.8091 |
| **rs13136503** | 78 | 32 | 1 | 10 | 0.8091 |
| **rs1776960** | 32 | 78 | 10 | 1 | 0.8091 |
| **rs13023745** | 0.4045 | 0.486 | 1 | 0 | 0.8045 |
| **rs369715** | 33 | 77 | 10 | 1 | 0.8045 |
| **rs4420136** | 87 | 23 | 2 | 9 | 0.8045 |
| **rs10774486** | 66 | 44 | 11 | NA | 0.8 |
| **rs7094705** | 34 | 76 | 10 | 1 | 0.8 |
| **rs8123890** | 34 | 76 | 10 | 1 | 0.8 |
| **rs9589196** | 34 | 76 | 10 | 1 | 0.8 |
| **rs4798131** | 34 | 76 | 10 | 1 | 0.8 |
| **rs202983** | 75 | 35 | 1 | 10 | 0.7955 |
| **rs11944192** | 75 | 35 | 1 | 10 | 0.7955 |
| **rs2790760** | 25 | 85 | 9 | 2 | 0.7955 |
| **rs2824224** | 26 | 84 | 9 | 2 | 0.7909 |
| **rs2831166** | 46 | 64 | 11 | NA | 0.7909 |
| **rs11731078** | 46 | 64 | 11 | NA | 0.7909 |
| **rs2828136** | 47 | 63 | 11 | NA | 0.7864 |
| **rs9530807** | 63 | 47 | 11 | NA | 0.7864 |
| **rs645121** | 63 | 47 | 11 | NA | 0.7864 |
| **rs419030** | 73 | 37 | 1 | 10 | 0.7864 |
| **rs2205081** | 27 | 83 | 9 | 2 | 0.7864 |
| **rs6084282** | 93 | 17 | 3 | 8 | 0.7864 |

| | controlMean | controlStd | caseMean | caseStd | ROCAUC |
|---|---|---|---|---|---|
| **rs12126093** | 72 | 38 | 1 | 10 | 0.7818 |
| **rs7662762** | 72 | 38 | 1 | 10 | 0.7818 |
| **rs10747922** | 38 | 72 | 10 | 1 | 0.7818 |
| **rs2807911** | 28 | 82 | 9 | 2 | 0.7818 |
| **rs604861** | 28 | 82 | 9 | 2 | 0.7818 |
| **rs12544391** | 92 | 18 | 3 | 8 | 0.7818 |
| **rs1430332** | 71 | 39 | 1 | 10 | 0.7773 |
| **rs9610304** | 71 | 39 | 1 | 10 | 0.7773 |
| **rs316672** | 39 | 71 | 10 | 1 | 0.7773 |
| **rs885043** | 30 | 80 | 9 | 2 | 0.7727 |
| **rs2967721** | 90 | 20 | 3 | 8 | 0.7727 |
| **rs7809206** | 40 | 70 | 10 | 1 | 0.7727 |
| **rs9330237** | 70 | 40 | 1 | 10 | 0.7727 |
| **rs12212424** | 70 | 40 | 1 | 10 | 0.7727 |
| **rs4770722** | 31 | 79 | 9 | 2 | 0.7682 |
| **rs42056** | 51 | 59 | 11 | NA | 0.7682 |
| **rs17617168** | 89 | 21 | 3 | 8 | 0.7682 |
| **rs137954** | 89 | 21 | 3 | 8 | 0.7682 |
| **rs7723428** | 68 | 42 | 1 | 10 | 0.7636 |
| **rs12198445** | 32 | 78 | 9 | 2 | 0.7636 |
| **rs2390086** | 32 | 78 | 9 | 2 | 0.7636 |
| **rs12043571** | 32 | 78 | 9 | 2 | 0.7636 |
| **rs1911338** | 32 | 78 | 9 | 2 | 0.7636 |
| **rs2398438** | 88 | 22 | 3 | 8 | 0.7636 |
| **rs1447793** | 98 | 12 | 4 | 7 | 0.7636 |
| **rs6585992** | 43 | 67 | 10 | 1 | 0.7591 |
| **rs971075** | 43 | 67 | 10 | 1 | 0.7591 |
| **rs244040** | 43 | 67 | 10 | 1 | 0.7591 |
| **rs9421580** | 67 | 43 | 1 | 10 | 0.7591 |

| | controlMean | controlStd | caseMean | caseStd | ROCAUC |
|---|---|---|---|---|---|
| **rs2488314** | 77 | 33 | 2 | 9 | 0.7591 |
| **rs4668599** | 77 | 33 | 2 | 9 | 0.7591 |
| **rs4506743** | 77 | 33 | 2 | 9 | 0.7591 |
| **rs7540147** | 33 | 77 | 9 | 2 | 0.7591 |
| **rs10502878** | 77 | 33 | 2 | 9 | 0.7591 |
| **rs2170872** | 23 | 87 | 8 | 3 | 0.7591 |
| **rs6442476** | 23 | 87 | 8 | 3 | 0.7591 |
| **rs797227** | 87 | 23 | 3 | 8 | 0.7591 |
| **rs6025874** | 87 | 23 | 3 | 8 | 0.7591 |
| **rs4723858** | 57 | 53 | 11 | NA | 0.7591 |
| **rs722834** | 57 | 53 | 11 | NA | 0.7591 |
| **rs7330496** | 97 | 13 | 4 | 7 | 0.7591 |
| **rs6567044** | 56 | 54 | 11 | NA | 0.7545 |
| **rs16996423** | 106 | 4 | 5 | 6 | 0.7545 |
| **rs996563** | 45 | 65 | 10 | 1 | 0.75 |
| **rs17558488** | 104 | 6 | 5 | 6 | 0.7455 |
| **rs12699991** | 53 | 57 | 11 | NA | 0.7409 |
| **rs12567699** | 103 | 7 | 5 | 6 | 0.7409 |
| **rs4670788** | 103 | 7 | 5 | 6 | 0.7409 |
| **rs16874205** | 103 | 7 | 5 | 6 | 0.7409 |
| **rs1388855** | 52 | 58 | 11 | NA | 0.7364 |
| **rs5949773** | 59 | 51 | 11 | NA | 0.7318 |
| **rs12470121** | 49 | 61 | 11 | NA | 0.7227 |

| | WilcoxRes.p | Frequency |
|---|---|---|
| **rs2410284** | 0.1572 | 0.964 |
| **rs7131786** | 0.0479 | 0.746 |
| **rs7857956** | 0.0654 | 0.754 |
| **rs6549439** | 0.0688 | 0.732 |

| | WilcoxRes.p | Frequency |
|---|---|---|
| **rs904081** | 0.7038 | 0.898 |
| **rs179645** | 0.6765 | 0.904 |
| **rs7143032** | 0.6957 | 0.872 |
| **rs10488260** | 2e-04 | 0.68 |
| **rs13132035** | 8e-04 | 0.622 |
| **rs13136503** | 0.1641 | 0.675 |
| **rs1776960** | 0.1427 | 0.728 |
| **rs13023745** | 0.8428 | 0.868 |
| **rs369715** | 0.2106 | 0.655 |
| **rs4420136** | 0.0014 | 0.619 |
| **rs10774486** | 0.898 | 0.829 |
| **rs7094705** | 0.2682 | 0.613 |
| **rs8123890** | 0.2962 | 0.631 |
| **rs9589196** | 0.2742 | 0.645 |
| **rs4798131** | 0.2751 | 0.682 |
| **rs202983** | 0.3385 | 0.615 |
| **rs11944192** | 0.3577 | 0.576 |
| **rs2790760** | 0.0077 | 0.446 |
| **rs2824224** | 0.0167 | 0.43 |
| **rs2831166** | 0.9541 | 0.807 |
| **rs11731078** | 0.9384 | 0.812 |
| **rs2828136** | 0.9661 | 0.784 |
| **rs9530807** | 0.9641 | 0.788 |
| **rs645121** | 0.9738 | 0.799 |
| **rs419030** | 0.5186 | 0.531 |
| **rs2205081** | 0.0285 | 0.421 |
| **rs6084282** | 0 | 0.368 |
| **rs12126093** | 0.5961 | 0.542 |
| **rs7662762** | 0.5576 | 0.519 |

| | WilcoxRes.p | Frequency |
|---|---|---|
| **rs10747922** | 0.594 | 0.577 |
| **rs2807911** | 0.0409 | 0.369 |
| **rs604861** | 0.0367 | 0.39 |
| **rs12544391** | 0 | 0.457 |
| **rs1430332** | 0.6718 | 0.514 |
| **rs9610304** | 0.6802 | 0.486 |
| **rs316672** | 0.6866 | 0.458 |
| **rs885043** | 0.0931 | 0.393 |
| **rs2967721** | 3e-04 | 0.342 |
| **rs7809206** | 0.7583 | 0.431 |
| **rs9330237** | 0.7436 | 0.361 |
| **rs12212424** | 0.7469 | 0.485 |
| **rs4770722** | 0.1381 | 0.33 |
| **rs42056** | 0 | 0.691 |
| **rs17617168** | 6e-04 | 0.307 |
| **rs137954** | 6e-04 | 0.328 |
| **rs7723428** | 0.8447 | 0.351 |
| **rs12198445** | 0.1981 | 0.29 |
| **rs2390086** | 0.2003 | 0.269 |
| **rs12043571** | 0.2098 | 0.267 |
| **rs1911338** | 0.184 | 0.279 |
| **rs2398438** | 0.0012 | 0.369 |
| **rs1447793** | 0 | 0.319 |
| **rs6585992** | 0.8793 | 0.319 |
| **rs971075** | 0.885 | 0.252 |
| **rs244040** | 0.8899 | 0.31 |
| **rs9421580** | 0.8884 | 0.272 |
| **rs2488314** | 0.2606 | 0.281 |
| **rs4668599** | 0.2548 | 0.313 |

| | WilcoxRes.p | Frequency |
|---|---|---|
| **rs4506743** | 0.2803 | 0.259 |
| **rs7540147** | 0.2648 | 0.335 |
| **rs10502878** | 0.2606 | 0.269 |
| **rs2170872** | 0.0029 | 0.31 |
| **rs6442476** | 0.0028 | 0.253 |
| **rs797227** | 0.0031 | 0.253 |
| **rs6025874** | 0.0023 | 0.333 |
| **rs4723858** | 0 | 0.562 |
| **rs722834** | 0 | 0.595 |
| **rs7330496** | 0 | 0.285 |
| **rs6567044** | 0 | 0.608 |
| **rs16996423** | 0 | 0.37 |
| **rs996563** | 0.9394 | 0.255 |
| **rs17558488** | 0 | 0.28 |
| **rs12699991** | 0 | 0.424 |
| **rs12567699** | 0 | 0.285 |
| **rs4670788** | 0 | 0.287 |
| **rs16874205** | 0 | 0.32 |
| **rs1388855** | 0 | 0.342 |
| **rs5949773** | 0 | 0.323 |
| **rs12470121** | 1 | 0.253 |
